# Robust coexistence in competitive ecological communities

**DOI:** 10.1101/2024.07.10.602979

**Authors:** Pablo Lechón-Alonso, Srilena Kundu, Paula Lemos-Costa, José A. Capitan, Stefano Allesina

## Abstract

Darwin already recognized that competition is fiercest among conspecifics, a principle that later made intraspecific competition central to ecological theory through concepts such as niche differentiation and limiting similarity. Beyond shaping coexistence, strong intraspecific competition can also stabilize community dynamics by ensuring that populations return to equilibrium after disturbance. Here we investigate a more fundamental question: how intraspecific competition influences the very existence of a steady state (feasibility) in large random ecological communities dominated by competition. We show that, analogous to classical results on stability, there is a critical level of intraspecific competition above which a feasible steady state is guaranteed to exist. We derive a general expression for the probability of feasibility and prove that, asymptotically (as species number grows), the transition to stability occurs before the transition to feasibility with probability one. Thus, in large competitive communities, any feasible equilibrium is automatically stable. This ordering persists even when many species in the initial pool cannot coexist and extinctions occur: the dynamics prune the community, shifting feasibility and stability thresholds but never reversing their order. These results imply that large competitive communities generically converge to a globally stable equilibrium, making sustained oscillations or chaos unlikely—consistent with experimental observations.

## Introduction

The notions of intraspecific competition and population self-regulation play a central role in ecological theory, leading to foundational concepts such as limiting similarity and niche differentiation[1–6]. They also are crucial for coexistence: for example, May[7] showed that ecological dynamics can be stabilized by imposing sufficiently strong intraspecific competition, and that for large systems with random species interactions the stability transition is universal (independent of the probability distribution used to model interactions) and sharp (always achieved at the same critical value of intraspecific competition). Importantly, the existence of an equilibrium (feasibility) is a necessary condition for coexistence. Further, results on stability can dramatically change when accounting for feasibility. The first contribution to this problem can be traced back to a response to May’s work on stability[8], showing that in competitive Lotka-Volterra models with random interactions the probability of stability, conditioned on the presence of a feasible equilibrium, does increase, rather than decrease with system size. Since then, significant advances have been gradually accumulating in the literature[9–15] (Supplementary Note 3, Sec. 2). However, the progress on the study of feasibility has yet to match that reached for stability[7, 16–18]. Here we show that accounting for feasibility is key to determining the expected dynamics of large competitive systems.

We compute the probability that a system would display a feasible equilibrium given its level of intraspecific competition, size, and the statistics on the random distribution of species interactions. We show that the transition to feasibility is universal, as for stability, but is not sharp—thereby extending, generalizing and refining previous results[8, 9, 12, 14, 19–22]. More importantly, we show that, in large random competitive systems, when we increase the strength of intraspecific competition, the transition to stability precedes that to feasibility[8, 9, 20]. Therefore, in these large systems feasibility implies stability. This result holds even when the initial pool of species cannot coexist, and extinctions ensue. That is, during ecological dynamics, species extinctions will alter the values for the stability and feasibility thresholds, but not their ordering. Differently from May, who focused on local asymptotic stability of an equilibrium, here we consider the stronger criterion of global stability—robust coexistence in which populations rebound from any perturbation short of extinction.

Overall, these findings have far-reaching implications for the ecological dynamics of large random competitive ecosystems. They rule out coexistence through limit cycles or chaos[23, 24], and instead indicate that the community dynamics converge to a highly-robust equilibrium, such that (1) any perturbation short of extinction would be buffered, (2) the community resists invasions, and (3) it can be assembled from the ground up[25, 26].

## Results

### Stability and feasibility thresholds

We consider the Generalized Lotka-Volterra[27–30] (GLV) model for *n* interacting populations:

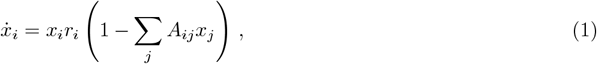

where intrinsic growth rates *r*_*i*_ are organized in the vector *r*, and interactions *A*_*ij*_ in the matrix *A*. We say that Eq. (1) has a feasible (i.e., biologically attainable) equilibrium whenever there is a **x**^⋆^ > **0** such that 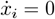 for all populations, and that such a feasible equilibrium is asymptotically stable if dynamics return to it following a perturbation.

Following May[7], we model *A* as the sum of two matrices, *A* = *αI* + *B*, where *B* collects the interspecific interactions (*B*_*ij*_, *i* ≠ *j*). *I* is the identity matrix, and *α* ≥ 0 the intraspecific competition in the system, assumed to be the same for all species (relaxing this assumption does not change the results, see Supplementary Note 3, Sec. 9)

Now we consider increasing *α* from zero to an arbitrary large positive value, and track the effects of this change on coexistence. We denote *α*_*S*_ as the level of intraspecific competition that is sufficient to guarantee the (global) stability of an equilibrium (Methods, and Supplementary Note 1, Sec. 3). Importantly, for any *α* ≥ *α*_*S*_ stability is maintained, i.e., once surpassed this threshold, dynamics always converge to a globally stable equilibrium. Here we ask whether, in analogy to stability, there exist a critical level *α*_*F*_ that a) guarantees the existence of a positive equilibrium, and b) further increasing *α* ≥ *α*_*F*_ retains the feasibility of the equilibrium. In particular, we want to probe the relationship between *α*_*F*_ and *α*_*S*_.

We start with a toy example (Fig. 1, *n* = 6, see Supplementary Note 3 and Supplementary Figure 3 for an example with two populations), illustrating the effect of increasing *α* on feasibility and stability. We first sample a baseline interaction matrix *B*, and compute the thresholds for stability, *α*_*S*_, and feasibility, *α*_*F*_ (Methods, and Supplementary Note 1, Sec. 3, and Supplementary Note 2, Sec. 4), marked in Fig. 1 with green and yellow triangles, respectively. We then construct three interaction matrices *A* that differ only in their intraspecific competition values, *α*_*I*_, *α*_*II*_, and *α*_*III*_ (Fig. 1). For each system, we integrate the dynamics in Eq. (1) from two different initial conditions (Fig. 1 bottom panels, solid vs. dashed lines), revealing three qualitatively distinct dynamical regimes. System I (Fig. 1, left) undergoes extinctions and displays alternative stable states, indicating that the coexistence equilibrium is not feasible, and the resulting sub-community equilibrium is not globally stable (indeed, neither stability nor feasibility thresholds have been surpassed, since *α*_*I*_ <*α*_*S*_ <*α*_*F*_). System II (Fig. 1, middle), also displays extinctions (as there is no feasible equilibrium), but starting from different initial conditions no longer leads to different outcomes, because the equilibrium of the sub-community is globally stable (*α*_*S*_ <*α*_II_ <*α*_*F*_, and see Supplementary Note 1, Sec. 3). Finally, system III (Fig. 1, right), exhibits all populations coexisting stably (*α*_*S*_ <*α*_*F*_ <*α*_III_). From this single realization, we can see that stability is achieved before feasibility. Next, we show that this is the case for large random ecological communities (but see Supplementary Note 2, Sec. 3 for results on small or arbitrary competitive systems).

**Figure 1.**
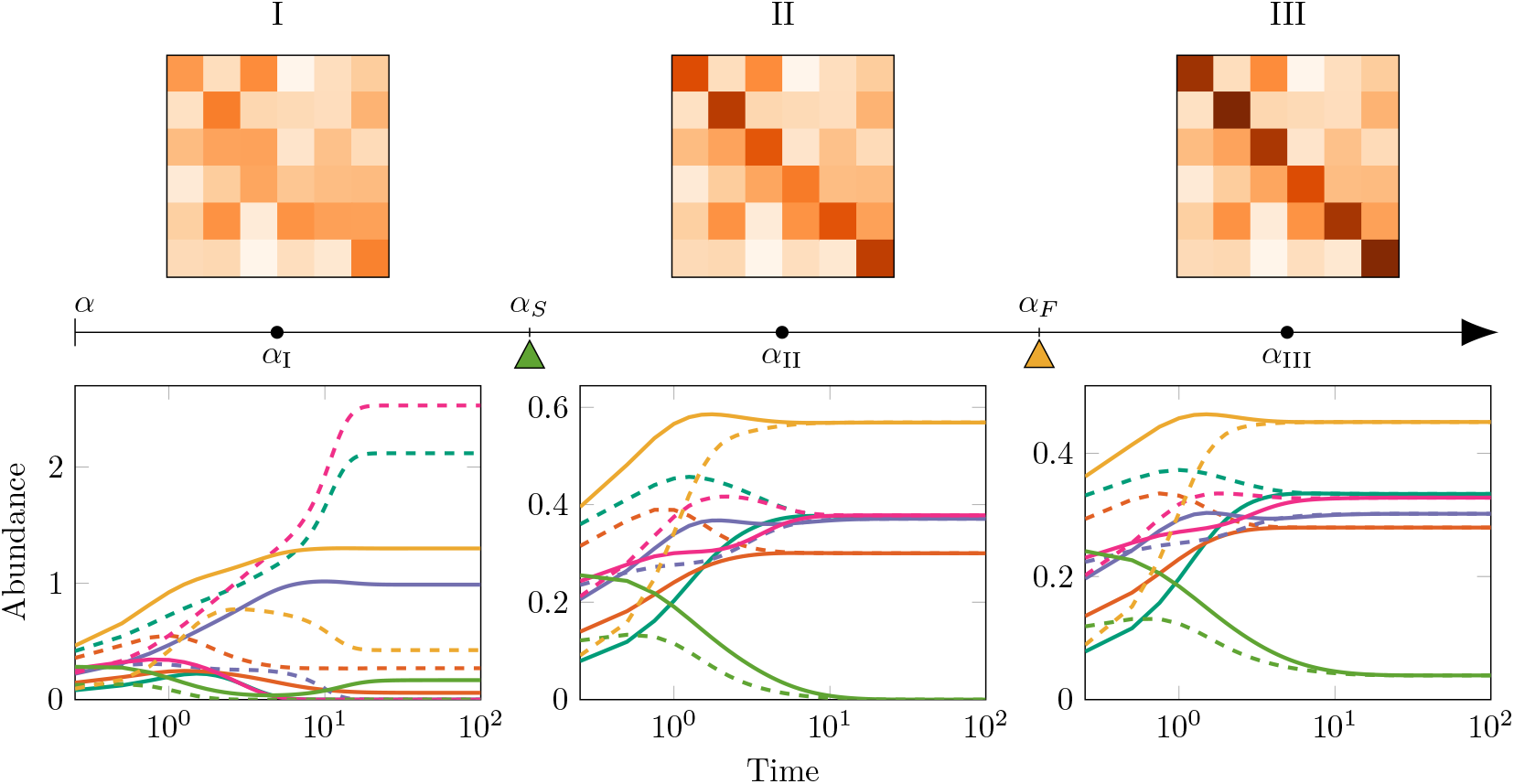
Three dynamical scenarios arising in an example random competitive community with six species (see Supplementary Figure 3 for an example with two populations). We consider the dynamics for a GLV model with *A* = *αI* + *B* (Eq.1), with *B* being the same across the three systems, and intraspecific competition *α* taking three different values (*α*_I_, *α*_II_, and *α*_III_). The three matrices, with the same off-diagonal coefficients and increasingly strong diagonal coefficients, are shown as tile plots above the arrow (darker colors reflect stronger competition). By increasing *α* from *α*_I_ to *α*_III_, we observe distinct dynamical behaviors emerging as *α* surpasses the stability (*α*_*S*_) and feasibility (*α*_*F*_) thresholds. First, when *α*_I_ <*α*_*S*_ <*α*_*F*_, initializing the population densities (colors) at different values (linetypes) leads to extinctions, and a different set of coexisting populations (priority effects or alternative stable states): the equilibrium is not feasible and the set of surviving species depends on the initial conditions. Second, when *α*_*S*_ <*α*_II_ <*α*_*F*_, the equilibrium is not feasible, but the same populations go extinct irrespective of initial conditions (in this case, the green population), while the remaining populations converge to a stable equilibrium (Supplementary Note 1, Sec. 3). Third, when *α*_*S*_ <*α*_*F*_ <*α*_III_, a feasible equilibrium exists, and is globally stable.

### Feasibility implies robust coexistence almost surely

For each unstable matrix *B*, we can always find a critical level *α*_*S*_ such that for any *α < α*_*S*_ the matrix *A* = *αI* + *B* is unstable, and for any value *α* ≥ *α*_*S*_ it is stable (Supplementary Note 1, Sec. 3). Tracking the existence of an equilibrium as *α* increases is more complicated: for feasibility, we need (*αI* +*B*)**x** = **1** to have a positive solution. As shown in Fig. 2a, when increasing *α*, a positive solution can appear or disappear several times; however, we prove that, for any *B*, a critical value *α*_*F*_ ensures a) that (*α*_*F*_ *I* + *B*)**x** = **1** has a positive solution, and b) that further increasing *α > α*_*F*_ maintains the positivity of the solution (Supplementary Note 2, Sec. 3).

**Figure 2.**
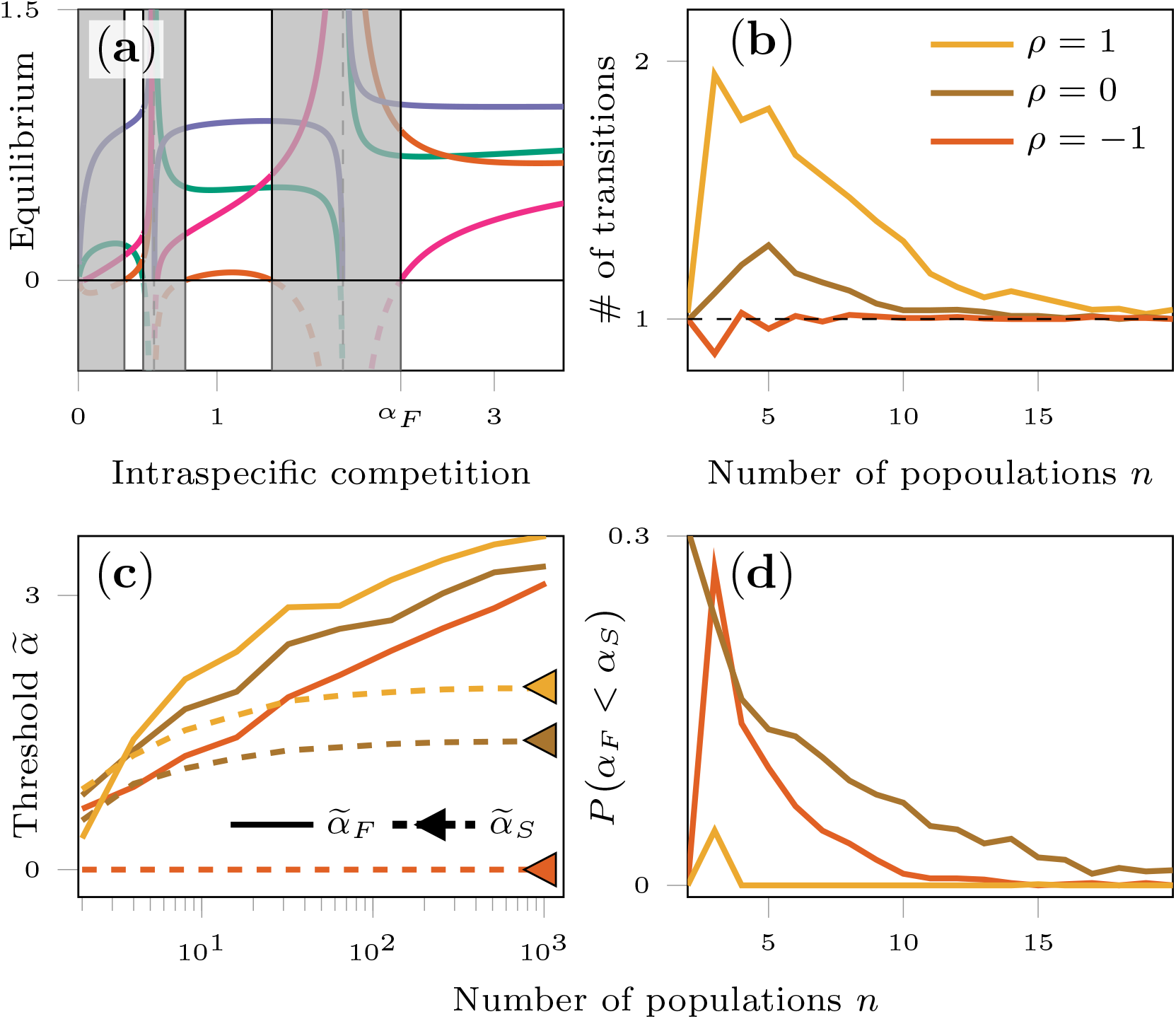
While low-dimensional systems may gain and lose feasibility multiple times, high-dimensional systems exhibit a single feasibility transition that almost always follows the stability transition. **(a)** Bifurcation diagram of equilibrium abundances as a function of intraspecific competition *α*. Shaded regions correspond to parameter values where no feasible equilibrium exists, while unshaded regions mark feasibility. For a system with four populations (colors), we see that when *α* = 0 no feasible solution of (*αI* + *B*)*x* = **1** exists; as *α* increases, feasible equilibria can appear and disappear multiple times, but for sufficiently large *α* ≥ *α*_*F*_ feasibility is always guaranteed (Supplementary Note 2, Sec. 3). **(b)–(d)** Results from simulations across many system sizes. For each *n*, we sample many random interaction matrices with entries drawn in correlated pairs from a bivariate normal distribution with mean 0, unit variance, and correlation *ρ* (colors), and calculate both feasibility and stability thresholds. **(b)** The average number of transitions in and out of feasibility (y-axis) decreases with *n* and converges to one, indicating that for large systems feasibility is excluded below the feasibility threshold *α* and guaranteed above it. **(c)** Normalized thresholds 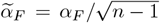 and 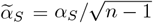 as a function of *n*. While 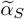 converges to a constant, 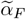 grows without bound. Triangles mark 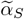 for *n* = 10, also shown in Fig. 3. **(d)** Probability that feasibility comes before stability *P* (*α*_*F*_ <*α*_*S*_) as a function of *n*, i.e. the chance that a feasible but unstable equilibrium exists. This probability decreases rapidly with system size.

Moreover, Fig. 2b shows that, as the size of the community increases, the number of transitions in and out of feasibility converges to one; asymptotically (i.e., as *n* → ∞), *α*_*F*_ behaves exactly as *α*_*S*_, with feasibility precluded below *α*_*F*_ and guaranteed above it. Simulations can also give us a qualitative understanding of the behavior of *α*_F_ and *α*_*S*_ as the number of populations in the system increases: Fig. 2c shows that, once properly normalized to account for matrix size 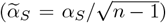, the threshold for stability converges to a constant, while that for feasibility 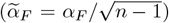 keeps increasing with no saturation. Consequently (Fig. 2d), asymptotically we have *α*_*F*_ *> α*_*S*_. Next, we show how these results can be derived mathematically.

In order to mathematically formalize the patterns from the toy example and auxiliary simulations (Figs. 1 and 2), we focus on the ensemble of large random ecosystems where each pair of interactions (*B*_*ij*_, *B*_*ji*_), with *i*≠ *j* is sampled from any bivariate distribution with mean *µ >* 0, variance *σ*^2^, and correlation *ρ* (Supplementary Note 3, Sec. 1); and the diagonal elements *B*_*ii*_ = 0. We call these systems competitive whenever **r** > **0** (i.e., all populations can grow in isolation), and ∑_*j*_ *B*_*ij*_ > 0 for every population *i* (i.e., each population has, on average, a negative effect on the growth of the other populations). We compute the probability *p*_*F*_ that one such random system of size *n* will have a feasible equilibrium as a function of *α*.

We begin by noting two simplifications that can be made without loss of generality (Methods). First, feasibility is unchanged if we subtract from each column of *B* its mean, obtaining a matrix *C* with zero column sums. This means that the absolute strength of competition is irrelevant: increasing the competitive effect of a given population on all others by the same amount does not alter whether the equilibrium is feasible.

Thus, we may restrict attention to the canonical system

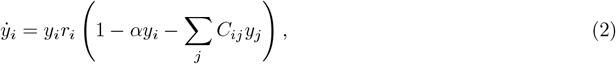

with *C* centered in this way. Second, varying *α* is equivalent to rescaling the variance *σ*^2^ of the distribution of species interactions, so we may fix *σ*^2^ = 1 for convenience. Together, these reductions yield a simplified yet fully general representation of the competitive system, which we adopt for the analysis below.

For any model in which populations are statistically equivalent (i.e., there is no a priori structure making some populations behave differently from the rest), the distribution of the equilibrium values, **y**^⋆^, has always the same mean and correlation structure (Methods). Moreover, when *n* is large, we can assume the components of **y**^⋆^ to be approximately normally distributed[9] and uncorrelated (Methods, and Supplementary Note 3, Sec. 7). Then, the probability of feasibility can be taken as the product of the probabilities that each component is positive, and can thus be written as the cumulative distribution function of a standard normal distribution, raised to the *n*^th^ power (Supplementary Note 3, Sec. 7):

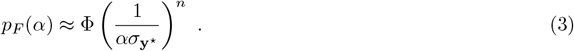

As long as populations are statistically equivalent, more complex models, e.g., with higher-order correlations or network structure, will follow the same expression, but with a different *σ*_***y***_⋆. Computing the variance 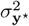 (which in general depends on *n, α*, and *ρ*) is sufficient to approximate the probability of feasibility. In our case, we use the theory of resolvents[31] (see Supplementary Notes 3.8 and 3.10 for an alternative method based on iterative solutions) to approximate the variance for any *ρ*, obtaining:

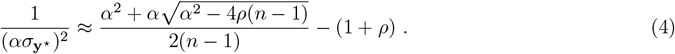

Substituting this expression into Eq. (3), we obtain an extremely accurate analytical approximation of the probability of feasibility (Fig. 3, black curves): the probability *p*_*F*_ (*α*) is vanishingly small when *α* = 0 (Supplementary Note 3, Sec. 3), and increases smoothly to 1 with increasing *α*. Eq. (4) shows that the details of the distribution of interactions (Fig. 3, linetypes), once accounted for the correlation (Fig. 3, colors), are irrelevant when the system is large. This means that asymptotically, the curve describing the probability of feasibility is universal. Interestingly, exactly as a negative correlation between interaction strengths improves the chance of stability[16], it also increases the probability of feasibility[21]. Contrary to current expectations[14], the effect of the pairwise correlation *ρ* on the probability of feasibility remains significant for large systems (*n* = 1000 in Fig. 3). Additionally, the transition to feasibility also remains smooth compared to stability (Fig. 3, and Supplementary Fig. 10, see SupplementaryNote 3, Sec. 8 for a detailed discussion) for large systems.

**Figure 3.**
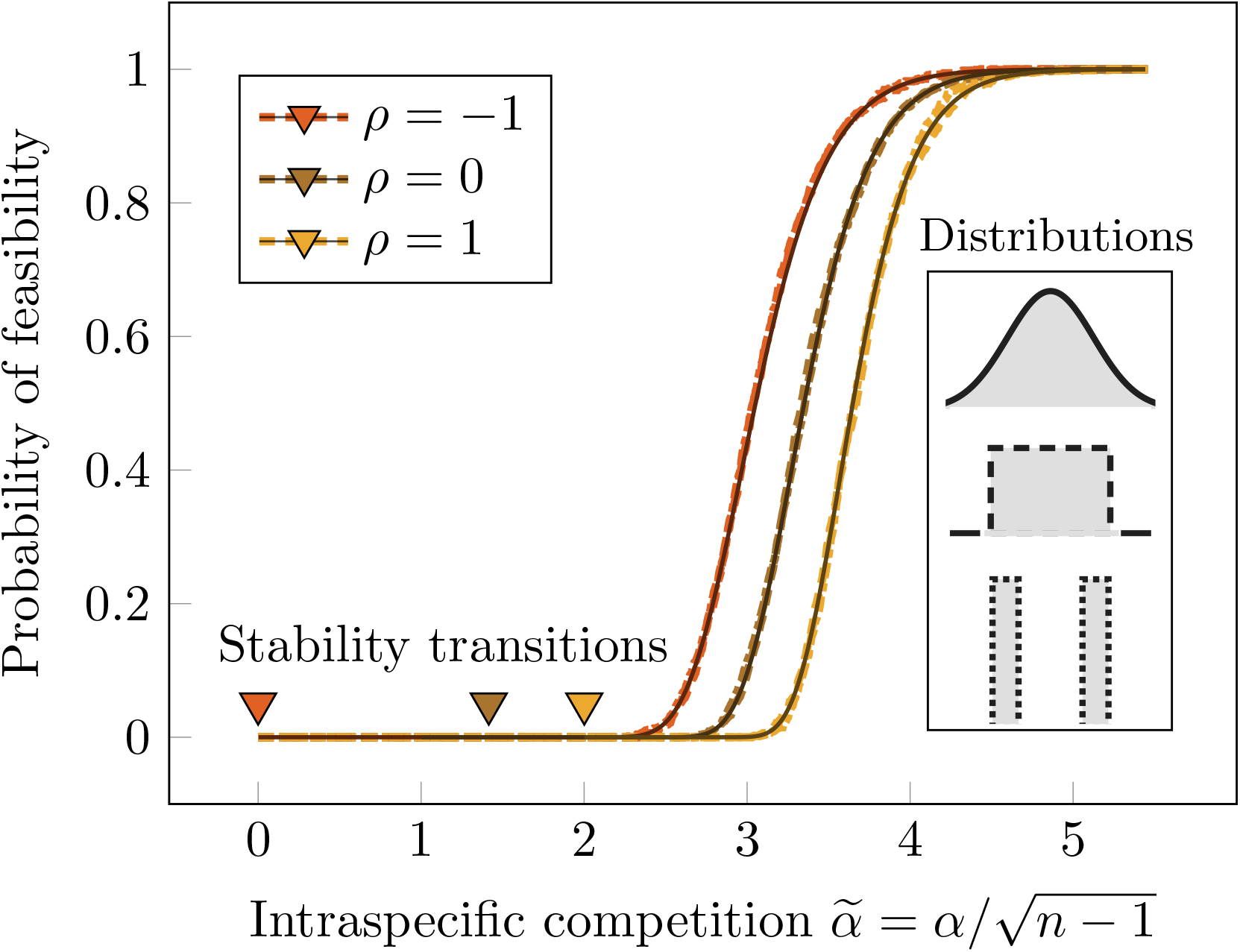
In large systems, feasible equilibria are stable with high probability. Probability of feasibility for a given level of intraspecific competition (we plot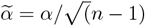, x-axis) for systems of size *n* = 1000. We sample entries from distributions (line types) whose marginals are either normal, uniform, or “discrete” (coefficients take positive or negative values with equal probability, see Supplementary Note 4). The curves for different distributions collapse to the same curve, illustrating that results are asymptotically universal (i.e., the higher moments of the distribution have no effect). However, even for such large matrices, the effect of the correlation (line colors) is noticeable. For each choice of correlation, the value *α* = *α*_*S*_ at which the stability of a feasible equilibrium is guaranteed is marked by a triangle: note that a negative *ρ* increases both the probability of feasibility and stability for any *α*, and a positive *ρ* decreases it[16]. Moreover the probability of feasibility at *α*_*S*_ is negligible. The black solid lines represent the analytical expectations obtained by approximating the variance using resolvents (Eqs. (3) and (4)).

Importantly, our calculations show that for large systems the amount of self-regulation required for feasibility is high enough (i.e., *α*_*S*_ <*α*_*F*_) to yield global stability (as argued before[9, 14, 20, 32, 33]). Then, a feasible system will be stable with probability 1, and thus *p*_*F*_ (*α*) measures the probability of coexistence for a large, random competitive system. These results have profound implications for dynamics. Competitive GLV systems can give rise to any type of dynamics, including limit cycles (when *n* ≥ 3) and chaos (*n* ≥ 4)[23, 24]. Importantly, however, non-equilibrium coexistence still requires the existence of a feasible equilibrium[25]. But we have seen that feasibility generically implies global stability, and therefore non-equilibrium coexistence is precluded.

When the system is not feasible, extinctions will ensue, pruning the original pool of species to a subsystem with different thresholds for feasibility and stability (Fig. 2c). As such, in principle, any type of dynamics could be observed. Next, we show that, despite the change in values of *α*_*S*_ and *α*_*F*_ caused by extinctions, the ordering *α*_*S*_ <*α*_*F*_ is maintained almost surely, provided that *n* is large. As a result, dynamics still converge to stable equilibria even when feasibility is not attained for the initial species pool.

### Robust coexistence emerges through community dynamics

So far, we have investigated how coexistence is affected by increasing *α*. However, intraspecific competition models the effects of a species on itself. As such, when species are lost to extinction, the level of intraspecific competition should not be affected: the coefficients on the diagonal do not change when the matrix is pruned of the populations that have gone extinct. Thus, we turn our attention to a pool of *n* populations where we fix *α < α*_*S*_ <*α*_*F*_ (analogous to system I in Fig. 1), and consider “top-down assembly” (where all the species in the pool are introduced in the environment at the same time, Fig. 4a). Since we are below the feasibility threshold, some populations go extinct and, as a result, *α*_*F*_ and *α*_*S*_ changes (both values depend on *n*; Fig. 2c). We re-compute their values every time an extinction occurs (plotted in Fig. 4b), and find that their initial ordering is maintained throughout assembly. Consequently, the initial species pool traverses the dynamical states illustrated in Fig. 1: I → II → III, converging to a globally stable equilibrium after ecological dynamics have elapsed. Furthermore, we show (Supplementary Note 3, Sec. 12) that the same equilibrium can be achieved via “bottom-up assembly”, through successive invasions[26].

**Figure 4.**
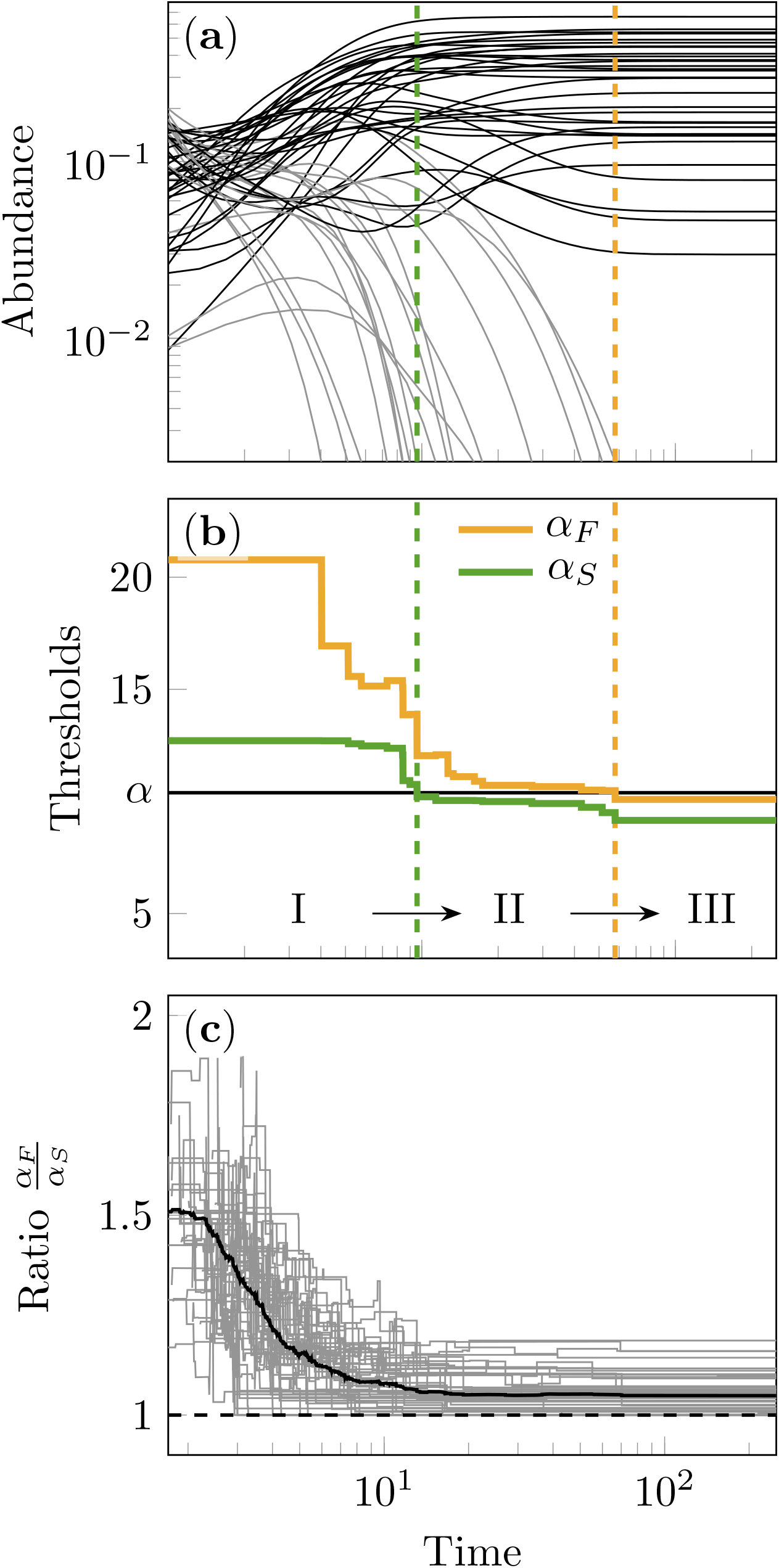
Community dynamics lead to robust coexistence.**(a)** We initialize a community with *n* = 50 populations and random interactions, setting intraspecific competition below the feasibility threshold (*α* = *α*_*F*_ */*2, black line, middle panel); then, there is no feasible equilibrium, and extinctions are guaranteed (grey lines, top panel); note that for this simulation intraspecific competition is also below the stability threshold (*α < α*_*S*_ <*α*_*F*_), and thus we are in phase I of Fig. 4. **(b)** As populations are lost to extinction, the values of *α*_*S*_ and *α*_*F*_ (green and yellow lines respectively, middle panel) for the surviving sub-community change. As the number of extinctions mounts, *α*_*S*_ crosses *α* (dashed green line, top and middle panels), and thus the system transitions to case II. Extinctions continue until *α*_*F*_ falls below *α* (dashed yellow line), and the system transitions to phase III. The extant populations coexist at a globally stable equilibrium, and those that went extinct since during phase II cannot re-invade the system. **(c)** The ratio *α*_*F*_ */α*_*S*_ is plotted as a function of time for many random realizations (individual trajectories in gray, average in black) of the communities with the same statistics as in the example. The ratio is always above 1, meaning that the relationship *α*_*F*_ *> α*_*S*_ maintains through the dynamics in all simulations.

To confirm that these observations are a generic feature, we plot the ratio *α*_*F*_ */α*_*S*_ when we run the same simulation in the previous example for many draws of random systems with the same characteristics (Fig. 4c). We see that the ratio is kept above one, implying that the relationship *α*_*S*_ <*α*_*F*_ is maintained throughout the dynamics for all the simulations. This shows that, with high probability, the dynamical outcome of large random competitive systems is always the same: a sub-set of the species pool will coexist robustly at an equilibrium that is characterized by *α*_*S*_ <*α*_*F*_ <*α*; the equilibrium is stable, and the last few populations to go extinct (if started in phase I), or all the extinct populations (if started in phase II) cannot re-invade the system.

## Discussion

Ecological dynamics can be extremely complex, with populations’ trajectories oscillating regularly or chaotically. The Generalized Lotka-Volterra model for multiple competitors embodies this complexity: for less than three competitors, trajectories always converge to steady states; for three or more species, one can observe cyclic dynamics; four or more competitors can coexist via nonperiodic, chaotic oscillations. Importantly, to ensure the long-term coexistence of the populations there must exist a feasible equilibrium: a choice of positive densities at which all populations, if unperturbed, could remain indefinitely. Here we ask what types of dynamics arise when the number of competitors is large and interactions are chosen at random: we derive the probability of feasibility, and in doing so, notice that as intraspecific interaction is increased, the transition to stability always precedes that for feasibility when the system is large. As a consequence, feasible systems will be stable. Importantly, all the results in this paper are true in probability. That is, while the probability of finding feasible but unstable systems is never exactly zero, it becomes vanishingly small with system size. Intraspecific competition models the fact that each species has access to private resources or regulating factors that are species-specific; as such, it plays an important role in niche differentiation, and sets a limit to niche similarity[1–6]. Our results show that, to ensure feasibility, one had to postulate a level of niche differentiation that is sufficient to guarantee stability. This observation reinforces the notion that, in competitive systems, feasibility is key for coexistence[5, 8, 22, 34]. This is in analogy with consumer-resource models[35], as well as in GLV models with symmetric interactions produced by niche overlap or species’ similarity[36], for which stability is guaranteed, and thus feasibility determines coexistence[26, 37]. Thus, our results predict that mechanisms that decrease effective niche overlap—whether metabolic, spatial, or trait-based—push the system past the feasibility threshold *α*_F_. While direct empirical test of these patterns is out of the scope of this work, it is an exciting avenue for future research. For example, in microbial systems one could quantify metabolic overlap from shared nutrient-uptake profiles or genome-encoded pathways, and use this measure to estimate how reductions in overlap map onto the effective self-regulation needed to surpass the feasibility threshold.

As a consequence, we have shown that, when the number of competitors is large, trajectories converge to stable steady states with high probability, ensuring the robust coexistence of the extant competitors. These findings are in line with empirical results: experimenting with organisms at the same trophic level (for which competition is believed to be the dominant mode of interaction) often results in equilibrium dynamics [38– 43], while sustained cycles and chaos are typically associated with predator-prey systems and structured populations[44–49].

Recently, it has been shown[50] that a dynamical phase characteristic of persistent fluctuations can arise for the high-diversity, high-interaction strength limit in bacterial systems dominated by competitive interactions. Furthermore, this observation mirrors dynamical phases arising in a GLV model with dispersal. Crucially, these findings hinge on the presence of constant immigration—without it, irrespective of interaction strength, robust coexistence emerges for large communities (see Fig. S1 in[50] for results when *n* = 50). As such, our findings are not in contradiction with these previous results[50]—including constant immigration for all species precludes extinctions, thereby preventing the system from relaxing to the equilibria studied here. Therefore, the results presented can be used to form an expectation for isolated systems in which populations can go extinct, or systems in which immigration is rare. For open systems in which immigration is widespread, outcomes could depart severely from what presented here.

In summary, we have shown that, for large competitive communities with random coefficients following the GLV model, the probability of non-equilibrium coexistence is negligible—feasible systems are almost surely stable, and thus chaos or limit cycles will not be observed. Moreover, coexistence via stable equilibria emerges through community dynamics. In conclusion, when considering the subset of coexisting populations, we find that each population rebounds from perturbations, that the community cannot be invaded and can be built through successive invasions—the hallmark of robust coexistence [4, 6].

## Methods

### A canonical form for competitive systems

We consider the competitive GLV system in Eq. (1), with **x** > **0**, *A* = *αI* + *B* and *B*^*T*^ **1** > **0**. We can further decompose the matrix *A* = *αI* + *C* + **1m**^*T*^, where *C* is the matrix obtained by subtracting from each column of *B* its (positive) mean, and **m** > **0** is a vector containing the column means. We can write the solution of (*αI* + *C* + **1m**^*T*^)**x**^⋆^ = **1** as a function of the solution of the simpler system (*αI* + *C*)**y**^⋆^ = **1** (Supplementary Note 2, Sec. 1), finding:

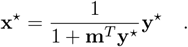

Thus, **x**^⋆^ is related to **y**^⋆^ through multiplication by the scalar 1*/*(1 + **m**^*T*^ **y**^⋆^). As such, we have a feasible equilibrium **x**^⋆^ > **0** when either a) **y**^⋆^ > **0** (i.e., **y**^⋆^ is feasible, and because 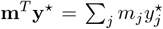 is the sum of products of nonnegative values, then the constant of proportionality is also positive), or, b) **y**^⋆^ < **0** and **m**^*T*^ **y**^⋆^ < −1 (i.e., all the components of **y**^⋆^ are negative, and sufficiently large to make the constant of proportionality negative as well). However, the average of **y**^⋆^ is positive (Supplementary Note 2, Sec. 1), thereby ruling out case b). We conclude that **x**^⋆^ is feasible if and only if **y**^⋆^ is feasible. Because infinitely many competitive matrices *B* will map into the same *C*, we say that this is the canonical matrix of interactions for the competitive system. Although increasing the strength of competition *µ* does not affect feasibility, it decreases equilibrium abundance values. As such, increasing *µ* too much will result in very low abundances, which could lead to extinctions in real systems where stochasticity plays a factor.

### Equivalence between α and σ

Consider a matrix of interactions *C*, such that the variance of *C*_*ij*_ is *σ*^2^. We show that there exists a rescaled system with the same feasibility properties, a different *α*, and a scaled interaction matrix such that *σ*^2^ = 1. The equilibrium for the system is

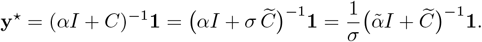

Setting **z** = *σ***y**^⋆^ we have that 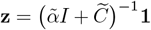, where 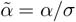 and 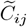 have unit variance. Since *σ >* 0, then **z** > **0** ⇔ **y**^⋆^ > **0**. Therefore, varying *σ* (i.e., the variance of the interaction matrix) is equivalent to varying intraspecific competition *α*. Without loss of generality, we can fix *σ*^2^ = 1 and study solely the effect of *α*.

### Intraspecific competition guaranteeing global stability

The Wigner semicircular law for symmetric matrices with i.i.d. entries with mean zero and variance 1*/n*, 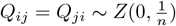, states that, as *n* → ∞, the eigenvalues of *Q* are contained in the semicircle centered at 0 with radius 2 almost surely. Because *Q* is symmetric, all its eigenvalues are real, and we can write

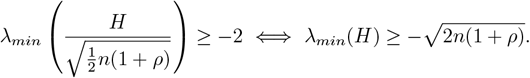

We consider a random interaction matrix *B* ∈ ℝ^n*×*n^ with 𝔼 [*B*_*ij*_] = 0, 𝕍 (*B*_*ij*_) = 1, and ℂ 𝕠 𝕣 (*B*_*ij*_, *B*_*ji*_) = *ρ*, and introduce intraspecific competition via *A* = *αI* + *B*. A standard argument based on Lyapunov functions (see Supplementary Note 1, Sec. 3) shows that global stability of the coexistence equilibrium is guaranteed by the positive definiteness of the symmetric part of *A*,

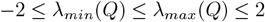

Let *H*:= (*B* + *B*^*T*^), then, 𝔼 (*H*_*ij*_) = 0, and 𝕍 (*H*_*ij*_) = ½ (1 + *ρ*). As such, the eigenvalue

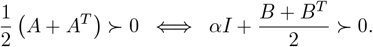

Therefore, positive definiteness of *A* + *A*^*T*^ is guaranteed as long as intraspecific competition is above the threshold 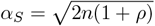.

### Intraspecific competition guaranteeing feasibility

One can show that, in competitive systems, there exists a level of intraspecific competition at which the feasibility of the equilibrium is guaranteed, and further strengthening intraspecific competition retains feasibility (Supplementary Note 2, Sec. 2). We denote this critical level as *α*_*F*_. To find this value for a given matrix of interaction strengths *B*, we consider the eigenvalues of several matrices derived from *B* (or, equivalently, *C*, Supplementary Note 2, Sec. 3). In particular, consider the real, positive eigenvalues of −*C* = **1m**^*T*^ − *B*, and call *α*_0_ the largest value such that the components of the solution (*α*_0_*I* +*C*)**y**^⋆^ = **1** diverge to ±∞ (if it exists, otherwise *α*_0_ = 0). Then, we form *n* matrices *Q*_*i*_, each of size *n* − 1. Each *Q*_*i*_ is obtained as the sum of two matrices, *Q*_*i*_ = **1**(**b**^(*i*)^)^*T*^ − *B*^(*i*)^, where **b**^(*i*)^ is the *i*^*th*^ row of matrix *B* with the *i*^*th*^ element removed, and *B*^(*i*)^ is the matrix *B* with the *i*^th^ row and the *i*^th^ column removed. Denote as *α*_*i*_ the maximum real eigenvalue of *Q*_*i*_ (if it exists, otherwise *α*_*i*_ = 0). Finally, we have that the value *α*_*F*_ that guarantees the feasibility of the system is *α*_*F*_ = max{*α*_0_, *α*_1_, *α*_2_, …, *α*_*n*_}.

Interestingly, the value of intraspecific competition sufficient to guarantee the global stability of any feasible equilibrium is also an eigenvalue, this time the largest positive eigenvalue of −(*B* + *B*^*T*^)*/*2 (see above, and Supplementary Note 1, Sec. 3).

### Distribution of y^⋆^ for systems with random parameters

For random models in which populations are statistically equivalent (Supplementary Note 3, Sec. 1), we have that the expectations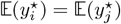, the variances 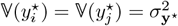, and the correlation 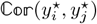 is the same for all pairs. Note that the sum of the components of **y**^⋆^ is fixed: **1**^*T*^ **y**^⋆^ = *n/α*, and therefore the mean correlation must be negative; because it has to be the same for all pairs, it is exactly 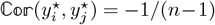, as shown also by calculating it explicitly (Supplementary Note 3, Sec. 7). Moreover, if one approximates the distribution of **y**^⋆^ as a multivariate normal distribution, finding the unknown variance (which will depend on the exact definition of the model, the statistics of the distribution, etc.) is sufficient to approximate the probability, by either ignoring the correlation between the components (because for large *n* it is negligible), or by computing an integral over the positive orthant (Supplementary Note 3, Sec. 7).

## Data availability

No data was used in this work

## Code availability

Code to reproduce all findings is available at

https://doi.org/10.5281/zenodo.17742734

## Acknowledgements

A. Skwara, Z.R. Miller, and C. Serván for discussion. This research was supported in part by grants from the NSF (DMS-2235451) and Simons Foundation (MPS-NITMB-00005320) to the NSF-Simons National Institute for Theory and Mathematics in Biology (NITMB). JAC acknowledges financial support from grant PRIORITY (PID2021-127202NB-C22), funded by MCIN/AEI/10.13039/501100011033 and “ERDF. A way of making Europe”. We acknowledge the University of Chicago’s Research Computing Center for their support of this work.

## Author contributions

SA conceived and designed the study. PLA and SA drafted the manuscript. PLA drafted the code and the figures. SA and JAC performed the derivations; JAC derived the approximation using resolvents; SA derived those based on iterative solutions and small-rank perturbations. All Authors contributed to the mathematical derivations, the implementation of the code, edited the manuscript, and discussed and interpreted the results.

