## supplementary information for "Robust coexistence in competitive ecological communities"

### Supplementary Note 1. Preliminaries: competitive Generalized Lotka-Volterra models

We consider the dynamics of ecological communities composed of $n$ interacting populations, and following the Generalized Lotka-Volterra (GLV) model[1–3]:

$$\dot{x}_i = x_i r_i \left(1 - \sum_j A_{ij} x_j\right) , \tag{1}$$

where $x_i(t) \equiv x_i$ is the density of population $i$ at time $t$, $\dot{x}_i \equiv dx_i(t)/dt$ its time derivative, $r_i$ is the intrinsic growth/death rate of population $i$, achieved when growing in isolation at low abundance, and $A_{ij}$ is the amount by which population $j$ depresses/enhances the growth of population $i$.

We analyze cases in which all populations can grow in isolation ($r > \mathbb{0}$), and, moreover, interactions are "mostly competitive" (defined below). Following May[4], we write $A$ as the sum of two matrices:

$$A = \alpha I + B \quad ,$$

with $\alpha \geq 0$. Then, $B$ collects all the inter-specific interactions, as well as the baseline levels of intraspecific interactions, and $\alpha$ can be interpreted as the "excess" intraspecific competition in the system.

We call the system *competitive* whenever $r > \mathbb{0}$, and $\frac{1}{n} B^T \mathbb{1} = m > \mathbb{0}$ (i.e., the column means of $B$ are all positive, and thus the columns of $B$ sum to positive values). While not *every* interaction needs to be competitive, each population must, on average, depress the growth of the others.

An equilibrium $x^\star$ for this system is a point at which all the derivatives in Eq. (1) vanish. We say that the equilibrium is feasible if $x^\star > \mathbb{0}$. In this work, we analyze the effect of $\alpha$ on the existence of a feasible equilibrium for competitive GLV systems.

#### 1.1 Local stability

The goal of this section is to show that there exists a level of $\alpha = \alpha_L$, corresponding to the celebrated May's bound[4], that guarantees that all the eigenvalues of $A$ have positive real part, and discuss its implication for local stability.

The eigenvalues of $A$ are related to the eigenvalues of $B$, which are generically complex numbers of the form $\lambda_j(B) = \gamma_j + i\beta_j$, with $i = \sqrt{-1}$. Because the matrix $B$ is composed of real numbers, whenever $\lambda_j(B) = \gamma_j + i\beta_j$ is an eigenvalue, then $\lambda_k(B) = \gamma_j - i\beta_j = \lambda_j^*(B)$, i.e., its conjugate, is also an eigenvalue. Adding a constant diagonal $\alpha I$ to $B$ results in a shift in the real part of all the eigenvalues: $\lambda_j(A) = \lambda_j(\alpha I + B) = \alpha + \gamma_j + i\beta_j$. Hence, whenever $\alpha > \alpha_L = \min\{0, \min_j \mathcal{Re}(\lambda_j)\}$, all eigenvalues of $\alpha I + B$ are contained in the right half of the complex plane, and thus have positive real part.

This is important for the local stability of an equilibrium of the GLV system in Eq.(1). An equilibrium $x^\star$ is said to be *locally asymptotically stable* if trajectories originating arbitrarily close to it eventually converge to it. An equilibrium is locally stable if all the eigenvalues of the Jacobian matrix of the system of differential equations, evaluated at the equilibrium, have negative real part. The Jacobian for Eq.(1) evaluated at $x^\star > \mathbb{0}$ is simply:

$$J|_{x^\star} = -D(x^\star \circ r)A \quad ,$$

where $D(z)$ is a matrix with the vector $z$ on the diagonal, and $\circ$ represents the Hadamard (element by element) product. Hence, if all the eigenvalues of $A$ have positive real part, and we were to choose $r$ such that $r_i = 1/x_i^\star$, the equilibrium would be locally asymptotically stable.

Whenever $A = \alpha I + B$, we have that both $A$ and $x^\star$ depend on $\alpha$, while the vector $r$ does not. We can write:

$$J|_{x^\star} = D(r)(-D(x^\star)A) \quad ,$$

to stress the fact that only the part in parenthesis is influenced by $\alpha$. Importantly, we could have that $-D(x^\star)A$ has only eigenvalues with a negative real part, while $J|_{x^\star}$ does not. For example, take the competitive GLV system with:

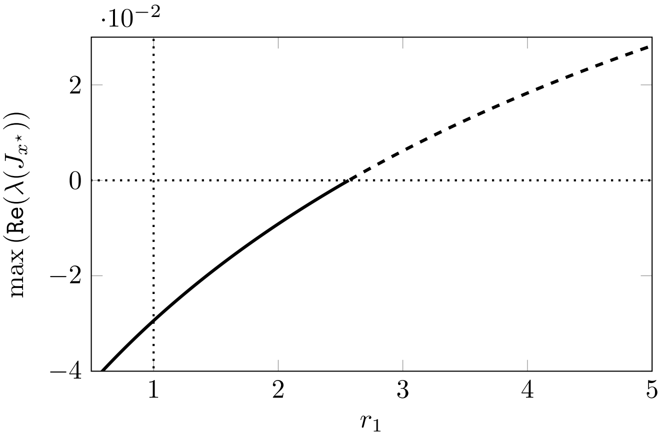


**Supplementary Figure 1**: Bifurcation diagram obtained by varying the growth rates for the three species system in Eq. (2). When the growth rates are chosen as $r = (r_1, 1, 1)^T$, the system experiences a transition from stability to instability as the maximum of $\mathscr{Re}(\lambda(J|_{x^\star}))$ changes sign from negative to positive. The bifurcation occurs at about $r_1 = 2.57$ as $r_1$ is varied in the range $[1/2, 5]$, but the system remains feasible throughout.

$$A = \begin{pmatrix} 1 & 2 & 3 \\ 2 & 3 & 0 \\ 0 & 3 & 1 \end{pmatrix} \qquad x^\star = \frac{1}{17}\begin{pmatrix} 1 \\ 5 \\ 2 \end{pmatrix} \quad . \tag{2}$$

The system is feasible (for any choice of $r > \mathbb{0}$). However, the local stability of the equilibrium depends on the choice of $r$. For $r = \mathbb{1}$, we have that the eigenvalues of $J|_{x^\star}$ are $(-1, -\frac{1}{34}(1 \pm i\sqrt{39}))$ and the equilibrium is thus locally stable. However, by increasing the value of $r_1$ we observe a bifurcation: the real part of the maximum eigenvalue becomes positive and the system loses stability, as shown in Figure 1.

Thus, in general, the stability of a feasible equilibrium $x^\star > \mathbb{0}$ might depend on the choice of $r$. However, if $A$ is "sufficiently" stable, then the stability of any *feasible* equilibrium is guaranteed for *any* choice of $r > \mathbb{0}$, as discussed below.

#### 1.2 Types of dynamics

Despite its simplicity, the competitive GLV model can give rise to any type of dynamics, including limit cycles and chaos. For purely competitive systems with positive growth rates, when there is a single population, dynamics will always converge to an equilibrium; for two populations, dynamics will generically converge to an equilibrium at which point either population, or both, are present; for three or more populations, one can observe limit cycles, and chaos for four or more populations[5, 6].

Importantly, a necessary condition for the coexistence of all the populations is the existence of a feasible equilibrium $x^\star > \mathbb{0}$, which is also the time-average of the orbits[7]. When such a feasible equilibrium does not exist, trajectories can either reach the boundaries of $\mathbb{R}^n_+$ (i.e., populations undergo extinction), or escape to infinity. When interactions are purely competitive, populations limit each others' growth and divergences to infinity are precluded; therefore, in the absence of a feasible equilibrium, some populations will necessarily go extinct.

##### 1.3 Global stability and saturated equilibria

The goal of this section is to show that there exists a further bound $\alpha = \alpha_S \geq \alpha_L$ that guarantees that a) any feasible equilibrium is locally asymptotically stable, i.e., whenever the densities are slightly perturbed away from the equilibrium, they will eventually return to it; b) the equilibrium is in fact *globally* stable, i.e., any perturbation short of extinction will eventually subside; c) dynamics are guaranteed to result in an equilibrium; d) this equilibrium is *saturated*: some of the populations coexist at a feasible, globally stable equilibrium, and the rest of the populations are extinct and cannot re-invade the system when introduced at low abundance.

In the section discussing local stability, we have shown that choosing $\alpha = \alpha_L$ guarantees local stability for a specific choice of $r$. However, whenever $A + A^T$ is positive definite (i.e., has only positive eigenvalues), then a) dynamics will always converge to an equilibrium, $\bar{x}$; b) the equilibrium $\bar{x}$ is saturated, meaning that if any population goes extinct, it cannot re-invade the system when starting from low abundance; c) $\bar{x}$ is globally stable (i.e., for any choice of $r > \mathbb{0}$ and initial conditions $x > \mathbb{0}$, dynamics asymptotically converge to $\bar{x}$).

We adapt the proof found for example in Hofbauer & Sigmund[7], because this result is important for our discussion on the dynamics of large competitive communities with random interactions.

We order the vector of densities of the populations in Eq. (1) as $x = (y, z)^T$, where $y$ are the densities of the (unknown) set of the $k$ populations that coexist at the saturated equilibrium, and $z$ the densities of the $n - k$ populations that are extinct at the saturated equilibrium. Therefore $\bar{x} = (y^\star, \mathbb{0})^T$. We also consider a vector of positive weights $w > \mathbb{0}$, which we will use to prove stability. We partition the vector of growth rates, the vector of weights $w$, and the matrix of interactions according to whether the populations belong to the set that coexists, or that that goes extinct:

$$w = \begin{pmatrix} u \\ v \end{pmatrix} > \mathbb{0} \quad r = \begin{pmatrix} a \\ b \end{pmatrix} > \mathbb{0} \quad A = \begin{pmatrix} \mathcal{A} & \mathcal{B} \\ \mathcal{C} & \mathcal{D} \end{pmatrix} \quad .$$

Next, we consider the candidate Lyapunov function[8]:

$$V(y, z) = \sum_i u_i \left( y_i - y_i{}^\star - y_i{}^\star \log \frac{y_i}{y_i{}^\star} \right) + \sum_l v_l z_l \quad .$$

The function is positive for any $y > \mathbb{0}$ and $z > \mathbb{0}$ (i.e., for any positive initial conditions), given that $u, v, y^\star$ are all positive. Moreover, the function is exactly zero at the saturated equilibrium $\bar{x} = (y^\star, \mathbb{0})^T$. We choose $u_i = 1/a_i$, and $v_l = 1/b_l$, and then differentiate with respect to time:

$$\frac{dV(y, z)}{dt} = \sum_i (y_i - y_i{}^\star) \left( 1 - \sum_j \mathcal{A}_{ij} y_j - \sum_k \mathcal{B}_{ik} z_k \right) + \sum_l z_l \left( 1 - \sum_j \mathcal{C}_{lj} y_j - \sum_k \mathcal{D}_{lk} z_k \right) \quad .$$

We define $\Delta y_i = y_i - y_i{}^\star$; noticing that $1 = \sum_j \mathcal{A}_{ij} y_j{}^\star$ we can write:

$$\frac{dV(y,z)}{dt} = -\sum_{i,j} \Delta y_i \mathcal{A}_{ij} \Delta y_j - \sum_{i,k} \Delta y_i \mathcal{B}_{ik} z_k - \sum_{l,k} z_l \mathcal{D}_{lk} z_k + \sum_l z_l \left(1 - \sum_j \mathcal{C}_{lj} y_j\right) \quad .$$

Now we add and subtract $\sum_l z_l \sum_j \mathcal{C}_{lj} y_j{}^\star$, thereby obtaining:

$$\begin{aligned} \frac{dV(y,z)}{dt} = &-\sum_{i,j} \Delta y_i \mathcal{A}_{ij} \Delta y_j - \sum_{i,k} \Delta y_i \mathcal{B}_{ik} z_k - \sum_{l,j} z_l \mathcal{C}_{lj} \Delta y_j \\ &- \sum_{l,k} z_l \mathcal{D}_{lk} z_k + \sum_l z_l \left(1 - \sum_j \mathcal{C}_{lj} y_j{}^\star\right) \quad . \end{aligned}$$

Thus, $dV(y,z)/dt$ is the sum of two parts. The first part can be rewritten as a quadratic form:

$$\begin{aligned} -\sum_{i,j} \Delta y_i \mathcal{A}_{ij} \Delta y_j - \sum_{i,k} \Delta y_i \mathcal{B}_{ik} z_k - \sum_{l,j} z_l \mathcal{C}_{lj} \Delta y_j - \sum_{l,k} z_l \mathcal{D}_{lk} z_k = -\sum_{i,j} (x_i - \bar{x}_i) A_{ij} (x_j - \bar{x}_j) = \\ -(x-\bar{x})^T A (x - \bar{x}) = -\frac{1}{2}(x-\bar{x})^T \left(A + A^T\right)(x - \bar{x}) \quad , \end{aligned}$$

and is thus negative whenever $A + A^T$ is positive definite.

Finally, $(1 - \mathcal{C}y^\star)_j$ is positive only when population $j$ can grow (and thus invade the system) when the system is resting at the saturated equilibrium. Hence, by the definition of a saturated equilibrium, this term is also negative, and the proof is complete.

Note that the real parts of the eigenvalues of $A$ are bounded from above and below by those of $H = \frac{1}{2}(A + A^T)$, as stated by Bendixson's inequality; in particular, if we order the eigenvalues of $H$ in ascending order, $\lambda_1(H) \leq \lambda_2(H) \leq \ldots \leq \lambda_n(H)$, we have

$$\lambda_1(H) \leq \mathcal{R}e \lambda_i(A) \leq \lambda_n(H) \quad ,$$

with equality guaranteed for symmetric matrices $A$ (in which case we have $H = A$). Thus, we have that $\alpha_L \leq \alpha_S$ (as expected, given that a globally stable equilibrium must be locally stable as well).

In summary, we have shown that, provided that $r > \mathbb{0}$ and that we chose $\alpha > \alpha_S$ such that the symmetric part of $A$, $H$, is positive definite, dynamics will always converge to an (unspecified) saturated equilibrium. This is a *sufficient* condition for dynamics to converge to a saturated equilibrium, which can be made slightly more general. For example, whenever one can choose a positive vector $h > \mathbb{0}$ such that $D(h)A + A^T D(h)$ is positive definite, then the stability of dynamics is also ensured[8]. Because we are interested in an upper bound that is easy to calculate, we just analyze the case in which $H$ is positive definite.

##### 1.4 Assembly

This section discusses the assembly of systems for which $\alpha > \alpha_S$, i.e., whether the systems can be built from the bottom up through successive invasions. We can devise an "assembly graph" in which the nodes represent the subset of populations that can coexist, and the edges invasions moving the system from one state to another[9, 10]. Naturally, this graph representation is an especially good choice for systems in which the dynamics of any feasible sub-community settle at an equilibrium, invasions are rare, and spaced apart in time[9].

When $\alpha > \alpha_S$, then this relationship is maintained for any sub-system (because all the sub-matrices of $H$ obtained by removing some rows and their corresponding columns are also positive definite). Hence, any feasible sub-community would also be (globally) stable, and might or might not be invasible by other populations. Take two sub-communities $x_k$, composed of $k$ populations and $x_{k+1}$ composed of the same populations, but with another population added. If both $x_k$ and $x_{k+1}$ are feasible, and both are stable because $\alpha > \alpha_S$, then the population that is added can attach itself to the resident community without causing extinctions.

Suppose that a GLV system has $\alpha > \alpha_S$ and a feasible equilibrium $x^\star > \mathbb{0}$. Then, determining whether there is a way to build the community from the ground up amounts to determining whether there is a path in the graph such that, at each step, a different population attaches itself to the community without causing extinctions.

### Supplementary Note 2. General systems

We start our discussion on the effect of the strength of intraspecific competition $\alpha$ on coexistence by considering the case in which matrix $B$ is a generic matrix. In the next section, we analyze the case in which $B$ is a large random matrix instead.

##### 2.1 Effect of average interaction strength

Here we show that the feasibility of the equilibrium $x^\star$ in Eq. (1) is related to that of the equilibrium of a simpler system, $y^\star$, thereby obtaining a canonical form for the study of feasibility in competitive GLV systems.

The system in Eq. (1) is feasible whenever $x^\star = A^{-1}\mathbb{1} = (\alpha I + B)^{-1}\mathbb{1} > \mathbb{0}$. We can write:

$$A = \alpha I + B = \alpha I + C + \mathbb{1} m^T \quad ,$$

where the columns of $C$ sum to zero ($C^T \mathbb{1} = \mathbb{0}$), the vector $m \geq \mathbb{0}$ (because the system is competitive), $I$ is the identity matrix of dimension $n$, and $\alpha \geq 0$. The rank-one matrix $\mathbb{1} m^T$ has constant columns with nonnegative coefficients, corresponding to the column means of $B$. Now consider the system:

$$(\alpha I + C) y^\star = \mathbb{1} \quad . \tag{3}$$

Note that the sum of all the $y_i^\star$ is positive:

$$\mathbb{1}^T(\alpha I + C)y^\star = \mathbb{1}^T\mathbb{1}$$
$$\alpha\mathbb{1}^T y^\star + \mathbb{1}^T C y^\star = n$$
$$\frac{1}{n}\mathbb{1}^T y^\star + 0 = \frac{1}{\alpha} > 0 \quad .$$

As such, the mean of the components of $y^\star$ is exactly $1/\alpha$, and is thus positive for any positive $\alpha > 0$. We can write $x^\star$, the solution of $Ax^\star = \mathbb{1}$, as a function of $y^\star$ using the Sherman-Morrison formula:

$$(\alpha I + B)x^\star = \mathbb{1}$$
$$(\alpha I + C + \mathbb{1}m^T)x^\star = \mathbb{1}$$
$$x^\star = (\alpha I + C + \mathbb{1}m^T)^{-1}\mathbb{1}$$
$$x^\star = \left((\alpha I + C)^{-1} - \frac{1}{1 + m^T(\alpha I + C)^{-1}\mathbb{1}}(\alpha I + C)^{-1}\mathbb{1}m^T(\alpha I + C)^{-1}\right)\mathbb{1}$$
$$x^\star = y^\star - \frac{1}{1 + m^T y^\star} y^\star m^T y^\star$$
$$x^\star = \left(1 - \frac{m^T y^\star}{1 + m^T y^\star}\right) y^\star$$
$$x^\star = \frac{1}{1 + m^T y^\star} y^\star \quad .$$

Thus, $x^\star$ simply related to $y^\star$, through multiplication by the scalar $1/(1 + m^T y^\star)$. As such, we have a feasible equilibrium $x^\star > \mathbb{0}$ whenever a) $y^\star > \mathbb{0}$ (i.e., $y^\star$ is feasible, and because $m^T y^\star = \sum_j m_j y_j^\star$ is the sum of products of nonnegative values, then the constant of proportionality is also positive), or, b) $y^\star < \mathbb{0}$ and $m^T y^\star < -1$ (i.e., all the components of $y^\star$ are negative, and sufficiently large to make the constant of proportionality negative as well). But we have just seen that the average of $y^\star$ is positive, thereby making case b) impossible. We conclude that $x^\star$ is feasible if and only if $y^\star$ is feasible.

Because infinitely many competitive systems in Eq. (1) map into the same Eq. (3), we say that the equation in $C$ is the *canonical* form for studying the feasibility of the competitive systems that can be formed by adding positive columns to $C$.

#### 2.2 A sufficiently large $\alpha$ ensures feasibility

Here we use the well-known Perron-Frobenius property for nonnegative matrices to show that for sufficiently large $\alpha = \alpha_\infty$ the feasibility of $y^\star$ (and thus of $x^\star$) is guaranteed, and is maintained for any $\alpha > \alpha_\infty$. To this end, we divide both sides of Eq. (3) by $\alpha > 0$:

$$\left(I + \frac{1}{\alpha}C\right) y^\star = \frac{1}{\alpha}\mathbb{1} \quad .$$

We have shown above that $\mathbb{1}^T y^\star = n/\alpha$, but then $\frac{\alpha}{n}\mathbb{1}^T y^\star = 1$. Multiplying the right hand side by this value, we get:

$$\left(I + \frac{1}{\alpha}C\right) y^\star = \frac{1}{n}\mathbb{1}\mathbb{1}^T y^\star \quad .$$

Bringing the right-hand side to the left:

$$\left(I + \frac{1}{\alpha}C - \frac{1}{n}\mathbb{1}\mathbb{1}^T\right) y^\star = \mathbb{0}$$

and, finally,

$$\left(\frac{1}{\alpha}C - \frac{1}{n}\mathbb{1}\mathbb{1}^T\right) y^\star = -y^\star$$
$$\left(\frac{1}{n}\mathbb{1}\mathbb{1}^T - \frac{1}{\alpha}C\right) y^\star = y^\star \quad ,$$

which shows that $y^\star$ is a right eigenvector of $\frac{1}{n}\mathbb{1}\mathbb{1}^T - \frac{1}{\alpha}C$, with associated eigenvalue $\lambda = 1$. Note that the matrix $\frac{1}{n}\mathbb{1}\mathbb{1}^T - \frac{1}{\alpha}C$ has left eigenvector $\mathbb{1}$, also associated with $\lambda = 1$. Now suppose we choose $\alpha = \alpha_\infty > n \max_{i,j} C_{ij}$; then, we have that a) the matrix $\frac{1}{n}\mathbb{1}\mathbb{1}^T - \frac{1}{\alpha_\infty}C$ has only positive entries; b) it is column stochastic (i.e., it is nonnegative with columns summing to 1), and therefore has spectral radius 1. But then $\frac{1}{n}\mathbb{1}\mathbb{1}^T - \frac{1}{\alpha_\infty}C$ has the Perron-Frobenius property, and thus $y^\star$ is positive, and the positivity of $y^\star$ is guaranteed for any $\alpha \geq \alpha_\infty$. Naturally, this choice of $\alpha = \alpha_\infty$ is overly conservative, but it is sufficient to show that there must exists some critical value $\alpha_F \leq \alpha_\infty$ such that: a) $y^\star$ is positive, and b) it remains positive for any $\alpha > \alpha_F$. Computing such a value is the main goal of the next two subsections.

#### 2.3 Transitions to and from feasibility

We can track the value of $y_i^\star(\alpha)$, as we vary $\alpha$, using Cramer's rule:

$$y_i^\star(\alpha) = \frac{\begin{vmatrix} \alpha + C_{1,1} & C_{1,2} & \cdots\ 1\ \cdots & C_{1,n} \\ C_{2,1} & \alpha + C_{2,2} & \cdots\ 1\ \cdots & C_{2,n} \\ \vdots & \vdots & \vdots & \vdots \\ C_{n,1} & C_{n,2} & \cdots\ 1\ \cdots & \alpha + C_{n,n} \end{vmatrix}}{|\alpha I + C|} = \frac{|Q_i(\alpha)|}{|Q(\alpha)|} \quad ,$$

where $Q(\alpha) = \alpha I + C$, and $Q_i(\alpha)$ is obtained by substituting the $i^{\text{th}}$ column of $Q(\alpha)$ with the vector $\mathbb{1}$.

When $|Q(\alpha)| = 0$, the matrix is singular; this means that $C$ has a real, negative eigenvalue $\lambda_i$ such that by setting $\alpha = -\lambda_i$ we place an eigenvalue of $Q(\alpha)$ exactly at zero. We have two cases: if, as $\alpha \to -\lambda_i$, the determinant of the numerator $|Q_i(\alpha)|$ tends to a nonzero value, then $y_i^\star(\alpha)$ diverges as $\alpha \to -\lambda_i$; if instead $|Q_i(\alpha)| \to 0$ as $\alpha \to -\lambda_i$, then the solution $y_i^\star(\alpha)$ does not diverge around $\alpha = -\lambda_i$. Importantly, given that the mean of the solution is fixed and positive, whenever some solutions diverge, some components will tend to $+\infty$, while others to $-\infty$ around these singularities. Hence, the system cannot be feasible around any singularity that gives rise to divergences. We denote the largest value of $\alpha$ leading to a divergence as $\alpha_0$. If no value of $\alpha$ leads to a divergence, then $\alpha_0 = 0$.

Now take $\alpha_L$ to be the minimum value of $\alpha$ such that $\min_j \mathcal{R}e\, \lambda_j(\alpha_L I + C) = 0$ (as seen in Sec. 1.1, this critical value is important for local stability); for any value $\alpha > \alpha_L$ the determinant $|Q(\alpha)|$ is positive, and thus the existence of a solution is guaranteed. This can be easily proven by considering that the determinant of a matrix is the product of its eigenvalues; then the eigenvalues of $Q(\alpha)$ for any $\alpha > \alpha_L$ are either real and positive, or complex and multiplied by their conjugate ($\lambda_j \lambda_j^* > 0$), yielding therefore a positive product, and thus determinant. Therefore, any divergence can only appear when $\alpha \leq \alpha_L$, and thus $\alpha_0 \leq \alpha_L$.

Having considered the denominator, we turn to the numerator $|Q_i(\alpha)|$: whenever this determinant is zero, $y_i^\star(\alpha) = 0$, and thus this component of the solution transitions from negative to positive, or vice versa (because $|Q_i(\alpha)|$ is a polynomial in $\alpha$ and, as such, is a continuous function everywhere). We can characterize this determinant by performing some simplifications. The derivation is easier to write when the focal population $i = n$ is in last position—this can be accomplished for any $i$ by reordering the rows and columns of the matrix. We have (for $i = n$):

$$|Q_i(\alpha)| = \begin{vmatrix} \alpha + C_{1,1} & C_{1,2} & \cdots & 1 \\ C_{2,1} & \alpha + C_{2,2} & \cdots & 1 \\ \vdots & \vdots & & \vdots \\ C_{n,1} & C_{n,2} & \cdots & 1 \end{vmatrix} = \begin{vmatrix} \alpha \widetilde{I} + \widetilde{C} & \mathbb{1} \\ c^T & 1 \end{vmatrix} ,$$

where $\widetilde{I}$ and $\widetilde{C}$ are the corresponding matrices with the last row and column removed, and $c^T$ is the last row of $C$ with the last element removed. We can subtract the last row from all other rows without altering the determinant:

$$\begin{vmatrix} \alpha \widetilde{I} + \widetilde{C} & \mathbb{1} \\ c^T & 1 \end{vmatrix} = \begin{vmatrix} \alpha \widetilde{I} + \widetilde{C} - \mathbb{1} c^T & \mathbb{0} \\ c^T & 1 \end{vmatrix} ,$$

which is a block-structured matrix, with determinant $|\alpha \widetilde{I} + \widetilde{C} - \mathbb{1} c^T|$. Finally, we note that $\tilde{C} - \mathbb{1} c^T = \tilde{B} - \mathbb{1} \tilde{m}^T - (\mathbb{1} b^T - \mathbb{1} \tilde{m}^T) = \tilde{B} - \mathbb{1} b^T$, and thus we can work either with matrix $C$ or $B$ (naturally, if $y_i^\star(\alpha) = 0$, necessarily $x_i^\star(\alpha) = 0$).

This means that $y_i^\star(\alpha) = 0$ whenever $\alpha \geq 0$ matches precisely a real, negative eigenvalue of $\tilde{B} - \mathbb{1} b^T$. Consider the eigenvalues of $\tilde{B} - \mathbb{1} b^T$, and call $\lambda_i$ the most negative, real eigenvalue (if it exists); then, if $-\lambda_i > \alpha_0$, at $\alpha_i = -\lambda_i$ necessarily $y_i^\star(\alpha)$ crosses from negative to positive as $\alpha$ increases (because we have established that when $\alpha \geq \alpha_F$, $y_i^\star(\alpha)$ must be positive).

Now repeat the procedure for all $i$, and collect all the $\alpha_i > \alpha_0$ representing the last transition to positive values for each $i$ (with $\alpha_i = 0$ if such a transition does not exist, or it appears before $\alpha_0$); then necessarily for any $\alpha > \alpha_F = \max\{\alpha_0, \alpha_1, \ldots, \alpha_n\}$ the solution $y^\star(\alpha)$ is positive, and further increasing $\alpha$ maintains positivity.

To summarise these results, we analyze a simple example, showing the location of $\alpha_0$ and $\alpha_i$ ($i = 1, \ldots, n$). Take the matrix $C$:

$$C = \begin{pmatrix} -4 & 2 & 3 \\ -4 & -4 & 2 \\ 8 & 2 & -5 \end{pmatrix} ,$$

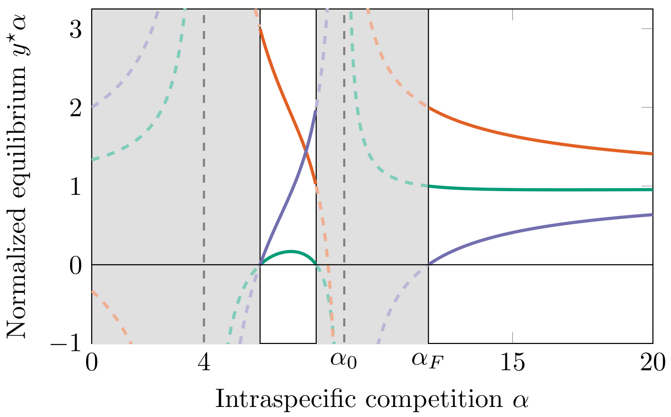


**Supplementary Figure 2**: The components of $y^\star(\alpha)$ as a function of $\alpha \geq 0$. Grey areas denote the values of $\alpha$ for which the equilibrium is not feasible, and the white regions correspond to values of $\alpha$ for which all components $y_i^\star(\alpha)$ (colors) are positive. As explained in the text, for two particular values of $\alpha = 4$ and $\alpha = 9$ (vertical dashed lines), the components of the solution diverge to $\pm\infty$, and thus feasibility is precluded around these points. We denote the largest value of $\alpha$ that leads to a divergence as $\alpha_0$. After the last divergence, the component associated with population 3 (purple) is still negative, and transitions to a positive value at exactly $\alpha_3 = \alpha_F = 12$. The solution remains positive for any $\alpha > \alpha_F$.

with eigenvalues $\lambda(C) = \{-9, -4, 0\}$. Then, $Q(\alpha)$ is singular when $\alpha \in \{4, 9\}$; at these points the three possible $Q_i(\alpha)$ are not singular, and therefore the solution $y^\star(\alpha)$ diverges around these points; we thus set $\alpha_0 = 9$.

The eigenvalues of $\widetilde{C}^{(1)} - \mathbb{1}(c^{(1)})^T$:

$$\widetilde{C}^{(1)} - \mathbb{1}(c^{(1)})^T = \begin{pmatrix} -4 & 2 \\ 2 & -5 \end{pmatrix} - \begin{pmatrix} 2 & 3 \\ 2 & 3 \end{pmatrix} = \begin{pmatrix} -6 & -1 \\ 0 & -8 \end{pmatrix}$$

are $-6$ and $-8$ and thus $y_1^\star(\alpha) = 0$ when we choose $\alpha = 6$ or $\alpha = 8$. Both transitions appear before the last divergence at $\alpha_0 = 9$, and hence the equilibrium of population 1 will be positive after the last divergence (and we set $\alpha_1 = 0$).

Repeating for population two, we have:

$$\widetilde{C}^{(2)} - \mathbb{1}(c^{(2)})^T = \begin{pmatrix} -4 & 3 \\ 8 & -5 \end{pmatrix} - \begin{pmatrix} -4 & 2 \\ -4 & 2 \end{pmatrix} = \begin{pmatrix} 0 & 1 \\ 12 & -7 \end{pmatrix} \quad ,$$

with eigenvalues $-\frac{1}{2}(7 \pm \sqrt{97})$; thus the equilibrium for population 2 will transition only at $\alpha = \frac{1}{2}(7 + \sqrt{97})$, which is smaller than 9; hence the solution for the second population will be positive after the last divergence (and we set $\alpha_2 = 0$ as well).

Finally, considering population 3, we have:

$$\widetilde{C}^{(3)} - \mathbb{1}(c^{(3)})^T = \begin{pmatrix} -4 & 2 \\ -4 & -4 \end{pmatrix} - \begin{pmatrix} 8 & 2 \\ 8 & 2 \end{pmatrix} = \begin{pmatrix} -12 & 0 \\ -12 & -6 \end{pmatrix} \quad ,$$

yielding a transition at $\alpha = 6$ and $\alpha = \alpha_3 = 12$.

Therefore, $\alpha_F = \max\{9, 0, 0, 12\} = 12$. Thus, for this system, all components of $y^\star$ will be positive for any $\alpha > 12$. This behavior is illustrated in Figure 2.

#### 2.4 Finding $\alpha_F$ via eigendecomposition

The results in the previous section are sufficient to devise an exact algorithm to determine $\alpha_F$ for a given system specified by the matrix $C$ (or $B$). It consists of three steps:

1. Compute the eigenvalues of $-C$, and store the largest positive eigenvalue leading to a divergence in $\alpha_0$; if none exist, $\alpha_0 = 0$.
2. For each $i$ in $1, \ldots, n$:
   (a) Form the matrix $\widetilde{C}^{(i)}$ (by removing the $i^{\text{th}}$ row and column from $C$) and the corresponding $(c^{(i)})^T$ (i.e., the $i^{\text{th}}$ row of $C$ with the $i^{\text{th}}$ coefficient removed);
   (b) Compute the eigenvalues of the matrix $\mathbb{1}(c^{(i)})^T - \widetilde{C}^{(i)}$;
   (c) Store the largest positive, real eigenvalue in $\alpha_i$; if no eigenvalues are real and positive, or if all are smaller than $\alpha_0$, then $\alpha_i = 0$.
3. $\alpha_F = \max\{\alpha_0, \alpha_1, \ldots, \alpha_n\}$.

From a computational standpoint, we note that the eigenvalue with the largest real part can be computed efficiently, for example via the Arnoldi iteration[11]. One could also utilize well-known bounds on eigenvalues to compute $\alpha_i$ only for the matrices that are more likely to produce a maximum.

For the example above, we find $\alpha_0 = 9$, $\alpha_1 = \alpha_2 = 0$, and $\alpha_3 = 12 = \alpha_F$.

A similar pattern can also be observed in the classic two-species Lotka-Volterra competition model with priority effects (Fig. 3). Mirroring the example in the main text, Phase I is characterized by *locally* stable boundary equilibria – alternative stable states. Global stability of an unspecified boundary equilibrium (in this case, that where only species $x_2$ is present) is achieved first (Phase II), and as $\alpha$ keeps increasing, feasibility is achieved at last (Phase III), such that the two species coexist at a globally stable equilibrium.

### Supplementary Note 3. Random systems

Having derived some general results in the previous section, we now turn to the case in which the matrix of interactions $B$ is a large random matrix. This case has been studied for decades, producing a wealth of results. Here we show that previous results can be extended, generalized and refined in light of the derivations above. This section has two main goals: a) compute the probability that the equilibrium $x^\star$ of Eq. (1) is positive, given a value of $\alpha$ and the statistics of the random matrix $B$; b) show how these derivations have profound implications for the type of dynamics that can be observed for these systems.

#### 3.1 Random systems with statistically equivalent populations

We start by defining what we mean by random systems. Naturally, dynamical systems in which some (all) of the parameters are sampled from random distributions are indeed random systems; here we focus on a subclass in which the model definition is closed under relabeling of the populations: by permuting the indices $x_i$ of Eq. (1) we obtain the same random model. In practice, this simply means that there is no special “structure” making any of the populations behave differently from the others—the populations are *statistically equivalent*.

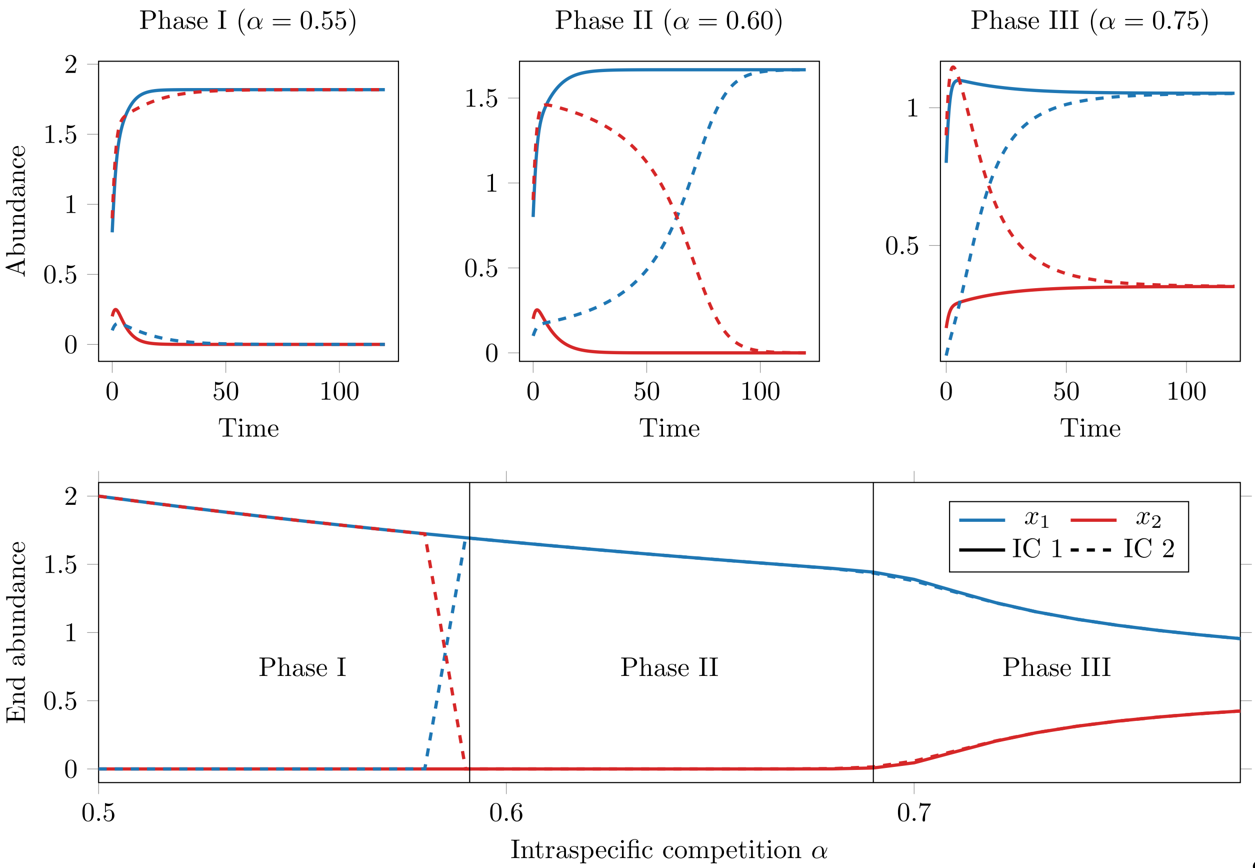


d

**Supplementary Figure 3**: Three dynamical phases in the classic two–species Lotka–Volterra competition model. Top row: time series for $\alpha \in \{\alpha_{\mathrm{I}}, \alpha_{\mathrm{II}}, \alpha_{\mathrm{III}}\}$ (here, $0.55, 0.60, 0.75$), with two initial conditions shown (linetypes; colors denote species). Bottom row: bifurcation diagram of equilibrium abundances versus the intraspecific competition strength $\alpha$ ($x$–axis), with line type indicating the initial condition (IC1 or IC2). As $\alpha$ increases (while $a_{12}, a_{21}$ are fixed and $r_1 = r_2 = 1$), the system transitions through three regimes separated by the vertical guides: **Phase I (low $\alpha$)**: interspecific effects dominate and priority effects appear—different initial conditions lead to alternative stable outcomes (each species can exclude the other). **Phase II (intermediate $\alpha$)**: competitive exclusion is deterministic—the same species survives from any initial condition (unique attracting boundary equilibrium). **Phase III (high $\alpha$)**: intraspecific competition dominates and a feasible interior coexistence equilibrium exists and attracts all initial conditions. All three panels use the same interspecific coefficients, varying only $\alpha$ across scenarios.

Among this subclass of models, the simplest to study is one in which the matrix of interactions is given by the sum of a random matrix $B$ and a positive constant $\mu$.

$$B' = B + \mu \mathbb{1}\mathbb{1}^T \quad ,$$

where we choose $B$ such that: $B_{ii} = 0$ (this assumption can be relaxed, Sec. 3.9), and $\mathbb{E}(B_{ij}) = 0$, $\mathbb{E}(B_{ij}^2) = 1$, $\mathbb{E}(B_{ij}B_{ji}) = \rho$ for every $i \neq j$. Thus the spectrum of $B$ follows the elliptic law[12]: the eigenvalues of $B/\sqrt{n}$ are approximately uniformly distributed in an ellipse in the complex plane, with center $(0, 0i)$, horizontal semi-axis $1 + \rho$, and vertical semi-axis $1 - \rho$.

As we have shown in Section 2.1, as long as the columns of $B'$ sum to positive values (i.e., when the systems are competitive), the value of $\mu$ has no effect on feasibility. The value of $\mu$ also has no effect on the stability of the system[13]: the eigenvalues of $B'$ will be distributed in an ellipse, possibly with an outlier eigenvalue on the right of the bulk, and thus not influencing stability. Hence, $\alpha_F$ (the value at which the equilibrium is feasible, and feasibility is maintained when we increase $\alpha$), $\alpha_L$ (the value at which all eigenvalues have positive real part), and $\alpha_S$ (the value at which the symmetric part of the matrix is positive definite) are the same for both $B$ and $B'$.

Thus, without loss of generality, we can consider the case in which $\mu = 0$, and therefore the matrix of interactions follows the elliptic law. This also allows us to compare our results directly with other studies that have considered random matrices with coefficients sampled from distributions with mean zero[14–16].

##### 3.2 Previous results

The oldest result in this area goes back to a response to May's article, by Alan Roberts[17]. Roberts used computer simulations to measure the probability that a feasible equilibrium would be locally stable for a GLV model, Eq. (1), in which populations were self–regulating ($A_{ii} = 1$), and interactions were randomly chosen to be $A_{ij} = \theta$ with probability $1/2$ and $A_{ij} = -\theta$ otherwise. He found that the probability of stability conditional on feasibility was very large, and increasing with the size of the system.

The next main contribution was obtained by Lewi Stone as part of his PhD thesis[14], developed under the supervision of Roberts. Using techniques similar to those used here, Stone was able to approximate the probability of feasibility for matrices with random interactions sampled independently from an arbitrary distribution with mean zero. For example, a formula reported in Stone[18], and adapted to the notation employed in this work, reads:

$$p_F(\alpha) = \Phi\left(\frac{\alpha}{\sqrt{n-1}(1+\frac{n-1}{\alpha^2})^{\frac{1}{4}}}\right)^n \quad ,$$

where $p_F(\alpha) = P(x^\star > \mathbb{0}|\alpha)$, and $\Phi(\cdot)$ is the cumulative distribution function for the standard normal distribution. This formula was derived in[14] by approximating the solution of the linear system using the Neumann series (see Sec. 3.7). Stone's thesis also reported larger computer simulations, in the spirit of Roberts, again confirming that feasible systems are almost invariably locally stable.

More recently, two articles[15, 16] used different techniques to show that there exist a critical value below which the probability of feasibility is negligible, and above which it is very high. In particular, using the notation adopted in this work, they found that the probability of feasibility is close to one for $\alpha \gg \sqrt{n}\sqrt{2\log n}$, and is close to zero for $\alpha \ll \sqrt{n}\sqrt{2\log n}$. Clenet *et al.*[16] considered matrices with correlated entries as well, finding that the threshold value for feasibility would not change substantially, and, importantly, noting that the transition would appear after that for global stability in these GLV systems. Also important for the work developed here, Liu *et al.*[19] showed, using yet a different technique, that correlation between the entries would have an effect on feasibility (increasing the probability of feasibility when the correlation is negative), but that the effect would be negligible for large systems.

Finally, Marcus *et al.*[20] considered stable, feasible GLV systems and increased the interaction strengths (equivalently, one could have decreased the strength of intraspecific interactions, as done here) until either the system becomes unfeasible, or unstable. They found that in general, the system loses feasibility before

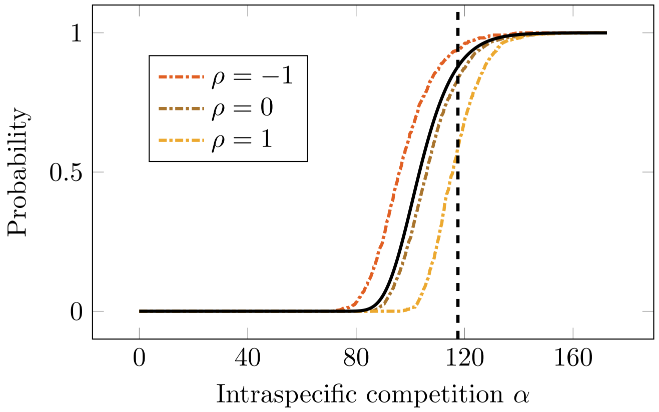


**Supplementary Figure 4**: Probability of feasibility (i.e., $P(x^\star > \mathbb{0})$, y-axis) for a given level of intraspecific interaction $\alpha$ (x-axis). The probability was computed by considering 1000 realizations of systems of size $n = 1000$ with interaction matrices $B$ following the elliptic law (with the colors representing the off-diagonal correlation). The black line represents the approximation by Stone[18] for the case $\rho = 0$. The vertical dashed line is the critical value of Bizeul & Najim[15].

stability, consistently with the previous findings outlined above. Interestingly for our argument, they notice that in symmetric systems, feasibility would be often lost due to the components of the solution diverging to $\pm\infty$, a fact that will be key for the developments below. A similar argument was put forward by Stone[18].

Figure 4 shows simulations measuring the probability of feasibility when varying both $\alpha$ (x-axis) and the correlation between entries (colors); the approximation by Stone[14] for $\rho = 0$, as well as the critical value by Bizeul & Najim[15] are also reported.

The goal of the next few sections is to generalize and improve these results in the light of what we have derived for general matrices. In particular, we will extend both the formula by Stone and the critical value of Bizeul & Najim for the case in which entries are correlated.

##### 3.3 Feasibility when $\alpha = 0$

For large random systems, the probability of feasibility when $\alpha = 0$ is negligible. We can compute this probability exactly whenever the coefficients of $B$ are sampled independently from a distribution that is symmetric about 0.

We want to compute the probability that $(\alpha I + C)y^\star = \mathbb{1}$ has a positive solution when $\alpha = 0$. Since the columns of $C$ sum to 0 (see Section 2.1), the right-hand side of the equation vanishes when we multiply the equations by the transpose of $C$:

$$\begin{aligned} Cy^\star &= \mathbb{1} \\ C^T C y^\star &= C^T \mathbb{1} \\ C^T C y^\star &= \mathbb{0} \quad . \end{aligned}$$

The matrix $C^T C$ is symmetric, positive semi-definite, and has an eigenvalue 0; all its eigenvectors are orthogonal. Then, whenever a feasible solution exists, solving the system of equations above is equivalent to solving $Cy^\star = \mathbb{0}$, i.e., $y^\star$ is a right eigenvector of $C$ associated with the zero eigenvalue. Thus, $y^\star$ is defined up to a constant of proportionality, allowing us to set one of the components arbitrarily. Here we set $y^\star_n = 1$, and we partition $C$ and $y^\star$ as:

$$C = \begin{pmatrix} \widetilde{C} & a \\ c^T & b \end{pmatrix} \quad y^\star = \begin{pmatrix} \widetilde{y} \\ 1 \end{pmatrix} \quad ,$$

where $b = -\sum_{j=1}^{n-1} a_j$. We can therefore write $n-1$ independent equations:

$$\begin{aligned} \widetilde{C}\widetilde{y} &= -a \\ \widetilde{y} &= -\widetilde{C}^{-1}a \quad . \end{aligned}$$

For feasibility, we need all $\widetilde{y} > \mathbb{0}$ (because we have chosen a positive $y^\star_n$). To compute the probability that $\widetilde{y} > \mathbb{0}$ exactly, we exploit the assumptions that we have made: because of symmetry, the probability of observing a given $C_{ij}$ (or the density of the distribution at $C_{ij}$, when the distribution is continuous), satisfies the equality $P(C_{ij}) = P(-C_{ij})$.

Then, we can follow the approach of Serván *et al.*[21], and define the diagonal matrices $K_i = D(s_i)$, where the $s_i$ are the $2^n$ distinct vectors of length $n$ containing either 1 or $-1$: $s_1 = (1, 1, 1, \ldots, 1)^T$, $s_2 = (1, -1, 1, \ldots, 1)^T$, $s_3 = (-1, -1, 1, \ldots, 1)^T$, $s_4 = (1, 1, -1, \ldots, 1)^T$, $\ldots$, $s_{2^n} = (-1, -1, -1, \ldots, -1)^T$. We partition these matrices as well:

$$K_i = \begin{pmatrix} \widetilde{K_i} & \mathbb{0} \\ \mathbb{0}^T & \sigma_i \end{pmatrix} \quad .$$

Because of symmetry and independence, we have $P(C) = P(CK_i)$, given that multiplying $C$ by $K_i = D(s_i)$ simply changes the signs of all the coefficients in the columns corresponding to negative signs in $s_i$. Finally, we write:

$$\begin{aligned} \widetilde{C}\widetilde{K}_1\widetilde{y} &= -\sigma_1 a \\ \widetilde{y} &= -\sigma_1 \widetilde{K}_1 \widetilde{C}^{-1} a \quad , \end{aligned}$$

where we have used the fact $K_1 = I$. Now notice that by exchanging $K_1$ with $K_i$, we can change the signs of the components of $\widetilde{y}$; then, for each possible pattern of signs of $-\widetilde{C}^{-1}a$ there are exactly two choices of $K_i$ that make the vector $\widetilde{y}$ all positive (one solution with $K_j = D(s_j)$, and the other with $K_l = D(-s_j)$). Given that $P(CK_i) = P(CK_j)$, and that there are $2^n$ matrices $K_i$, the probability of feasibility is exactly $2/2^n = 1/2^{n-1}$.

Thus, for a random system with $B_{ij}$ sampled independently from a distribution that is symmetric about its mean, for two populations we have fifty percent probability of obtaining a feasible equilibrium, and the probability rapidly decreases, so that we have less than one in a thousand chance to find a feasible equilibrium for $n = 11$ and less than one in a million for $n = 21$.

##### 3.4 Feasibility when $\alpha < \alpha_L$

We have shown above that the probability of feasibility is vanishingly small for $\alpha = 0$; here we show that the same holds true for any $\alpha < \alpha_L$. In this case, the support for the bulk of the spectrum of $\alpha I + B$ contains the point $(0, 0i)$, because the ellipse is not completely contained in the right half of the complex plane. Because the matrix is large, and eigenvalues are approximately uniformly distributed inside the ellipse, we can always find an eigenvalue that is arbitrarily close to the origin, thereby causing some solutions to reach arbitrarily large absolute values—given that the determinant of matrix $Q(\alpha) = \alpha I + B$ (defined in Sec. 2.3) must be arbitrarily close to zero. Because the mean of $y^\star$ is positive, components reaching large positive values must be balanced by some other components with negative values, and thus feasibility is precluded. Hence, the probability of feasibility at $\alpha < \alpha_L$ is close to zero. A similar argument is found (in the context of symmetric matrices), in Marcus *et al.*[20]. Therefore, we can assume that $\alpha_0 = \alpha_L$.

##### 3.5 A single transition to feasibility when $\alpha > \alpha_L$

Having shown that, with high probability, the system is not feasible for $\alpha < \alpha_L$, we consider the dependence of each component of $y^\star(\alpha)$ on $\alpha$, as it increases for $\alpha > \alpha_L$. We have seen in Section 2.3 that $y_i^\star(\alpha)$ transitions (either from positive to negative, or vice versa) whenever $\alpha = \lambda_i$, and $\lambda_i$ is a real, positive eigenvalue of the matrix

$$\widetilde{Q}_i = \mathbb{1}(b^{(i)})^T - \widetilde{B}^{(i)} \quad ,$$

where $\widetilde{B}^{(i)}$ is the matrix $B$ with the row and column corresponding to $i$ removed, and $(b^{(i)})^T$ is the $i^{\text{th}}$ row of $B$ with coefficient $i$ removed. If $\widetilde{B}^{(i)}$ is a large random matrix (i.e., if $B$ is a large random matrix following the elliptic law), then perturbing it by adding a rank-one matrix has a simple effect on its spectrum: at most one eigenvalue moves outside of the bulk, while the remaining eigenvalues are fundamentally unchanged[22] (an example is provided in Fig. 5). Thus, if the outlier eigenvalue is on the left of the bulk, then $\lambda_i$ is negative and $y_i^\star(\alpha)$ cannot cross zero when choosing $\alpha > 0$. If, on the other hand, the outlier eigenvalue $\lambda_i > \alpha_L$ is positive, we can make $y_i^\star(\alpha) = 0$ by choosing $\alpha = \lambda_i$. Then, this transition must bring $y_i^\star(\alpha)$ from negative to positive values: because we know that, for sufficiently large values of $\alpha$, $y_i^\star(\alpha) > 0$, and that there are no divergences after $\alpha_L$, then necessarily $y_i^\star(\alpha)$ becomes zero at this value, and remains positive when $\alpha$ is increased further (observe that $y_i^\star(\alpha)$ is a rational function —a quotient of two polynomials— in $\alpha$, which is a continuous function above the largest point at which it diverge, i.e., for $\alpha > \alpha_0$; therefore, the change of sign follows by continuity of $y_i^\star(\alpha)$ for $\alpha > \alpha_L$). Hence, we can set $\alpha_i = \lambda_i$.

We have just shown that each matrix $Q_i(\alpha)$ has at most a single real eigenvalue to the right of the bulk, with value $\lambda_i = \alpha_i$. Moreover, when this positive outlier eigenvalue exists, component $i$ transitions from negative to positive at $\alpha = \alpha_i$. But then we can order the $\alpha_i$ from smallest to largest, and at $\alpha_F = \max_i \alpha_i$, all components have crossed into positive values. Because for large random matrices feasibility is precluded (with very high probability) when $\alpha < \alpha_L$, we conclude that a) positivity is gained only at $\alpha_F$, and b) it is never

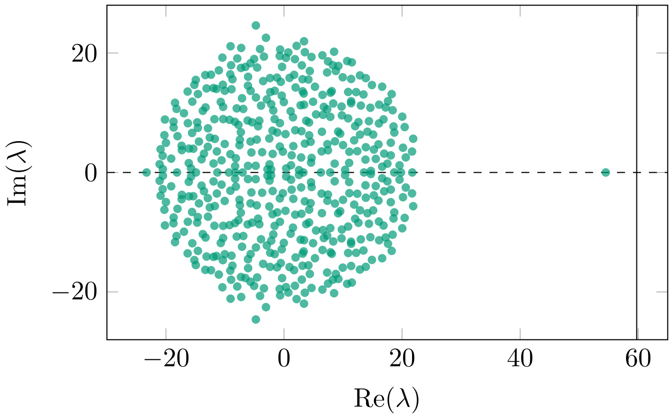


**Supplementary Figure 5**: Spectrum of $\widetilde{Q}_i = \mathbb{1}(b^{(i)})^T - \widetilde{B}^{(i)}$ for a given $i$, when $B$ is a large random matrix ($n = 1000$) following the circular law. When a large random matrix ($-\widetilde{B}^{(i)}$ in this case) is perturbed by summing a matrix with small rank (in this case, matrix $\mathbb{1}(b^{(i)})^T$ has rank 1), only a few of the original eigenvalues change substantially (one in the case of a rank-one matrix). Moreover, the eigenvalues are expected to be close to those of the small-rank matrix (vertical line).

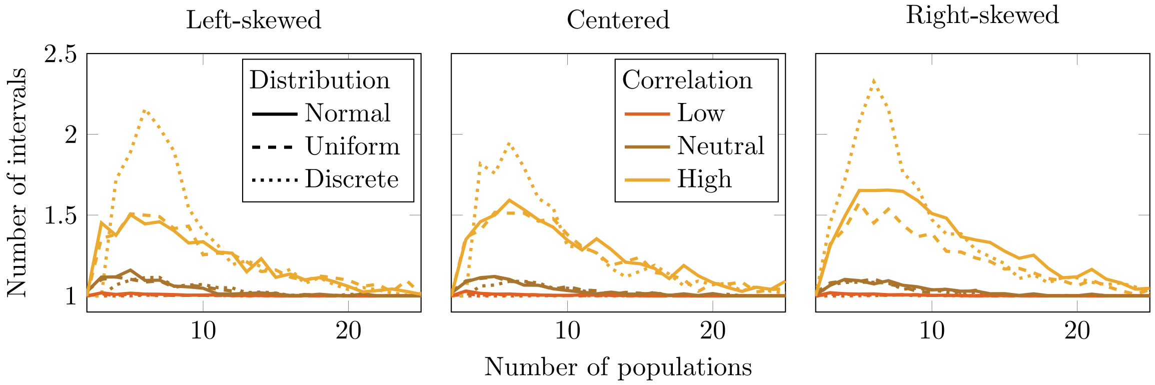


**Supplementary Figure 6**: Number of disjoint intervals yielding a feasible equilibrium converges to one for several distributions, skewness, and correlation values. For random systems of increasing size ($n$, x-axis), we count the number of disjoint intervals for $\alpha$ yielding a feasible equilibrium. As we increase $n$, the number of intervals converges to one, irrespective of correlation $\rho$ or choice of distribution. Then, large random matrices will yield a single transition, occurring at $\alpha_F$.

lost for larger values of $\alpha$. Thus, large random matrices produce a much simpler picture of the dependence on $\alpha$ than small, or arbitrary matrices, in which, when we grow $\alpha$, the equilibrium can transition from unfeasible to feasible several times (Sec. 2.3). These findings are confirmed by simulations: when we count the number of disjoint intervals for $\alpha$ yielding a feasile equilibrium for random systems of increasing size, we see a rapid convergence to a single interval (Fig. 6).

##### 3.6 Approximating the probability of feasibility using the theory of small-rank perturbations of random matrices

We can use the argument above on the perturbation of random matrices with small-rank matrices to provide an estimate of the probability of feasibility. We write $P(x^\star > \mathbb{0}|\alpha) = p_F(\alpha)$. In light of the previous paragraph, this amounts to $P(\lambda_i < \alpha)$ for all $n$ choices of $i$, where $\lambda_i$ is either the largest real eigenvalue of $\mathbb{1}(b^{(i)})^T - \widetilde{B}^{(i)}$, when it exists, or $\lambda_i = 0$, when either the matrix has no positive eigenvalues, or all are smaller than $\alpha_L$.

To approximate the location of $\lambda_i$, we follow O'Rourke and Renfrew[22], who have proven that, when $A$ is a large random matrix following the elliptic law, $uv^T$ is a rank-one matrix with real eigenvalue $\gamma$, and $|\gamma| > 1 + \rho$, then the spectrum of $A/\sqrt{n} + uv^T$ has an outlier, real eigenvalue, $\lambda_o$, with:

$$\lambda_o = \gamma + \frac{\rho}{\gamma} + o(1) \quad .$$

Let $\lambda_i$ be the outlier of matrix $\mathbb{1}(b^{(i)})^T - \widetilde{B}^{(i)}$. We want to find its location. Define $\gamma_i = \sum_j b_j^{(i)}$, i.e., the only nonzero eigenvalue of $\mathbb{1}(b^{(i)})^T$. Then, the outlier of $\frac{1}{\sqrt{n-1}}(\mathbb{1}(b^{(i)})^T - \widetilde{B}^{(i)})$ is located approximately at

$$\frac{\lambda_i}{\sqrt{n-1}} = \frac{\gamma_i}{\sqrt{n-1}} + \frac{\rho\sqrt{n-1}}{\gamma_i} \quad , \tag{4}$$

if $\left|\frac{\gamma_i}{\sqrt{n-1}}\right| > 1 + \rho$ holds. Otherwise, the outlier does not exists because it is contained in the bulk. For $\gamma_i > \sqrt{n-1}$, the right-hand side of Eq. (4) is maximized when $\gamma_i$ is maximized, for any value of $\rho$; hence, the maximum $\lambda_i = \hat{\lambda}$ tends to be associated with the largest $\gamma_i = \hat{\gamma}$, which in turn is simply the maximum row sum of $B$.

Now we assume that the $\gamma_i$s are independent and normally distributed, with mean zero and variance $n-1$; an assumption that is correct when $\rho = 0$ and $n$ is sufficiently large, and is met only approximately for nonzero values of $\rho$ (though the effect of the correlation on row sums must be negligible for large $n$). Thus, the variables $\tilde{\gamma}_i = \gamma_i/\sqrt{n-1}$ follow the standard normal distribution. The distribution of the maximum $\tilde{\gamma}$ therefore converges asymptotically (albeit quite slowly[23]) to the Gumbel distribution[15]. In particular, asymptotically we have:

$$P(\tilde{\gamma} < x) = \exp\left(-\exp\left(-((x - b_n)b_n\right)\right) \quad ,$$

where $b_n$ is a constant that can be chosen to guarantee a faster convergence to the Gumbel distribution. Here we follow Hall[23], and use $b_n = \sqrt{\mathcal{W}_0\left(\frac{n^2}{2\pi}\right)}$, where $\mathcal{W}_0(z)$ is the principal branch of the Lambert function. One can also Taylor-expand the function for large $n$, obtaining:

$$\mathcal{W}_0\left(\frac{n^2}{2\pi}\right) \approx \log\frac{n^2}{2\pi} - \log\log\frac{n^2}{2\pi} + \frac{\log\log\frac{n^2}{2\pi}}{\log\frac{n^2}{2\pi}} + \cdots \quad .$$

We want to derive an approximate distribution for $\alpha_F = \sqrt{n-1}\,\tilde{\lambda}$, where, according to the small-rank perturbations, we have

$$\tilde{\lambda} \approx \tilde{\gamma} + \frac{\rho}{\tilde{\gamma}} \quad .$$

We now solve this equation for $\tilde{\gamma}$, obtaining:

$$\tilde{\gamma} \approx \frac{\tilde{\lambda} + \sqrt{\tilde{\lambda}^2 - 4\rho}}{2} \quad .$$

Substituting this value in the definition of the Gumbel distribution above, we have:

$$P(\tilde{\lambda} < x) = \exp\left(-\exp\left(-\left(\frac{x + \sqrt{x^2 - 4\rho}}{2} - b_n\right) b_n\right)\right) \quad .$$

Concluding, the probability that the system is feasible at $\alpha$ is the probability that $\alpha_F < \alpha$, and thus $\alpha_F/\sqrt{n-1} = \tilde{\lambda} < \alpha/\sqrt{n-1}$:

$$p_F(\alpha) = P\left(\frac{\alpha_F}{\sqrt{n-1}} < \frac{\alpha}{\sqrt{n-1}}\right) = P(\tilde{\lambda} < \tilde{\alpha}) = \exp\left(-\exp\left(-\left(\frac{\tilde{\alpha} + \sqrt{\tilde{\alpha}^2 - 4\rho}}{2} - b_n\right) b_n\right)\right) \quad , \tag{5}$$

where $\tilde{\alpha} = \alpha/\sqrt{n-1}$.

The Gumbel cumulative density function $\exp(-\exp(-x))$ describes a random variable with mode 0, median $-\log\log 2 \approx 0.366$ and mean $\approx 0.57721$ (the Euler-Mascheroni constant). Thus, we expect the $\alpha_F$ for a given $n$ and $\rho$ to have the following statistics:

***Mode of $\alpha_F$.***

We solve:

$$\begin{aligned}
\left(\frac{\tilde{\alpha} + \sqrt{\tilde{\alpha}^2 - 4\rho}}{2} - b_n\right) b_n &= 0 \\
\tilde{\alpha} &= b_n + \frac{\rho}{b_n} \\
\alpha &= \sqrt{n-1}\left(b_n + \frac{\rho}{b_n}\right) \quad .
\end{aligned}$$

Choosing $b_n \approx \sqrt{2\log n}$ and $\rho = 0$, we recover the critical value of Bizeul & Najim[15].

***Median and mean of $\alpha_F$***

We now solve for $x = k$, obtaining:

$$\begin{aligned}
\left(\frac{\tilde{\alpha} + \sqrt{\tilde{\alpha}^2 - 4\rho}}{2} - b_n\right) b_n &= k \\
\tilde{\alpha} &= b_n + \frac{k}{b_n}\frac{b_n\rho}{b_n^2 + k} \\
\alpha &= \sqrt{n-1}\left(b_n + \frac{k}{b_n}\frac{b_n\rho}{b_n^2 + k}\right) \quad .
\end{aligned}$$

Choosing $k = -\log\log 2$ we obtain the approximate median, and using the Euler-Mascheroni constant, the mean.

While Eq. (5) is relatively simple and appealing, it tends to overestimate the probability of feasibility (i.e., underestimate the magnitude of the maximum $\lambda_i$) for low $\alpha$, even when the matrices have considerable size. In fact, the equation is derived under the assumption that the maximum $\hat{\lambda} = \lambda_i$ is associated with the index $i$ corresponding to the row with maximum row sum. While this is often the case, it is not universally true: the exact location of $\lambda_i$ depends in fact on all of the eigenvalues and eigenvectors of both $\widetilde{B}^{(i)}$ and $\mathbb{1}(b^{(i)})^T$—accordingly, the theorem by O'Rourke and Renfrew[22] contains an unknown term of order one, describing

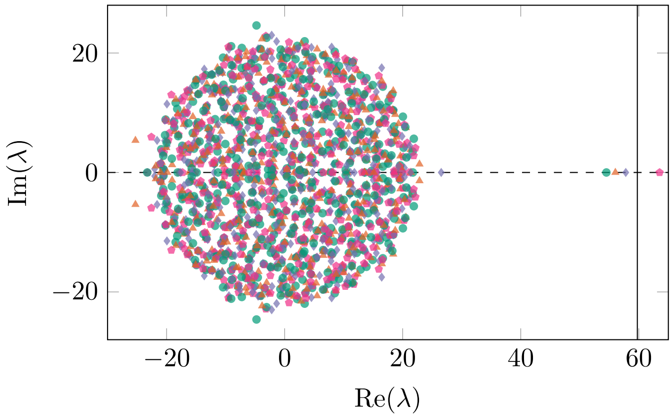


**Supplementary Figure 7**: Spectra of several $Q_i = \mathbb{1}(b^{(i)})^T - \widetilde{B}^{(i)}$ matrices. In green we have the spectrum of the matrix $Q_i$ for the $i$ associated with the largest row sum of $B$ (i.e., $\gamma_i = \hat{\gamma}$), in orange that for the $i$ associated with the second largest row sum, in purple that for the third largest row sum, and in pink that for the fourth largest row sum. Even though we would expect the outlier associated with the largest row sum to be the largest (and thus $\hat{\lambda} \approx \hat{\gamma}$, given that $\rho = 0$), we find that in this case the largest outlier is that associated with the fourth largest row sum. Because of this, we underestimate the value of $\alpha_F$ (vertical line).

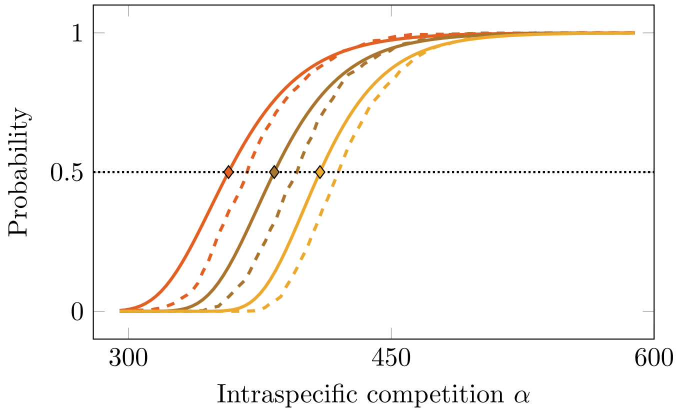


**Supplementary Figure 8**: Simulation results (dashed lines, colors representing the off-diagonal correlation, $n = 10000$), vs. the approximation in Eq. (5) for the probability of feasibility (y-axis) when $\alpha$ is varied (x-axis). The diamonds mark the median of the approximate distribution—i.e., the value of $\alpha$ for which the approximation returns $p_F = 1/2$. Note that the approximation captures the spacing caused by the correlation, but tends to overestimate the probability of feasibility, especially when $\alpha$ is low.

the fluctuations in the outlier eigenvalue. Now, consider the index $j$ associated with the row with the second largest row sum; for a large random matrix, $\gamma_j$ will be extremely close to $\gamma_i$, and the corresponding $\lambda_j$ might well be lager than $\lambda_i$, because of the fluctuations captured in the term in $o(1)$. The same argument can be made for the third-largest $\gamma_k$, and so forth, at least up to a certain proportion of the $\gamma_k$. Overall, this means that the estimate of $\hat{\lambda}$ tends to be conservative (especially for low $\alpha$), and that therefore the probability of feasibility is typically overestimated in this region. Fig. 7 exemplifies this issue.

Despite these limitations, Eq. (5) still captures qualitatively the results of simulations: the ordering and spacing of the curves when varying $\rho$ is correct, and the formula, despite its simplicity, describes the shape of the transition to feasibility quite well. It also shows that these curves are *universal*—as long as the matrix $B$ follows the elliptic law, the formula holds asymptotically, irrespective of the fine details of the distribution of $B_{ij}$. Figure 8 shows the approximation along with the results of simulations for random systems with varying correlation parameter $\rho$.

#### 3.7 More accurate approximations of the probability of feasibility

In the section above, we have derived a coarse approximation for the probability of feasibility $p_F(\alpha)$, producing a simple and intuitive equation. The approximation is however inaccurate, and tends to overestimate the probability of feasibility for low values of $\alpha$. To derive a more accurate approximation, we first state a general result: for any random model with statistically equivalent populations, the mean and correlation between the equilibrium values are fixed; the difference between the models is captured by the variance of the equilibrium values. Having established this result, we provide two methods to approximate the variance of the solution's components, one using the theory of iterative solutions of linear systems, and the other using the theory of resolvents.

When populations are statistically equivalent, then some measures of the distribution of the components of $y^\star$ are especially simple to derive. We have already seen that the expectation of $y_i^\star$, $\mathbb{E}(y_i^\star) = 1/\alpha$, and we denote the variance as $\mathbb{V}(y_i^\star) = \mathbb{E}((y_i^\star)^2) - \mathbb{E}(y_i^\star)^2$, the covariance between $y_i^\star$ and $y_j^\star$, $\mathbb{Cov}(y_i^\star, y_j^\star) = \mathbb{E}(y_i^\star y_j^\star) - \mathbb{E}(y_i^\star)\mathbb{E}(y_j^\star)$, and their correlation $\mathbb{Cor}(y_i^\star, y_j^\star)$.

Given that the sum of $y_i^\star$ is exactly $n/\alpha$, then, $(\sum_j y_j^\star)^2 = n^2/\alpha^2$. Expanding, we obtain:

$$\begin{aligned}
\left(\sum_j y_j^\star\right)^2 &= \frac{n^2}{\alpha^2}\\
\sum_j (y_j^\star)^2 + \sum_i \sum_{j \neq i} y_i^\star y_j^\star &= \frac{n^2}{\alpha^2}\\
n\mathbb{E}((y_i^\star)^2) + n(n-1)\mathbb{E}(y_i^\star y_j^\star) &= \frac{n^2}{\alpha^2}\\
\mathbb{V}(y_i^\star) + \mathbb{E}(y_i^\star)^2 + (n-1)\left(\mathbb{Cov}(y_i^\star, y_j^\star) + \mathbb{E}(y_i^\star)\mathbb{E}(y_j^\star)\right) &= n\mathbb{E}(y_i^\star)^2\\
\mathbb{V}(y_i^\star) + \mathbb{E}(y_i^\star)^2 + (n-1)\left(\mathbb{Cov}(y_i^\star, y_j^\star) + \mathbb{E}(y_i^\star)^2\right) &= n\mathbb{E}(y_i^\star)^2\\
\mathbb{V}(y_i^\star) + (n-1)\mathbb{Cov}(y_i^\star, y_j^\star) &= 0 \quad ,
\end{aligned}$$

where we have exploited the fact that, for a model with statistically equivalent populations, $\mathbb{E}(y_i^\star)\mathbb{E}(y_j^\star) = \mathbb{E}(y_i^\star)^2$, and that the covariance $\mathbb{Cov}(y_i^\star, y_j^\star)$ must be the same for all $i, j$. Thus, for *any* unstructured model, we have that the expected covariance is negative. Moreover, for a given variance $\mathbb{V}(y_i^\star) = \sigma^2_{y^\star}$, we have: $\sigma^2_{y^\star} + (n-1)\sigma^2_{y^\star}\mathbb{Cor}(y_i^\star, y_j^\star) = 0$ and thus $\mathbb{Cor}(y_i^\star, y_j^\star) = -1/(n-1)$ for all $i$ and $j$. Note that this is the maximally negative correlation one can have for equicorrelated random variables. Thus, $y^\star$ has expectation $(1/\alpha)\mathbb{1}$, and covariance matrix $\sigma^2_{y^\star}\left(\frac{n}{n-1}I - \frac{1}{n-1}\mathbb{1}\mathbb{1}^T\right)$. Importantly, the only difference between different unstructured models with random parameters is the value of $\sigma^2_{y^\star}$, while the rest is unchanged. Analyzing increasingly complex models therefore reduces to the problem of determining the variance of $y^\star$.

The joint distribution of $y^\star$ is well-approximated by a multivariate normal[14] in the limit of large $n$, because of the central limit theorem. To show this, we expand the solution using the Neumann series (see Section 3.7). Whenever $\alpha > \alpha_L$, the solution can be written as:

$$y^\star = \frac{1}{\alpha}\left(I + \frac{1}{\alpha}C\right)^{-1}\mathbb{1} = \frac{1}{\alpha}\sum_{k=0}^{\infty}\left(-\frac{1}{\alpha}\right)^k C^k \mathbb{1} \quad .$$

It is well known that any vector $b$ follows a multivariate normal distribution if and only if, for an arbitrary vector $w$ of the same dimension, the scalar product $w^T b$ is distributed according to a univariate normal distribution. Then, for an arbitrary vector $w$, we consider the scalar variable

$$\xi = w^T y^\star = \frac{1}{\alpha} \sum_{k=0}^{\infty} \left(-\frac{1}{\alpha}\right)^k w^T C^k \mathbb{1} \quad .$$

We want to show that $\xi$ is normally distributed. Using the spectral decomposition $C = P\Lambda P^{-1}$, where $\Lambda$ stands for the diagonal matrix of eigenvalues, and the columns of $P$ contain eigenvectors, we can write

$$w^T C^k \mathbb{1} = \sum_{\ell=1}^{n} \lambda_\ell^k (P^T w)_\ell (P^{-1}\mathbb{1})_\ell \quad .$$

Then, $w^T C^k \mathbb{1}$ is a sum of $n$ variables that are equally distributed. Thus, by the central limit theorem, in the limit $n \to \infty$, the distribution of $w^T C^k \mathbb{1}$ can be well-approximated by a univariate normal. The normality of $\xi$ follows from the properties of linear combinations of normal random variables. Therefore, $y^\star$ is asymptotically distributed as

$$y^\star \sim \mathcal{N}\left(\frac{1}{\alpha}\mathbb{1}, \sigma_{y^\star}^2 \left(\frac{n}{n-1} I - \frac{1}{n-1}\mathbb{1}\mathbb{1}^T\right)\right) \quad .$$

Because the joint distribution of $y^\star$ is well-approximated by a multivariate normal, we can calculate the probability of feasibility as a function of $\alpha$, $n$, as well as $\sigma_{y^\star}^2$ by integrating the probability density function in the positive orthant of $\mathbb{R}^n$—what is called an "orthant probability":

$$p_F(n, \alpha, \sigma_{y^\star}^2) = \int_0^\infty \int_0^\infty \cdots \int_0^\infty \phi(y^\star) dy_1^\star dy_2^\star \cdots dy_n^\star \quad ,$$

where $\phi(y^\star)$ is the probability density function of the normal distribution above. For equicorrelated random variables, the multiple integral above can be reduced to a single integral (albeit involving complex variables when the correlation is negative, such as in this case[24]). Equivalently, one can use the theory of order statistics to write the density function of the first order statistics (i.e., the $\min_i(y_1^\star, y_2^\star, \ldots, y_n^\star)$) and integrate this density function for all positive values of the minimum.

When $n$ is large, the fact that the components of $y^\star$ are slightly negatively correlated becomes less and less important, and we can treat them approximately as if they were independent. This simplification is common in high-dimensional probability: correlations of order $1/n$ vanish as $n \to \infty$.

Under this independence assumption, and given that the components of $y^\star$ are identically distributed, we can focus on the probability that $y_i^\star > 0$, and $p_F$ is simply this value raised to the $n^{\text{th}}$ power.

Each random variable $y_i^\star$ has mean $\mathbb{E}(y_i^\star) = 1/\alpha$. To measure how much a given realization deviates from this mean, we define

$$\zeta_i = y_i^\star - \frac{1}{\alpha}.$$

Thus, $\zeta_i$ has mean zero, and feasibility requires

$$y_i^\star > 0 \iff \zeta_i > -\frac{1}{\alpha}.$$

To make the distribution dimensionless and comparable to the standard normal, we divide by the standard deviation $\sigma_{y^\star}$:

$$\xi_i = \frac{\zeta_i}{\sigma_{y^\star}}.$$

Now each $\xi_i$ is a random variable following the standard normal distribution (mean zero and unit variance). The feasibility condition becomes

$$\xi_i > -\frac{1}{\alpha \sigma_{y^\star}}.$$

For a standard normal variable $\xi$, the probability that it is larger than a threshold $t$ is

$$\Pr(\xi > t) = 1 - \Phi(t),$$

where $\Phi$ is the cumulative distribution function (CDF) of the standard normal. If we plug $t = -1/(\alpha \sigma_{y^\star})$, we get

$$\Pr\Big(\xi_i > -\tfrac{1}{\alpha \sigma_{y^\star}}\Big) = 1 - \Phi\Big(-\tfrac{1}{\alpha \sigma_{y^\star}}\Big).$$

Because the standard normal distribution is symmetric about 0, we have

$$1 - \Phi(-z) = \Phi(z).$$

Therefore,

$$\Pr\Big(\xi_i > -\tfrac{1}{\alpha \sigma_{y^\star}}\Big) = \Phi\Big(\tfrac{1}{\alpha \sigma_{y^\star}}\Big). \tag{6}$$

Since we are treating the $\xi_i$ as independent, the probability that *all* $n$ inequalities hold is just the product of the probabilities for each one:

$$p_F(n, \alpha, \sigma^2_{y^\star}) \;\approx\; \Big[\Pr\Big(\xi_i > -\tfrac{1}{\alpha \sigma_{y^\star}}\Big)\Big]^n.$$

Substituting the expression 6, we obtain

$$p_F(n, \alpha, \sigma^2_{y^\star}) \;\approx\; \Phi\left(\frac{1}{\alpha \sigma_{y^\star}}\right)^n. \tag{7}$$

##### 3.8 Computing $\boldsymbol{\sigma_{y^\star}}$ using resolvents

As described in the section above, one can approximate the probability of feasibility using Eq. (7) (cf., Eq. 4 of the main text). To this end, one just needs to compute the variance of a single component of the equilibrium

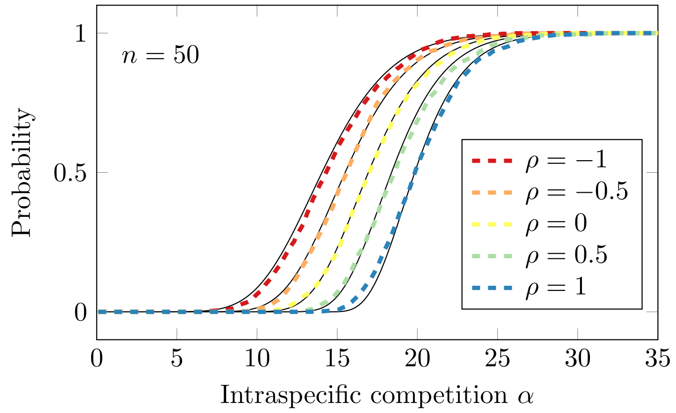


**Supplementary Figure 9**: Approximation using the resolvent method, cf. Eqs. (7) and (31), for matrices $B$ of size $n = 50$, whose entries are sampled from normal distribution with off-diagonal correlation $\rho$ (color). The approximations (solid lines) match the simulations (dashed lines) quite perfectly.

solution $y^\star$. Here, we provide an approximation for the standard deviation $\sigma_{y^\star}$, obtained by computing the expectation of the resolvent of the linear system

$$(\alpha I + B)x^\star = \mathbb{1}\,.$$

As a result of this calculation, we obtain an approximated formula for the deviation $\sigma_{y^\star}$ which, together with Eq. (7), is able to capture accurately the behavior of the probability of feasibility as a function of $\alpha$, even for small sizes. Importantly, this approximation is valid for arbitrary values of the correlation, $\rho \in [-1, 1]$. For example, Fig. 9 shows the quality of the approximation for $n = 50$, and different correlation values.

Because $B$ is a random matrix, the solution of the linear system (if it exists) is given by the random vector

$$x^\star = (\alpha I + B)^{-1}\mathbb{1}\,.$$

To calculate $\sigma_{y^\star}$: we first compute, in the limit of large system sizes ($n \to \infty$), the expectations of the equilibrium solution, $\mathbb{E}(x^\star)$, and the expectation of the scalar product $x^{\star T} x^\star$, $\mathbb{E}(x^{\star T} x^\star)$. With these two results, it is straightforward to evaluate asymptotically the variance $\sigma^2_{x^\star}$ of a single population as $\sigma^2_{x^\star} = \frac{1}{n}(\mathbb{E}(x^{\star T} x^\star) - \mathbb{E}(x^\star)^T \mathbb{E}(x^\star))$. Then, we use the proportionality relation between $x^\star$ and $y^\star$ to approximate the deviation $\sigma_{y^\star}$: because $y^\star = (\alpha I + C)^{-1}\mathbb{1}$ and $C = B - \mathbb{1} m^T$, one can use the Sherman-Morrison's formula to obtain

$$y^\star = \frac{1}{1 - m^T x^\star} x^\star.$$

Ignoring the correlation between $x^\star$ and $1 - m^T x^\star$, and approximating the expectation of the ratio by the ratio of expectations, we can write

$$y^\star = \frac{1}{1 - m^T x^\star} x^\star \approx \mathbb{E}\left(\frac{1}{1 - m^T x^\star}\right) x^\star \approx \frac{1}{\mathbb{E}\,(1 - m^T x^\star)} x^\star,$$

and therefore

$$\sigma_{y^\star} \approx \frac{\sigma_{x^\star}}{\mathbb{E}(1 - m^T x^\star)}, \tag{8}$$

where the standard deviations in this equation refer to the deviation of a single component of vectors $x^\star$ and $y^\star$ —all the components have the same expected deviation.

Because $\sigma^2_{x^\star} = \frac{1}{n}(\mathbb{E}({x^\star}^T x^\star) - \mathbb{E}(x^\star)^T \mathbb{E}(x^\star))$, approximating $\sigma_{y^\star}$ simply requires evaluating the expectations $\mathbb{E}(x^\star)$, $\mathbb{E}({x^\star}^T x^\star)$, and $\mathbb{E}(1 - m^T x^\star)$. We start with the cases $\rho = \pm 1$, which are easier to derive, and allow us to devise a route to obtain the general result for arbitrary $\rho$. In all the cases, it will be critical to employ a sort of "symmetrization" of matrix $B$ (which is only ensured for $\rho = 1$, in which case the matrix is indeed symmetric).

***Symmetric case,*** $\rho = 1$.

First, we consider the case $\rho = 1$, i.e., the fully symmetric case ($B_{ij} = B_{ji}$). We have to compute the aforementioned expectations.

*Computation of* $\mathbb{E}(x^\star)$ *using resolvents.* To begin with, we will focus on evaluating the expectation $\mathbb{E}(x^\star)$:

$$\mathbb{E}(x^\star) = \mathbb{E}((\alpha I + B)^{-1})\mathbb{1}.$$

Let $z \in \mathbb{C}$ be a complex number with non-zero imaginary part. Then the resolvent of matrix $B$ is defined as the inverse $(zI + B)^{-1}$, and we want to evaluate its expectation

$$G(z) = \mathbb{E}((zI + B)^{-1}),$$

computed over the ensemble of random matrices to which $B$ belongs. $G(z)$ can be expressed formally via the corresponding Neumann series expansion (which is always convergent if $\mathcal{Im}(z) > 0$). Our aim in this section is to obtain an asymptotic expression for $G(z)$, valid when $n \to \infty$, and then compute the expectations $\mathbb{E}(x^\star)$, $\mathbb{E}({x^\star}^T x^\star)$ and $\mathbb{E}(1 - m^T x^\star)$ in terms of $G(z)$. Once we have all the expressions, we will write $z = \alpha + i\epsilon$, and take the limit $\epsilon \to 0$ to obtain the final result.

In the limit $n \to \infty$, the expectation $G(z)$ of the resolvent satisfies the so-called matrix Dyson equation [25]:

$$G(z) = \frac{1}{z} I + \frac{1}{z}\mathbb{E}(BG(z)B)G(z). \tag{9}$$

This equation is exact for normally-distributed matrix entries $B_{ij}$, and is asymptotically true for arbitrary distributions (see Appendix G for a derivation of this equation).

In order to solve this nonlinear equation, we make a guess for the expectation $\Sigma := \mathbb{E}(BG(z)B)$. Let $G_{ij}$ be the elements of matrix $G(z)$. Since $\mathbb{E}(B_{ij}B_{kl}) = \delta_{ik}\delta_{jl} + \delta_{il}\delta_{jk}$ (where $\delta_{ij}$ stands for the Kronecker symbol) holds for $\rho = 1$, then

$$\Sigma_{il} = \mathbb{E}\left(\sum_{j,k} B_{ij} G_{jk} B_{kl}\right) = \sum_{j,k} \mathbb{E}(B_{ij}B_{kl}) G_{jk},$$

because $G_{jk}$ is an expectation itself. Hence

$$\Sigma_{il} = G_{li} + \delta_{il} \sum_j G_{jj}.$$

Therefore, in the limit $n \to \infty$, it is expected that the diagonal of matrix $\Sigma$ dominates the off-diagonal entries. Thus, we adopt the guess $\Sigma = a(z)I$, and look for solutions for the function $a(z)$ —observe that, given that populations are statistically equivalent, we expect all the diagonal elements of the expectation $\mathbb{E}(BG(z)B)$ to be the same.

We substitute this assumption into Eq. (9) and solve for $G(z)$:

$$G(z) = \frac{1}{z}I + \frac{a(z)}{z}G(z); \qquad G(z) = (z - a(z))^{-1}I.$$

Now we can compute $\Sigma$ in a different way:

$$\Sigma = \mathbb{E}(BG(z)B) = (z - a(z))^{-1}\mathbb{E}(B^2) = \frac{n-1}{z - a(z)}I = a(z)I,$$

where we have used that $\mathbb{E}((B^2)_{ii}) = \sum_j \mathbb{E}(B_{ij}B_{ji}) = \sum_j \mathbb{E}(B_{ij}^2) = n - 1$, and the last step follows because the sum over $j$ only contains $n-1$ terms (given that $B_{ii} = 0$). In addition, if $i \neq k$, we have that $\mathbb{E}((B^2)_{ik}) = \sum_j \mathbb{E}(B_{ij}B_{jk}) = \sum_j(\delta_{ij}\delta_{jk} + \delta_{ik}\delta_{jj}) = \delta_{ik} + \delta_{ik}\sum_j \delta_{jj} = 0$. Thus, $\mathbb{E}(B^2) = (n-1)I$ and the last equation implies that the unknown function $a(z)$ satisfies the quadratic equation

$$a(z)(z - a(z)) = n - 1.$$

Solving for $a(z)$, we obtain

$$a(z) = \frac{z \pm \sqrt{z^2 - 4(n-1)}}{2}.$$

According to the definition of the resolvent, $(zI + B)^{-1}$, it must behave as $z^{-1}I$ as $z \to \infty$. Therefore, Dyson equation (9) implies that $z^{-1}\Sigma \to 0$ as $z \to \infty$, which means that we have to choose the minus sign in $a(z)$. Consequently, the resolvent expectation is expressed as

$$G(z) = (z - a(z))^{-1}I = \frac{2}{z + \sqrt{z^2 - 4(n-1)}}I = \frac{z - \sqrt{z^2 - 4(n-1)}}{2(n-1)}I.$$

Having computed the resolvent expectation in the asymptotic limit $n \to \infty$, we immediately find an expression for the expected value of $x^\star$:

$$\mathbb{E}(x^\star) = \lim_{\epsilon \to 0} \mathbb{E}(((\alpha + i\epsilon)I + B)^{-1})\mathbb{1} = \lim_{\epsilon \to 0} G(\alpha + i\epsilon)\mathbb{1} = \frac{\alpha - \sqrt{\alpha^2 - 4(n-1)}}{2(n-1)}\mathbb{1}. \tag{10}$$

*Computation of* $\mathbb{E}(x^{\star T}x^\star)$. The second expectation to be computed can be written as

$$\mathbb{E}(x^{\star T}x^\star) = \mathbb{1}^T\mathbb{E}(((\alpha I + B)^{-1})^T(\alpha I + B)^{-1})\mathbb{1} = \mathbb{1}^T\mathbb{E}((\alpha I + B)^{-2})\mathbb{1},$$

because $((\alpha I + B)^{-1})^T = ((\alpha I + B)^T)^{-1} = (\alpha I + B)^{-1}$ (since $B^T = B$). Here we denote by $(\alpha I + B)^{-2}$ the square power of $(\alpha I + B)^{-1}$. Thus, we need to evaluate the expectation of $(\alpha I + B)^{-2}$. Let us define it as

$$K(z) = \mathbb{E}((\alpha I + B)^{-2}).$$

We show in the Appendix that $K(z)$ can be written in a simple way in terms of the derivative of $G(z)$, namely:

$$K(z) = -G'(z) = \frac{1}{2(n-1)}\left(\frac{z}{\sqrt{z^2 - 4(n-1)}} - 1\right) I, \tag{11}$$

which implies that

$$\mathbb{E}(x^{\star T} x^\star) = \lim_{\epsilon \to 0} \mathbb{1}^T K(\alpha + i\epsilon)\mathbb{1} = \frac{n}{2(n-1)}\left(\frac{\alpha}{\sqrt{\alpha^2 - 4(n-1)}} - 1\right). \tag{12}$$

Using the expression obtained above for $\mathbb{E}(x^\star)$, we find, after simplifications, the following asymptotic limit for the variance $\sigma^2_{x^\star}$:

$$\sigma^2_{x^\star} = \frac{1}{n}(\mathbb{E}(x^{\star T} x^\star) - \mathbb{E}(x^\star)^2) = \frac{1}{2(n-1)^2}\left(n - 1 - \alpha^2 + \frac{\alpha(\alpha^2 - 3(n-1))}{\sqrt{\alpha^2 - 4(n-1)}}\right). \tag{13}$$

*Computation of* $\mathbb{E}(1 - m^T x^\star)$. To evaluate the last expectation we need, we first observe that $(\alpha I + B)x^\star = \mathbb{1}$ implies, by left-multiplying by $\mathbb{1}^T$, that

$$\alpha \mathbb{1}^T x^\star + \mathbb{1}^T B x^\star = n.$$

Equivalently,

$$\frac{\alpha}{n}\mathbb{1}^T x^\star + m^T x^\star = 1,$$

because, by definition, $m = \frac{1}{n}B^T\mathbb{1}$. Therefore

$$1 - m^T x^\star = \frac{\alpha}{n}\mathbb{1}^T x^\star.$$

The expectation $\mathbb{E}(\mathbb{1}^T x^\star) = \mathbb{1}^T \mathbb{E}(x^\star)$ follows from $\mathbb{E}(x^\star)$: using Eq. (10), we finally obtain

$$\mathbb{E}(1 - m^T x^\star) = \frac{\alpha}{n}\mathbb{1}^T \mathbb{E}(x^\star) = \frac{\alpha^2 - \alpha\sqrt{\alpha^2 - 4(n-1)}}{2(n-1)}.$$

Substituting this result and Eq. (13) into Eq. (8), and after some simplifications, the approximation of $\sigma_{y^\star}$ finally reads

$$\sigma_{y^\star} \approx \frac{1}{\alpha}\sqrt{\frac{\alpha}{2\sqrt{\alpha^2 - 4(n-1)}} - \frac{1}{2}}. \tag{14}$$

This result is valid for $\rho = 1$, large $n$, and is well defined for $\alpha > 2\sqrt{n-1}$. In particular, this condition is satisfied for $\alpha \geq 2\sqrt{n} = \alpha_S$, i.e., above the global stability threshold for $\rho = 1$. The choice of the distribution for $B_{ij}$ has no effect (asymptotically). This approximation compares very well with simulated data for the distributions considered here (Fig. 9), even for small $n$.

***Skew-symmetric case,*** $\rho = -1$**.**

In order to proceed as in the symmetric case, we have to find a Dyson equation satisfied by the resolvent expectation $\mathbb{E}((\alpha I + B)^{-1})$. The formalism of the matrix Dyson equation can only be applied to Hermitian matrices [25]; however, for $\rho = -1$ matrix $B$ is skew-symmetric ($B^T = -B$). In order for the Dyson equation to hold, we transform the original matrix in order to make it Hermitian: let $Q = iB$, where $i$ is the imaginary unit. Then $Q^\dagger = (iB)^\dagger = -iB^T = iB = Q$, and hence $Q$ is Hermitian (here, $^\dagger$ means complex conjugate transpose).

To solve $(\alpha I + B)x^\star = 1$ we focus on the transformed linear system

$$(i\zeta I + iB)x^\star = i\mathbb{1},$$

for $\zeta \in \mathbb{C}$. Then let $z = i\zeta$ (with positive imaginary part) and write the system to solve as $(zI + Q)x^\star = i\mathbb{1}$, with $Q$ Hermitian. We just need to compute the expectation

$$G(z) = \mathbb{E}((zI + Q)^{-1})$$

The same matrix Dyson equation holds, see Eq. (9):

$$G(z) = \frac{1}{z}I + \frac{1}{z}\mathbb{E}(QG(z)Q)G(z). \tag{15}$$

Following the same arguments as before, we look for solutions such that $\mathbb{E}(BG(z)B) = a(z)I$. Therefore,

$$\mathbb{E}(QG(z)Q) = i^2\mathbb{E}(BG(z)B) = -a(z)I.$$

We introduce this expression into the Dyson equation and solve for $G(z)$:

$$G(z) = (z + a(z))^{-1}I.$$

Now, given that $G(z)$ is a multiple of the identity, we can evaluate $\mathbb{E}(QG(z)Q)$ in a different way:

$$\mathbb{E}(QG(z)Q) = -\mathbb{E}(BG(z)B) = -(z + a(z))^{-1}\mathbb{E}(B^2).$$

As for the symmetric case, one can check that $\mathbb{E}(B_{ij}^2) = 0$ for $i \neq j$. Only diagonal expectations remain: $\mathbb{E}(B_{ii}^2) = \sum_j \mathbb{E}(B_{ij}B_{ji}) = -\sum_j \mathbb{E}(B_{ij}^2) = -(n-1)$, because $B_{ii} = 0$ by definition. Therefore,

$$\mathbb{E}(QG(z)Q) = \frac{n-1}{z + a(z)}I.$$

As a consequence, $a(z)$ must satisfy the quadratic equation

$$a(z)(z + a(z)) + n - 1 = 0,$$

which yields

$$a(z) = \frac{-z \pm \sqrt{z^2 - 4(n-1)}}{2},$$

and

$$G(z) = \frac{2}{z \pm \sqrt{z^2 - 4(n-1)}} I.$$

In order for $G(z) \sim \frac{1}{z} I$ as $z \to \infty$, we have to choose the positive sign. Then, we can write

$$G(z) = \frac{z - \sqrt{z^2 - 4(n-1)}}{2(n-1)} I.$$

Now we go back to the original variable $z = i\zeta$ and matrix $Q = iB$:

$$G(i\zeta) = \mathbb{E}((i\zeta I + iB)^{-1}) = i^{-1}\mathbb{E}((\zeta I + B)^{-1}) = -i\mathbb{E}((\zeta I + B)^{-1}) = -iG(\zeta),$$

where $G(\zeta) = \mathbb{E}((\zeta I + B)^{-1})$ is the expectation we want to evaluate. On the other hand,

$$G(i\zeta) = -i\frac{\sqrt{\zeta^2 + 4(n-1)} - \zeta}{2(n-1)} I = -iG(\zeta).$$

Thus, by substituting $\zeta = \alpha + i\epsilon$, and taking the limit $\epsilon \to 0$, we find the following expression for the resolvent expectation in the skew-symmetric case:

$$G(\alpha) = \lim_{\epsilon \to 0} \mathbb{E}(((\alpha + i\epsilon)I + B)^{-1}) = \frac{\sqrt{\alpha^2 + 4(n-1)} - \alpha}{2(n-1)} I. \tag{16}$$

This result automatically implies that $\mathbb{E}(x^\star) = g(\alpha)\mathbb{1}$, where we have defined

$$g(\alpha) = \frac{\sqrt{\alpha^2 + 4(n-1)} - \alpha}{2(n-1)}. \tag{17}$$

*Computation of* $\mathbb{E}(x^{\star T} x^\star)$. Because $x^\star = (\alpha I + B)^{-1}\mathbb{1}$ and $x^{\star T} = \mathbb{1}^T(\alpha I - B)^{-1}$ (we have used, in the last calculation, that $B^T = -B$), we can write the expectation $\mathbb{E}(x^{\star T} x^\star)$ as

$$\mathbb{E}(x^{\star T} x^\star) = \mathbb{1}^T \mathbb{E}((\alpha I - B)^{-1}(\alpha I + B)^{-1})\mathbb{1}.$$

Now we resort to the identity

$$(\alpha I - B)^{-1}(\alpha I + B)^{-1} = \frac{1}{2\alpha}(\alpha I + B)^{-1} + \frac{1}{2\alpha}(\alpha I - B)^{-1},$$

which can be easily proven by left- and right-multiplying both sides by $\alpha I - B$ and $\alpha I + B$, respectively. By taking expectations, we obtain

$$\mathbb{E}((\alpha I - B)^{-1}(\alpha I + B)^{-1}) = \frac{1}{2\alpha}\mathbb{E}((\alpha I + B)^{-1}) + \frac{1}{2\alpha}\mathbb{E}((\alpha I - B)^{-1}).$$

It holds that $\mathbb{E}((\alpha I - B)^{-1}) = \mathbb{E}((\alpha I + B)^{-1})$ —this can be verified by solving Eq. (15) for $Q = iB^T = -iB$. Therefore,

$$\mathbb{E}((\alpha I - B)^{-1}(\alpha I + B)^{-1}) = \frac{1}{\alpha}\mathbb{E}((\alpha I + B)^{-1}) = \frac{g(\alpha)}{\alpha} I, \tag{18}$$

and $\mathbb{E}(x^{\star T} x^\star) = ng(\alpha)/\alpha$. Notice that $\mathbb{E}(x^\star)^T \mathbb{E}(x^\star) = ng(\alpha)^2$, and thus

$$\sigma_{x^\star}^2 = \frac{1}{n}(\mathbb{E}(x^{\star T} x^\star) - \mathbb{E}(x^\star)^T \mathbb{E}(x^\star)) = \frac{g(\alpha)}{\alpha} - g(\alpha)^2. \tag{19}$$

*Final approximation for $\sigma_{y^\star}$ in the skew-symmetric case.* The calculation we did for the expectation $\mathbb{E}(1 - m^T x^\star)$ in the symmetric case is still valid in the skew-symmetric case,

$$\mathbb{E}(1 - m^T x^\star) = \frac{\alpha}{n}\mathbb{1}^T \mathbb{E}(x^\star) = \alpha g(\alpha).$$

Finally, substituting the last expression and Eq. (19) into Eq. (8), we find

$$\sigma_{y^\star} \approx \frac{\sqrt{g(\alpha)(\alpha^{-1} - g(\alpha))}}{\alpha g(\alpha)} = \frac{1}{\alpha}\sqrt{\frac{1}{\alpha g(\alpha)} - 1} = \frac{1}{\alpha}\sqrt{\frac{\sqrt{\alpha^2 + 4(n-1)}}{2\alpha} - \frac{1}{2}}, \tag{20}$$

where we have used that $1/g(\alpha) = (\sqrt{\alpha^2 + 4(n-1)} + \alpha)/2$ and performed some straightforward manipulations. Again, this expression is universal and does not depend (in the limit of large $n$) on the distribution chosen to draw matrix elements. It compares very well with simulations (Fig. 9).

***General case: arbitrary correlation.***

We take further advantage of the theory of resolvents to compute an approximation for $\sigma_{y^\star}$ for arbitrary values of the correlation coefficient $\rho$. We recover the two limits studied above ($\rho = 1$ and $\rho = -1$) as particular cases of this more general calculation. The rationale of the proof in the general case is based again on a matrix Dyson equation. Having studied first the limiting cases $\rho = 1$ and $\rho = -1$, for which the derivation is simpler, one can intuit the general route to be followed to tackle the general case with arbitrary correlation.

One critical feature of the Dyson equation used in both cases is that it applies for Hermitian matrices. For $\rho = 1$, matrix $B$ is symmetric by definition, and for $\rho = -1$ we applied the matrix Dyson equation used for $\rho = 1$ to matrix $Q = iB$, which is Hermitian (since satisfies $Q^\dagger = Q$). For arbitrary $\rho$ we resort to the "Hermitization trick" due to Girko [26]: consider, for $z \in \mathbb{C}$, the $(2n) \times (2n)$ matrix

$$Q(z) = \begin{pmatrix} \mathbb{0} & zI + B \\ z^\star I + B^T & \mathbb{0} \end{pmatrix},$$

where $z^\star$ denotes the conjugate of complex number $z$, and $\mathbb{0}$ stands for the null matrix of dimension $n$. Matrix $Q(z)$ is called the Hermitization of matrix $zI + B$, for $B$ real. One can see that $Q$ is Hermitian, i.e., $Q^\dagger = Q$.

$Q(z)$ belongs to a general class of self-adjoint (Hermitian) random matrices with correlated entries, for which Alt and Krüger proved in [27] that the resolvent of matrix $Q$, defined as

$$S(z,w) := (iwI_2 + Q(z))^{-1},$$

satisfies, in the limit of large $n$, the following non-linear matrix Dyson equation:

$$-\begin{pmatrix} I & \mathbb{0} \\ \mathbb{0} & I \end{pmatrix} = \left( \begin{pmatrix} iwI & -zI \\ -z^\star I & iwI \end{pmatrix} + \Lambda(S(z,w)) \right) S(z,w), \tag{21}$$

where $I$ is the identity matrix of dimension $n$, $I_2$ is the $(2n) \times (2n)$ identity matrix, and $\Lambda$ is a matrix operator defined, for an arbitrary complex, square matrix $A$ with dimension $(2n) \times (2n)$, as

$$\Lambda(A) := \mathbb{E}((Q(z) - Z)A(Q(z) - Z)),$$

where $Z := \begin{pmatrix} \mathbb{0} & zI \\ z^\star I & \mathbb{0} \end{pmatrix}$. Observe that the expectation $\mathbb{E}((iwI_2+Q(z))^{-1})$ also satisfies the same matrix Dyson equation, which is somewhat an extension of the Dyson equation used for $\rho = 1$ to "hermitized" matrices.

Can we extract the resolvent $(zI + B)^{-1}$ from the resolvent $S(z,w)$ of the Hermitized matrix? Let us partition $S$ in two $n \times n$ square blocks as

$$S(z,w) = \begin{pmatrix} S_{11} & S_{12} \\ S_{21} & S_{22} \end{pmatrix}.$$

Then, matrix $S$ satisfies

$$\begin{pmatrix} iwI & zI + B \\ z^\star I + B^T & iwI \end{pmatrix} \begin{pmatrix} S_{11} & S_{12} \\ S_{21} & S_{22} \end{pmatrix} = \begin{pmatrix} I & \mathbb{0} \\ \mathbb{0} & I \end{pmatrix},$$

which implies

$$iwS_{11} + (zI + B)S_{21} = I.$$

Therefore, the resolvent $(zI + B)^{-1}$ we want to calculate is the lower-left block of the augmented resolvent $S(z,w)$ evaluated at $w = 0$. Our goal is to compute an asymptotic approximation to $S(z,w)$ via the solution of the matrix Dyson equation (21).

As shown in [27], in the limit $n \to \infty$, the solution to Eq. (21) is of the form

$$S(z,w) = \begin{pmatrix} iv(z,w)I & b(z,w)^\star I \\ b(z,w)I & iv(z,w)I \end{pmatrix}, \tag{22}$$

where the imaginary part of $v$ is positive, $b$ takes values on the complex plane, and $S$ satisfies $\mathcal{I}m(S) = \frac{1}{2i}(S - S^\dagger)$, which is positive definite. Since we are interested in the particular value $w = 0$, we can compute explicit expressions for functions $v$ and $b$ in terms of $z$. From now on, we omit the dependence on $z$ and $w$ in $v$ and $b$.

In order to obtain explicit forms for $v$ and $b$ in terms of $z$, we first evaluate the expectation $\Lambda(S(z,w)) = \mathbb{E}((Q(z)-Z)S(z,w)(Q(z)-Z))$ at the solution given by Eq. (22). This yields

$$\Lambda(S(z,w)) = \begin{pmatrix} iv\mathbb{E}(BB^T) & b\mathbb{E}(B^2) \\ b^\star\mathbb{E}((B^T)^2) & iv\mathbb{E}(B^TB) \end{pmatrix} \tag{23}$$

Recall that $\mathbb{E}(B_{ij}^2) = 1$ and $\mathbb{E}(B_{ij}B_{ji}) = \rho$ for $j \neq i$, which implies that $\mathbb{E}(B^TB) = \mathbb{E}(BB^T) = (n-1)I$ and $\mathbb{E}(B^2) = \mathbb{E}((B^T)^2) = \rho(n-1)I$. Then $\Lambda(S(z,w))$ simplifies to

$$\Lambda(S(z,w)) = (n-1)\begin{pmatrix} ivI & b\rho I \\ b^\star\rho I & ivI \end{pmatrix}.$$

We obtain the non-linear system of equations satisfied by $v$ and $b$ by substituting Eqs. (22) and (23) into the matrix Dyson equation (21):

$$\begin{aligned} -1 &= -wv - zb + (n-1)(-v^2 + \rho b^2), \\ 0 &= wb^\star - zv + (n-1)(vb^\star + \rho vb). \end{aligned} \tag{24}$$

The two remaining equations are simply the complex conjugate of the two equations above. From the second equation we can solve for $v$:

$$v = -\frac{wb^\star}{(n-1)b^\star + (n-1)\rho b - z}, \tag{25}$$

and substitute into the first equation to get a fourth-order equation to be satisfied by $b$:

$$((n-1)\rho b^2 - zb + 1)((n-1)b^\star + (n-1)\rho b - z)^2 + w^2 b^\star((n-1)b^\star + (n-1)\rho b - z) - (n-1)w^2 b^{\star 2} = 0.$$

Since the solution of this polynomial equation will be a continuous function of $w$ at $w = 0$, we can set $w = 0$ to get

$$((n-1)\rho b^2 - zb + 1)((n-1)b^\star + (n-1)\rho b - z)^2 = 0.$$

For $v$ to be well defined, the only solution for $b$ is the one that cancels the first factor:

$$b(z,0) = \frac{z - \sqrt{z^2 - 4\rho(n-1)}}{2\rho(n-1)}, \tag{26}$$

where we have chosen the minus sign so that the resolvent $(zI + B)^{-1}$ behaves as $z^{-1}I$ in the limit $z \to \infty$ (recall that $(zI+B)^{-1} \approx S_{21}|_{w=0} = b(z,0)I$ in the limit of large $n$). Therefore, we can obtain the expectation $\mathbb{E}(x^\star)$ by substituting $z = \alpha + i\epsilon$ in the expression for $b(z,0)$ and letting $\epsilon \to 0$:

$$\mathbb{E}(x^\star) = \lim_{\epsilon\to 0} b(\alpha + i\epsilon, 0)\mathbb{1} = \frac{\alpha - \sqrt{\alpha^2 - 4\rho(n-1)}}{2\rho(n-1)}\mathbb{1}. \tag{27}$$

Note that this expectation is always a positive real number for $\alpha > 2\sqrt{\rho(n-1)}$, even for negative $\rho$. By taking the limit $\rho \to 0$ in the last equation we obtain $\mathbb{E}(x^\star) = \alpha^{-1}\mathbb{1}$ for $\rho = 0$. One can easily check that Eq. (27) reduces to Eq. (10) for $\rho = 1$ and Eq. (17) for $\rho = -1$.

*Computation of* $\mathbb{E}(x^{\star T} x^\star)$. As for the derivations above, we need to calculate the expectation $\mathbb{E}(x^{\star T} x^\star)$ in order to approximate $\sigma_y$ according to Eq. (8). Here we follow an approach similar to the $\rho = -1$ case. Let us focus on the matrix product

$$W := (iwI_2 + Q(z))^{-1\dagger}(iwI_2 + Q(z))^{-1} = (-iw^\star I_2 + Q(z))^{-1}(iwI_2 + Q(z))^{-1},$$

where in the last step we have used $Q^\dagger = Q$. This is in analogy with the product $(zI - B)^{-1}(zI + B)^{-1}$, which was used to compute the expectation $\mathbb{E}(x^{\star T} x^\star)$ in the skew-symmetric case ($\rho = -1$). For arbitrary correlation we use the resolvent of the Hermitization instead. By writing $W$ as a block matrix, it satisfies

$$(iwI_2 + Q(z))^\dagger W(iwI_2 + Q(z)) = \begin{pmatrix} -iw^\star I & zI + B \\ z^\star I + B^T & -iw^\star I \end{pmatrix} \begin{pmatrix} W_{11} & W_{12} \\ W_{21} & W_{22} \end{pmatrix} \begin{pmatrix} iwI & zI + B \\ z^\star I + B^T & iwI \end{pmatrix} = \begin{pmatrix} I & \mathbb{0} \\ \mathbb{0} & I \end{pmatrix}.$$

Setting $w = 0$, the lower-right block of this equality implies

$$(z^\star I + B^T)W_{11}(zI + B) = I, \qquad W_{11} = (z^\star I + B^T)^{-1}(zI + B)^{-1}.$$

Therefore, the expectation of the upper-left block of matrix $W$ is precisely related to the expectation we want to compute:

$$\mathbb{E}(x^{\star T} x^\star) = \mathbb{1}^T \mathbb{E}((\alpha I + B^T)^{-1}(\alpha I + B)^{-1})\mathbb{1}.$$

Thus, our next goal is to compute $W$. To this end, as we did in the skew-symmetric case, we write

$$W = c_1(-iw^\star I_2 + Q(z))^{-1} + c_2(iwI_2 + Q(z))^{-1}.$$

Substituting into $(iwI_2 + Q(z))^\dagger W(iwI_2 + Q(z)) = I_2$ we obtain the condition

$$c_1(iwI_2 + Q(z)) + c_2(-iw^\star I_2 + Q(z)) = I_2,$$

which implies that $c_1 = -c_2$ and $i(c_1 w - c_2 w^\star) = 1$. Therefore, $c_1 = \frac{i}{w + w^\star} = -c_2$ and we can write

$$W = -\frac{i}{w + w^\star}\left((iwI_2 + Q(z))^{-1} - (-iw^\star I_2 + Q(z))^{-1}\right) = -\frac{i}{w + w^\star}(S(z, w) - S(z, -w^\star)).$$

We now use the asymptotic limit of matrix $S$, given by Eq. (22), and Eq. (25) to obtain

$$W_{11} = \frac{v(z,w) - v(z,-w^\star)}{w + w^\star} I = -\frac{b^\star}{(n-1)b^\star + (n-1)\rho b - z} I,$$

which turns out to be independent of $w$ (this is true in the limit $n \to \infty$). Therefore,

$$\mathbb{E}((\alpha I + B^T)^{-1}(\alpha I + B)^{-1}) = \lim_{\epsilon \to 0} \frac{-b^\star(\alpha + i\epsilon, 0)}{(n-1)b^\star(\alpha + i\epsilon, 0) + (n-1)\rho b(\alpha + i\epsilon, 0) - (\alpha + i\epsilon)} I. \tag{28}$$

Using Eq. (26), and after some simplifications, we finally get the asymptotic formula

$$\mathbb{E}((\alpha I + B^T)^{-1}(\alpha I + B)^{-1}) = \frac{2}{\alpha(\alpha + \sqrt{\alpha^2 - 4\rho(n-1)}) - 2(1+\rho)(n-1)} I. \tag{29}$$

One can use this expression to recover Eq. (12) for $\rho = 1$, and Eq. (18) for $\rho = -1$.

*Final result for the deviation $\sigma_{y^\star}$.* Let us define

$$g(\alpha) := \frac{\alpha - \sqrt{\alpha^2 - 4\rho(n-1)}}{2\rho(n-1)}, \qquad g_2(\alpha) := \frac{2}{\alpha^2 + \alpha\sqrt{\alpha^2 - 4\rho(n-1)}) - 2(1+\rho)(n-1)}.$$

Hence, $\mathbb{E}(x^\star) = g(\alpha)\mathbb{1}$ and $\mathbb{E}({x^\star}^T x^\star) = n g_2(\alpha)$. This allows to approximate the variance of $x^\star$ as

$$\sigma^2_{x^\star} = \frac{1}{n}(\mathbb{E}({x^\star}^T x^\star) - \mathbb{E}(x^\star)^T \mathbb{E}(x^\star)) = g_2(\alpha) - g(\alpha)^2.$$

As in the particular cases $\rho = \pm 1$, the expectation $\mathbb{E}(1 - m^T x^\star)$ can be computed as

$$\mathbb{E}(1 - m^T x^\star) = \frac{\alpha}{n} \mathbb{1}^T \mathbb{E}(x^\star) = \alpha g(\alpha).$$

Finally, substituting into Eq. (8), we obtain:

$$\sigma_{y^\star} \approx \frac{\sqrt{g_2(\alpha) - g(\alpha)^2}}{\alpha g(\alpha)} = \frac{1}{\alpha}\sqrt{\frac{g_2(\alpha)}{g(\alpha)^2} - 1}. \tag{30}$$

After some simplifications, we deduce:

$$\frac{1}{\alpha \sigma_{y^\star}} \approx \sqrt{\frac{\alpha^2 + \alpha\sqrt{\alpha^2 - 4\rho(n-1)}}{2(n-1)} - (1+\rho)}, \tag{31}$$

which is Eq. 4 of the main text. For $\rho \leq 0$ this expression always yields a real number, and for $\rho > 0$ does so when $\alpha \geq 2\sqrt{\rho(n-1)}$; in both cases, the stability threshold $\alpha_S = \sqrt{2n(1+\rho)}$ is contained within the interval in which Eq. (31) is valid. The formula reduces to Eqs. (14) and (20) for $\rho = 1$ and $\rho = -1$, respectively. The agreement with the probability of feasibility obtained by sampling matrices is remarkable (Fig. 9), even for moderate matrix sizes.

Observe that $(\alpha \sigma_{y^\star})^{-1}$, i.e., the argument of the standardized normal cumulative density function in Eq. (7), can be written in terms of the re-scaled intraspecific competition $\tilde{\alpha} := \alpha/\sqrt{n-1}$, thereby being independent of $n$:

$$\frac{1}{\alpha \sigma_{y^\star}} \approx \sqrt{\frac{\tilde{\alpha}^2 + \tilde{\alpha}\sqrt{\tilde{\alpha}^2 - 4\rho}}{2} - (1+\rho)}.$$

However, this does not make all the probability of feasibility curves to collapse for different system sizes, because the probability of feasibility is precisely $\Phi((\alpha\sigma_{y^\star})^{-1})^n$, and rising the cumulative density function to the power of $n$ introduces an intrinsic separation between the curves for finite $n$ (see Fig. 10, which provides a comparison between the resolvent approximation and the approximation via Richardson iteration presented below). This separation converges to zero as $n$ increases. We show here that, as $n \to \infty$, the median of the PDF for feasibility as function of $\alpha$, given by $\frac{d}{d\alpha}p_F(\alpha)$, grows unboundedly, whereas the width of the distribution tends to zero as $n$ increases. The convergence to zero width is extremely slow, though. Mathematically, in the limit of infinitely many species, the transition from infeasibility to feasibility is sharp, although the transition point (which can be assimilated to the median of the PDF) occurs at $\tilde{\alpha}_{1/2} = \infty$. Given the slow rate of convergence to this limiting situation, even for very large values of $n$ we will observe a smooth curve for the probability of feasibility (see Fig. 10).

For finite $n$, we define the critical value of $\tilde{\alpha}$ as the intraspecific competition strength $\tilde{\alpha}_{1/2}$ such that $p_F(\tilde{\alpha}_{1/2}) = \frac{1}{2}$. Using the resolvent approximation, this condition is equivalent to

$$\sqrt{\frac{\tilde{\alpha}_{1/2}^2 + \tilde{\alpha}_{1/2}\sqrt{\tilde{\alpha}_{1/2}^2 - 4\rho}}{2}} - (1+\rho) = \sqrt{2}\mathrm{erf}^{-1}\left(2e^{-\frac{1}{n}\log 2} - 1\right),$$

where we used that the inverse CDF of the standard normal is expressed as $\Phi^{-1}(y) = \sqrt{2}\mathrm{erf}^{-1}(2y-1)$. Let $x_n := \sqrt{2}\mathrm{erf}^{-1}\left(2e^{-\frac{1}{n}\log 2} - 1\right)$. Then we can solve for $\tilde{\alpha}_{1/2}$ to obtain

$$\tilde{\alpha}_{1/2} = \frac{1+\rho+x_n^2}{\sqrt{1+x_n^2}}.$$

Because $\lim_{n\to\infty} x_n = \sqrt{2}\mathrm{erf}^{-1}(1) = \infty$, we obtain that $\lim_{n\to\infty}\tilde{\alpha}_{1/2} = \lim_{n\to\infty}|x_n| = +\infty$. Therefore, the "center" of the PDF, $\alpha_{1/2} = \sqrt{n-1}\tilde{\alpha}_{1/2}$, tends to infinity as well. This is consistent with previous estimations of this threshold[15, 16].

With regard to the width of the distribution, we use the quantiles $\tilde{\alpha}_\beta$ and $\tilde{\alpha}_{1-\beta}$ to define the width of the distribution: let $\tilde{\alpha}_\beta$ be such that $p_F(\tilde{\alpha}_\beta) = \beta$ (with $0 < \beta < 1$), i.e.,

$$\sqrt{\frac{\tilde{\alpha}_{\beta}^2 + \tilde{\alpha}_{\beta}\sqrt{\tilde{\alpha}_{\beta}^2 - 4\rho}}{2}} - (1+\rho) = \sqrt{2}\mathrm{erf}^{-1}\left(2e^{\frac{1}{n}\log\beta} - 1\right) =: x_\beta,$$

and the same definitions for $\tilde{\alpha}_{1-\beta}$ and $x_{1-\beta}$. Let $\beta$ be small (for example, $\beta = 0.01$). The width of the PFD can be estimated as the difference $\tilde{\alpha}_{1-\beta} - \tilde{\alpha}_\beta$, which asymptotically tends to zero. Indeed,

$$\lim_{n\to\infty}(\tilde{\alpha}_{1-\beta} - \tilde{\alpha}_\beta) = \lim_{n\to\infty}\left(\frac{1+\rho+x_{1-\beta}^2}{\sqrt{1+x_{1-\beta}^2}} - \frac{1+\rho+x_\beta^2}{\sqrt{1+x_\beta^2}}\right) = \lim_{n\to\infty}(|x_{1-\beta}| - |x_\beta|).$$

The last limit can be evaluated as follows:

$$\lim_{n\to\infty}(|x_{1-\beta}| - |x_\beta|) = \sqrt{2}\lim_{n\to\infty}\left(\left|\mathrm{erf}^{-1}\left(2e^{\frac{1}{n}\log(1-\beta)} - 1\right)\right| - \left|\mathrm{erf}^{-1}\left(2e^{\frac{1}{n}\log\beta} - 1\right)\right|\right) =$$
$$\sqrt{2}\lim_{n\to\infty}\left(\mathrm{erf}^{-1}(1) - \mathrm{erf}^{-1}(1)\right) = 0,$$

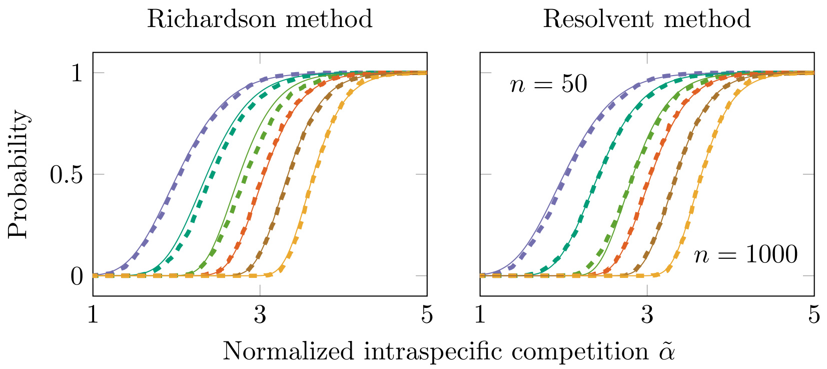


**Supplementary Figure 10**: Approximate probability of feasibility for the two approximations presented in the text. Left panel: approximation based on Richardson's iteration; Right panel: approximation based on resolvents. Intraspecific competition is normalized ($\tilde{\alpha} = \alpha/\sqrt{n-1}$) so that the argument of $\Phi(\cdot)$, i.e., the CDF of the standard normal distribution that appears in Eq. 4 of the main text, does not depend on $n$. We observe a very good agreement with simulations for both methods, even for small sizes: colder colors illustrate the case of $n = 50$ (with purple for $\rho = -1$, teal for $\rho = 0$ and green for $\rho = 1$) and warmer colors the case of $n = 1000$ (with orange, brown and yellow marking $\rho = -1, 0, 1$, respectively). Dashed lines report simulated data. While Richardson's method tends to overestimate $p_F$ for small sizes, the resolvent method overestimates (underestimates) it for negative (positive) correlations and small sizes. Finally, note the very slow convergence to a sharp transition with $n$, and the displacement to the transition to the right.

because both $e^{\frac{1}{n}\log(1-\beta)}$ and $e^{\frac{1}{n}\log\beta}$ tend to one as $n \to \infty$. Therefore, in the limit $n \to \infty$, the width of the distribution tends to zero, thus the transition becomes sharper and sharper, and displaces to the right without reaching a finite transition value in the limit, because the transition occurs at $\tilde{\alpha}_{1/2} \to \infty$ (the median grows unboundedly). It is important to remark that the convergence of $p_F(\tilde{\alpha})$ to a step function centered at $\tilde{\alpha} = \infty$ is very slow, as Fig. 10 shows.

Finally, we use this approximation to estimate the probability of feasibility at the stability threshold, $\alpha_S \approx \sqrt{2n(1+\rho)}$, obtaining:

$$p_F(\alpha_S) \approx \Phi\left(\sqrt{\frac{1+\rho+\sqrt{n(1+\rho)(n-(n-2)\rho)}}{n-1}}\right)^n .$$

The probability of feasibility at the stability threshold tends to zero as $n$ increases. For example, for $n = 50$, we find $p_F(\alpha_S) \approx 2.4 \times 10^{-4}$ for $\rho = 0$, $p_F(\alpha_S) \approx 6 \times 10^{-8}$ for $\rho = 1$ and $p_F(\alpha_S) \approx 8.9 \times 10^{-16}$ for $\rho = -1$.

##### 3.9 Resolvent-based approximation for arbitrary correlation and low variability in self-regulation.

Here, we consider the case in which the interaction matrix is defined as

$$A = \alpha I + D(a) + B,$$

where again $B_{ij}$ $(i \neq j)$ are i.i.d. random variables with zero mean and variance 1, $D(x)$ stands for a diagonal matrix with vector $x$ in the diagonal, and $a = (a_k)$ is a vector of i.i.d. random variables with the same distribution as $B_{ij}$ but zero mean and variance $\sigma_\alpha^2$. As usual, we assume that $a_k$ and $B_{ij}$ are

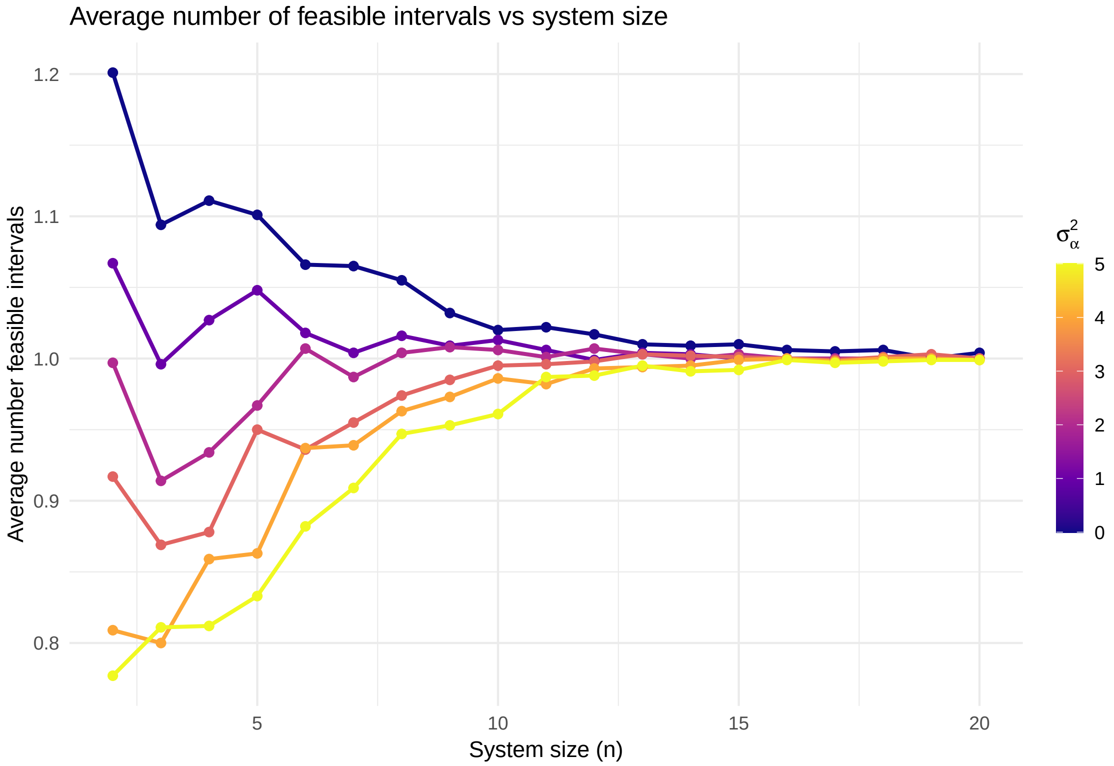


**Supplementary Figure 11**: Introducing heterogeneous self-regulation does not alter the asymptotic convergence to a single transition to feasibility. Average number of feasible intervals as a function of system size ($n$) across different variance levels ($\sigma_\alpha^2$). Each line represents a different diagonal variance parameter ranging from $\sigma_\alpha^2 = 0$ (purple, deterministic diagonal) to $\sigma_\alpha^2 = 5$ (yellow, high variance). Results are averaged over 1000 simulations for each $(n, \sigma_\alpha^2)$ combination. The matrix of interactions is formed by normally distributed off-diagonal elements (without correlation or skewness), with diagonal elements drawn from the specified variance distribution. The number of feasible intervals represents the count of parameter regions where the system maintains feasibility.

independent variables. Because $B_{ii} = 0$, this new definition of interaction coefficients introduces variability in self-regulation. Intra-specific interactions now have mean $\alpha$ and variance $\sigma_\alpha^2$.

As done in the main text, we first show that for large $n$, the system exhibits a single transition to feasibility. In the figure below, we plot the average number of feasible intervals for increasing values of diagonal variance $\sigma_\alpha^2$. We see that increasing such variance only decreases the number of transitions, but the average number of transitions converges to one at approximately the same rate. Therefore, asymptotically, the picture is the same: there is only one transition to feasibility.

The matrix Dyson equation formalism applied to the hermitization of matrix $W = D(a) + B$ does not apply in this case because the entries of that matrix are not i.i.d. variables, given the different variance of the diagonal. However, if $\sigma_\alpha = 1$ all the entries are i.i.d., then the procedure applied above for $\sigma_\alpha = 0$ remains valid and we expect (by continuity of the solution) that it will be valid in a neighborhood of $\sigma_\alpha = 1$. So we will extend that methodology for $\sigma_\alpha \approx 1$ and will check the validity of the approximation as the variance of the self-regulation terms increases.

The hermitization procedure applied in the previous subsection requires to insert, in Eq. (23), the modifications $\mathbb{E}(W^T W) = \mathbb{E}(W W^T) = \mathbb{E}((D(a) + B)^T (D(a) + B)) = \mathbb{E}(D(a)^2) + \mathbb{E}(B^T B) = (\sigma_\alpha^2 + n - 1)I$ and $\mathbb{E}(W^2) = \mathbb{E}((W^T)^2) = \mathbb{E}((D(a) + B)^2) = \mathbb{E}(D(a)^2) + \mathbb{E}(B^2) = (\sigma_\alpha^2 + \rho(n-1))I$, which hold true because

$\mathbb{E}(B) = \mathbb{E}(D(a)) = \mathbb{0}$. Observe that these expectations reduce to the former ones for $\sigma_\alpha = 0$. In order to compute the expectations $\mathbb{E}(x^\star)$ and $\mathbb{E}({x^\star}^T x^\star)$, we have to solve the matrix Dyson equation (21) with a modified definition of the function $\Lambda(\cdot)$, which now reads as

$$\Lambda(S(w,z)) = \begin{pmatrix} iv\mathbb{E}(WW^T) & b\mathbb{E}(W^2) \\ b^\star\mathbb{E}((W^T)^2) & iv\mathbb{E}(W^TW) \end{pmatrix} = \begin{pmatrix} (\sigma_\alpha^2 + n - 1)I & (\sigma_\alpha^2 + \rho(n-1))I \\ (\sigma_\alpha^2 + \rho(n-1))I & (\sigma_\alpha^2 + n - 1)I \end{pmatrix}$$

when $S(w,z)$ is substituted by the asymptotic solution given by Eq. (22). Now we solve Eq. (21) looking for solutions of the same form as above (Eq. (22)). In this case, we have to solve the following quadratic system in $(v,b)$:

$$\begin{aligned} -1 &= -wv - zb + (n-1)(-v^2 + \rho b^2) + \sigma_\alpha^2(-v^2 + b^2), \\ 0 &= wb^\star - zv + (n-1)(vb^\star + \rho vb) + \sigma_\alpha^2(b^\star + b)v, \end{aligned} \tag{32}$$

where, again, the symbol $^\star$ denotes complex conjugate. In fact, we are only interested in obtaining $b(z, w = 0)$. Solving for $v$ in the second equation yields

$$v = -\frac{wb^\star}{(\sigma_\alpha^2 + n - 1)b^\star + (\sigma_\alpha^2 + \rho(n-1))b - z}.$$

Substituting into the first equation and setting $w = 0$ we find that $b(z,0)$ satisfies the quadratic equation $(\sigma_\alpha^2 + \rho(n-1))b^2 - zb + 1 = 0$, which can be solved to obtain

$$b(z,0) = \frac{z - \sqrt{z^2 - 4(\sigma_\alpha^2 + \rho(n-1))}}{2(\sigma_\alpha^2 + \rho(n-1))}, \tag{33}$$

where we have chosen the minus sign so that the resolvent $(zI + B)^{-1}$ behaves as $z^{-1}I$ in the limit $z \to \infty$. Changing $z = \alpha + i\epsilon$ and taking the limit $\epsilon \to 0$ we finally obtain $\mathbb{E}(x^\star) = h(\alpha)\mathbb{1}$, where we have defined

$$h(\alpha) := \frac{\alpha - \sqrt{\alpha^2 - 4(\sigma_\alpha^2 + \rho(n-1))}}{2(\sigma_\alpha^2 + \rho(n-1))} = \frac{2}{\alpha + \sqrt{\alpha^2 - 4(\sigma_\alpha^2 + \rho(n-1))}}. \tag{34}$$

In order to evaluate, in the case of non-zero variance in self-regulation, the expectation $\mathbb{E}({x^\star}^T x^\star)$, we resort to the same argument as before, and use Eq. (28) with $b$ now given by Eq. (33). This gives $\mathbb{E}((\alpha I + D(a) + B^T)^{-1}(\alpha I + D(a) + B)^{-1}) = h_2(\alpha)I$ with

$$h_2(\alpha) := \frac{2}{\alpha^2 + \alpha\sqrt{\alpha^2 - 4(\sigma_\alpha^2 + \rho(n-1))} - 2(1+\rho)(n-1) - 4\sigma_\alpha^2} I. \tag{35}$$

Finally, as in Eq. (30), we can approximate

$$\alpha^2 \sigma_{y^\star}^2 \approx \frac{h_2(\alpha)}{h(\alpha)^2} - 1.$$

This leads to the following approximation for the variance $\sigma_{y^\star}^2$:

$$\frac{1}{(\alpha\sigma_{y^\star})^2} \approx \frac{\alpha^2 + \alpha\sqrt{\alpha^2 - 4(\sigma_\alpha^2 + \rho(n-1))} - 2\sigma_\alpha^2(1-\rho)}{2(\sigma_\alpha^2 + n - 1)} - (1+\rho). \tag{36}$$

We can use this formula to evaluate the probability of feasibility as $p_F(\alpha, \sigma_\alpha^2) = \Phi((\alpha\sigma_{y^\star})^{-1})^n$ when small variability is introduced in self-regulation coefficients ($\sigma_\alpha^2 > 0$). Consistently, this formula reduces to Eq. (31) for $\sigma_a^2 = 0$. Moreover, we expect this formula to hold in a neighborhood of $\sigma_\alpha = 1$, so by continuity this approximation should be accurate for small deviations $\sigma_\alpha \in [0, 1+\delta]$, for $\delta > 0$ small. The overall effect of the variability in self-regulation is to increase the values of average self-regulation ($\alpha$) that allow for feasibility. Does the result $\alpha_F > \alpha_S$ hold (almost surely) in this case?

Universality results in random matrix theory [28] state that for the case studied here ($\sigma_\alpha \approx 1$), the stability threshold also follows the Wigner law (See Methods). As such, since the effect of adding $\sigma_\alpha$ on feasibility, is to move the curve to the right, we expect our results to remain the same (that is, $\alpha_S < \alpha_F$ almost surely). Finally, we take advantage of Eq. (36) to approximate the probability of feasibility as $p_F(\alpha_S, \sigma_\alpha) = \Phi((\alpha_S\sigma_{y^\star})^{-1})^n$ at the threshold of global stability, $\alpha_S$. For instance, we can obtain, for $\sigma_\alpha = 1$ and $n = 50$, such value for $\rho = 0$, $p_F(\alpha_S, \sigma_\alpha) = 1.3 \times 10^{-4}$. Therefore, we find again that $\alpha_F > \alpha_S$ holds almost surely for small variances.

#### 3.10 Computing $\sigma_{y^\star}$ by iteration

In this section, we approximate $\sigma_{y^\star}$ by approximating $y^\star$ for a given $\alpha$. To this end, we use two well-known results.

***Neumann series.***

Whenever the spectral radius of $A$, $\xi(A) < 1$, the inverse of $(I - A)$ can be written as:

$$(I - A)^{-1} = \sum_{k=0}^{\infty} A^k \quad ,$$

where $A^0 = I$.

Take the system in $y^\star$, and divide both sides by $\alpha > 0$:

$$\left(I + \frac{1}{\alpha}C\right) y^\star = \frac{1}{\alpha}\mathbb{1} \quad .$$

Whenever the spectral radius of $C$, $\xi(C) < \alpha$, we can write:

$$\begin{aligned} y^\star &= \frac{1}{\alpha}\left(I + \frac{1}{\alpha}C\right)^{-1}\mathbb{1} \\ &= \frac{1}{\alpha}\sum_{k=0}^{\infty}\left(-\frac{1}{\alpha}C\right)^k\mathbb{1} \\ &= \frac{1}{\alpha}\left(I - \frac{1}{\alpha}C + \frac{1}{\alpha^2}C^2 - \cdots\right)\mathbb{1} \\ &= \frac{1}{\alpha}\mathbb{1} - \frac{1}{\alpha^2}C\mathbb{1} + \frac{1}{\alpha^3}C^2\mathbb{1} - \cdots \quad . \end{aligned}$$

We note that whenever $\alpha > \alpha_L$, the Neumann series converges.

***Richardson iteration.***

The system $Ax = b$ can be solved iteratively, starting from an initial guess $x(0)$ as:

$$x(k+1) = (I - \omega A)x(k) + \omega b \quad ,$$

where $\omega$ is an iteration parameter that can always be chosen such that the iteration converges to the solution of $Ax = b$. When $A = A^T$ and is positive definite, the optimal iteration parameter is simply:

$$\omega = \frac{2}{\lambda_1(A) + \lambda_n(A)} \quad ,$$

i.e., the reciprocal of the average between the smallest and the largest eigenvalue of $A$.

If we multiply the equations we want to solve by $(I + \frac{\kappa}{\alpha} C^T)$, the right-hand side of the equations is unchanged, because $C^T \mathbb{1} = \mathbb{0}$:

$$\left(I + \frac{\kappa}{\alpha} C^T\right)\left(I + \frac{1}{\alpha} C\right) y^\star = \frac{1}{\alpha}\mathbb{1} \quad .$$

However, the left-hand side changes, and thus the speed at which an iterative method would converge to its solution depends on $\kappa$.

Based on numerical simulations, to speed up convergence we choose three different values of $\kappa$, depending on the correlation between the entries of $B$; in particular, we choose $\kappa = 1$ when the correlation is $\rho = -1$ and $B$ is skew-symmetric; $\kappa = 0$ when $\rho = 0$, and $\kappa = -1$ when $\rho = 1$ and $B$ is symmetric.

We start with $\rho = -1$, and multiply both sides of the system of equations by $\left(I + \frac{1}{\alpha} C^T\right)$:

$$\begin{aligned}
\left(I + \frac{1}{\alpha} C^T\right)\left(I + \frac{1}{\alpha} C\right) y^\star &= \frac{1}{\alpha}\left(I + \frac{1}{\alpha} C^T\right)\mathbb{1} \\
\left(I + \frac{1}{\alpha}(C + C^T) + \frac{1}{\alpha^2} C^T C\right) y^\star &= \frac{1}{\alpha}\mathbb{1} \\
\left(I + \frac{1}{\alpha}(B - \mathbb{1}m^T + B^T - m\mathbb{1}^T) + \frac{1}{\alpha^2}(B^T - m\mathbb{1}^T)(B - \mathbb{1}m^T)\right) y^\star &= \frac{1}{\alpha}\mathbb{1} \\
\left(I + \frac{1}{\alpha}(B - \mathbb{1}m^T - B - m\mathbb{1}^T) + \frac{1}{\alpha^2}(-B - m\mathbb{1}^T)(B - \mathbb{1}m^T)\right) y^\star &= \frac{1}{\alpha}\mathbb{1} \\
\left(I + \frac{1}{\alpha}(-\mathbb{1}m^T - m\mathbb{1}^T) + \frac{1}{\alpha^2}(-B^2 - n\, mm^T)\right) y^\star &= \frac{1}{\alpha}\mathbb{1} \\
S y^\star &= \frac{1}{\alpha}\mathbb{1} \quad .
\end{aligned}$$

Note that, a) the multiplication does not alter the right-hand side of the equations; b) $S$ is positive semi-definite, because it is of the form $G^T G$; c) when $\rho = -1$, $B$ is skew-symmetric, and thus $B^T = -B$; d) when $\rho = -1$, necessarily $m^T \mathbb{1} = 0$ (hence the two matrices $\mathbb{1}m^T$ and $m\mathbb{1}^T$ are nilpotent—have all eigenvalues zero).

Next, we choose the iteration parameter, by approximating $S \approx (I - \frac{1}{\alpha^2} B^2)$. Given that the eigenvalues of $B^2$ are the squared eigenvalues of $B$, we have $\lambda_1(B^2) \approx -4n$ and $\lambda_n(B^2) \approx 0$. Hence:

$$\omega = \frac{2}{(1 - \frac{1}{\alpha^2} 0) + (1 + \frac{1}{\alpha^2} 4n)} = \frac{\alpha^2}{\alpha^2 + 2n} \quad .$$

Finally, we take $y(0) = \frac{1}{\alpha}\mathbb{1} - \frac{1}{\alpha^2} C\mathbb{1}$ (as given by the first two terms of the Neumann series), $y(1) = (I - \omega S) y(0) + \frac{1}{\alpha}\omega\mathbb{1}$, and compute the approximate variance as:

$$\mathbb{V}(y_i^\star) = \mathbb{E}((y_i^\star)^2) - \mathbb{E}(y_i^\star)^2 \approx \frac{1}{n}\mathbb{E}(y(1)^T y(1)) - \frac{1}{\alpha^2} \quad .$$

Note that this approximation will depend on the first few moments of the distribution of $B_{ij}$ (and not only on their mean and variance, as in the approximation using small-rank perturbations, or that using resolvents). The expectation of $y(1)^T y(1)$ is presented in Appendix E.

For $\rho = 1$, we multiply both sides of the system of equations by $\left(I - \frac{1}{\alpha}C^T\right)$, instead:

$$\begin{aligned}
\left(I - \frac{1}{\alpha}C^T\right)\left(I + \frac{1}{\alpha}C\right) y^\star &= \frac{1}{\alpha}\left(I - \frac{1}{\alpha}C^T\right)\mathbb{1} \\
\left(I + \frac{1}{\alpha}(C - C^T) - \frac{1}{\alpha^2}C^T C\right) y^\star &= \frac{1}{\alpha}\mathbb{1} \\
\left(I + \frac{1}{\alpha}(B - \mathbb{1}m^T - B^T + m\mathbb{1}^T) - \frac{1}{\alpha^2}(B^T - m\mathbb{1}^T)(B - \mathbb{1}m^T)\right) y^\star &= \frac{1}{\alpha}\mathbb{1} \\
\left(I + \frac{1}{\alpha}(B - \mathbb{1}m^T - B + m\mathbb{1}^T) - \frac{1}{\alpha^2}(B - m\mathbb{1}^T)(B - \mathbb{1}m^T)\right) y^\star &= \frac{1}{\alpha}\mathbb{1} \\
\left(I + \frac{1}{\alpha}(m\mathbb{1}^T - \mathbb{1}m^T) - \frac{1}{\alpha^2}(B^2 - n\,mm^T)\right) y^\star &= \frac{1}{\alpha}\mathbb{1} \\
H y^\star &= \frac{1}{\alpha}\mathbb{1} \quad .
\end{aligned}$$

Note that, a) the multiplication again does not alter the right-hand side of the equations; b) $H$ is positive semi-definite as long as $\alpha > \alpha_L$; c) when $\rho = 1$, $B$ is symmetric, and thus $B^T = B$.

Next, we choose the iteration parameter, by approximating $H \approx (I - \frac{1}{\alpha^2}B^2)$. Given that the eigenvalues of $B^2$ are the squared eigenvalues of $B$, we have $\lambda_n(B^2) \approx 4n$ and $\lambda_1(B^2) \approx 0$. Hence:

$$\omega = \frac{2}{(1 - \frac{1}{\alpha^2}0) + (1 - \frac{1}{\alpha^2}4n)} = \frac{\alpha^2}{\alpha^2 - 2n} \quad .$$

Finally, we take $y(0) = \frac{1}{\alpha}\mathbb{1} - \frac{1}{\alpha^2}C\mathbb{1}$, $y(1) = (I - \omega H)y(0) + \frac{1}{\alpha}\omega\mathbb{1}$, and compute the approximate variance as before. Also for this case, the details of the calculation are presented in Appendix E.

For $\rho = 0$ we choose $\kappa = 0$, leaving the system unchanged. Given that we can safely assume that $\alpha > \alpha_L$, we have that $I - \omega(I + \frac{1}{\alpha}C) = (1 - \omega)I - \frac{\omega}{\alpha}C$ has spectral radius less than one (and thus the iteration converges) when we choose $\omega = 1$, such that the Richardson iteration matches what one would obtain by truncating the Neumann series. Finally, we set $y(0) = \frac{1}{\alpha}\mathbb{1} - \frac{1}{\alpha^2}C\mathbb{1}$ and compute the variance as done for the other cases. The details of the calculation are also presented in Appendix E.

We conclude by stressing that we a) have started with a guess given by the truncation of the Neumann series that includes only the first two terms, and b) have performed only one step of the Richardson iteration. While the calculations are already quite elaborate for these choices, including either more terms of the Neumann series, or performing more than one step for the Richardson iteration (possibly, updating the parameter $\omega$) would almost certainly result in an even more accurate approximation—allowing one to model smaller systems, or systems in which the distribution of coefficients has extreme skewness/kurtosis, etc. Appendix E shows that the approximation above is however quite precise for moderate $n$ and the distributions we have been analyzing in this work.

#### 3.11 Consequences for dynamics

In Sections 3.6 and 3.8, we have shown that, for large $n$, a random competitive system that is feasible is also stable. The competitive GLV system can give rise to all types of dynamics, including limit cycles (for $n \geq 3$) and chaos (for $n \geq 4$) [5, 6]. Non-equilibrium coexistence via limit cycles or chaos requires a *feasible* equilibrium[7], but we have seen in Sec. 1.3 that whenever $\alpha > \alpha_F$, then $\alpha > \alpha_S \geq \alpha_L$, and thus the stability of the equilibrium is ensured for any choice of $r > \mathbb{0}$. Hence, we conclude that limit cycles or chaos are going to be almost impossible to observe for large random competitive systems.

If the relationship $\alpha_F > \alpha_S$ is maintained for moderate sizes $n$ as well (as suggested by the calculations in Sec. 3.8), then the dynamical consequences are even more far-reaching. In a competitive GLV system, lack of feasibility guarantees that one or more populations will go extinct[7]. Suppose that we initialize a random competitive system with $n$ populations at arbitrary initial conditions; this system will have a certain $\alpha$. If $\alpha > \alpha_F$, the system is feasible and stable; trajectories will converge to the equilibrium, and if the equilibrium is perturbed, dynamics will return to it. If, on the other hand, $\alpha < \alpha_F$, then the system is not feasible, and therefore some populations will go extinct. There are two distinct cases:

*Case i):* $\alpha_S < \alpha$: the system will converge to a saturated equilibrium (as shown by the Lyapunov function in Section 1.3); the same populations will go extinct irrespective of the (positive) initial conditions; the populations that go extinct cannot re-invade the system; the populations that coexist do so stably, and when their densities are perturbed, the system will asymptotically return to the saturated equilibrium.

*Case ii):* $\alpha < \alpha_S$: this is the most interesting case. Some populations will go extinct, because there is no feasible equilibrium. However, which populations will go extinct *depends on the initial conditions*. The system can therefore display priority effects and alternative stable states. Every time a population goes extinct, the system size shrinks to $n-1$, $n-2$, etc. Accordingly, the threshold values $\alpha_S$ and $\alpha_F$ are also decreased, while the value of $\alpha$ is unchanged. As such, provided that the $\alpha_S$ for the reduced system is still smaller than the corresponding $\alpha_F$, the system will keep losing populations until $\alpha > \alpha_S$, thus eventually attaining Case i).

Therefore, for large $n$ (and provided that $\alpha_S < \alpha_F$ when populations are lost), the ultimate result is always the same: the surviving populations coexist robustly at a globally-attractive equilibrium. Moreover, while the populations that went extinct at the onset of the dynamics could potentially re-invade the system (only when we start in Case ii), the last few populations (i.e., those that went extinct once $\alpha > \alpha_S$) cannot.

These results, obtained for isolated systems in which populations that are lost cannot reinvade the system are in contrast to those obtained when instead a small but constant influx of each population enters the system at all times; in this case, Bunin[29] showed that for some critical value of $\alpha$ dynamics transition from equilibria to complex, chaotic oscillations. Further analysis shows that the constant influx of immigrants is key to achieve non-equilibrium coexistence, such that when turning off the immigration chaos typically *fades away*[30].

##### 3.12 Consequences for assembly

We can go a step further, and consider whether these systems, obtained by collapsing a larger system, can instead be built from the ground up. As discussed above, we consider the case in which invasions are spaced apart, rather than having a constant influx as in the study by Bunin[29] .

Take a system with $n$ populations coexisting at a feasible equilibrium $x^\star$ (i.e., we have $\alpha > \alpha_F$), possibly obtained by collapsing a larger system, as detailed in the previous section. If we have $\alpha > \alpha_F > \alpha_S$, which is almost invariably the case for a large system, then any feasible sub-community is automatically stable, and might or might not be invaded by the remaining populations.

In such a case, determining whether the equilibrium $x^\star$ can be built by sequential introduction of population can be accomplished by considering an assembly graph where the nodes are the feasible sub-communities that can be formed by selecting some of the $n$ population (out of $2^n$ possibilities, including the "empty" state in which all populations are absent), and an edge connects two nodes if an invasion can move the system from one state to the other[9]. If the system can be built from the ground up, then there exists a path in the graph connecting the empty state all the way up to the state with $n$ populations (Fig. 12).

We performed simulations in which a) we built a feasible system of $n$ populations, b) built the corresponding assembly graph, and c) tallied the probability of finding such a path in the graph as we increased $n$. Figure 13 shows that, even for moderate $n$, the probability is very high, and rapidly converges to one.

Taken collectively, our results show that *robust, stable coexistence* (of a sub-community) *is the norm* for large random competitive communities. For large system with random parameters, dynamics will not display cycles or chaos. Moreover, the final community (i.e., that obtained once the dynamics have elapsed, and extinctions have occurred) is such that priority effects are excluded. The final community can be assembled from the ground-up by introducing one population at a time.

### Supplementary Note 4. Appendices

#### A Distribution of the coefficients $B_{ij}$

Here we detail the distributions we have used to sample interaction coefficients, including their probability distribution/mass functions, and the first six moments (needed for the Richardson approximation). All the distributions are shifted and re-scaled in order to have mean zero and unit variance. We chose a distribution that is discrete and has finite support; a distribution that is continuous and has finite support, and one that is continuous and has infinite support. For each distribution, we include a parameter $\beta$ that can be varied to regulate the skewness of the distributions. The three distributions are drawn in Fig. 14.

#### B Skew normal distribution

The PDF is given by

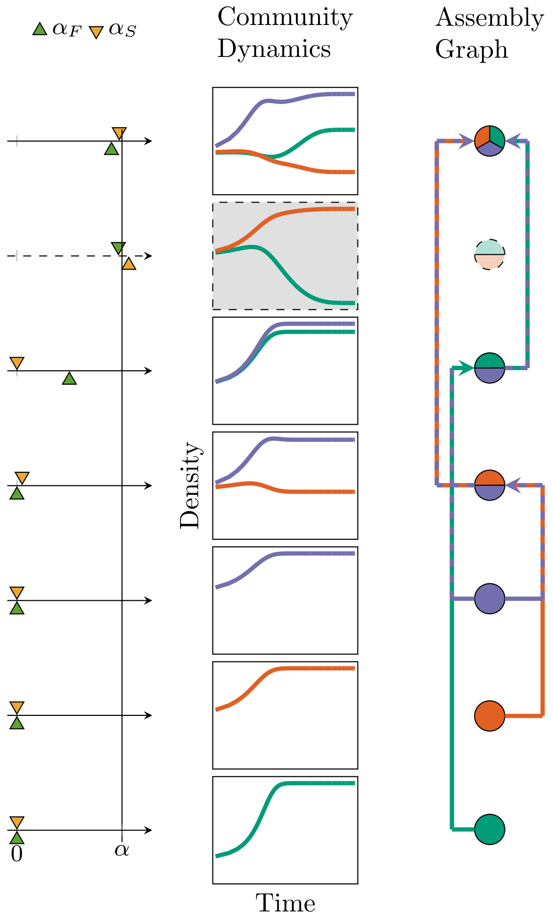


**Supplementary Figure 12**: Assembly graph for a simple competitive community. We show how the interplay between the values of $\alpha$, $\alpha_S$ and $\alpha_F$ (left) determine dynamics (center), and therefore the existence and connectivity of nodes (right) in the assembly graph associated with the system. Starting from the top, we consider a three-species community with $\alpha > \alpha_F > \alpha_S$; thus, the three populations coexist stably (center, showing typical dynamics) and this community is a node in the assembly graph (right). For the three two-species communities that we can form, we have that $\alpha > \alpha_F$ for two of them, while for one community $\alpha_F > \alpha$, and therefore dynamics will lead to extinctions; consequently, the corresponding node is not present in the assembly graph. Finally, by definition, all single-species communities are feasible and stable. We connect two nodes in the assembly graph if an invasion can transition the system from a sub-community (node in the graph) to another. We say that the full community can be built from the ground up via sequential invasions if there is a path starting at a node encoding a single-species community and ending with the full community.

$$\Phi(x) = \frac{2}{\sigma\sqrt{2\pi}} \exp\left(-\frac{x-\mu}{2\sigma^2}\right) \int_{-\infty}^{\beta\left(\frac{x-\mu}{\sigma}\right)} \frac{1}{\sqrt{2\pi}} \exp\left(-\frac{t^2}{2}\right)$$

with expectation

$$\mathbb{E}(x) = \mu + \frac{\sqrt{\frac{2}{\pi}}\beta\sigma}{\sqrt{1+\beta^2}}$$

and variance

$$\mathbb{V}(x) = \sigma^2\left(1 - \frac{2\beta^2}{\pi(1+\beta^2)}\right)$$

We then solve the system of equations $\mathbb{E}(x) = 0$ and $\mathbb{V}(x) = 1$ for $\mu$ and $\sigma$, obtaining

$$\mu^\star = -\frac{\sqrt{2}\beta}{\sqrt{\pi + \beta^2(\pi - 2)}} \qquad \sigma^\star = \frac{\sqrt{\pi(1+\beta^2)}}{\sqrt{\pi + \beta^2(\pi-2)}}$$

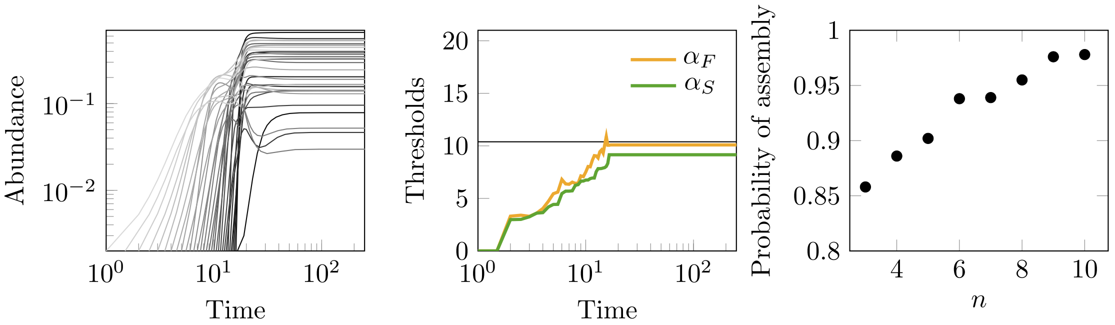


**Supplementary Figure 13**: Bottom-up assembly without extinctions is likely in competitive systems. The first two panels show the assembly of the community in the main text example (Fig. 4), by sequentially invading the system, one population at a time ("bottom-up" assembly). The simulation shows a case in which at each invasion (bottom-right, color corresponding to the time of invasion) the system grows in size, virtually without extinctions, since $\alpha_F$ (almost always) remains below $\alpha$. Additionally, we have $\alpha_S < \alpha_F$ throughout assembly, such the system remains in phase III throughout. As populations are added to the community, both $\alpha_F$ and $\alpha_S$ grow, eventually reaching the values found for top-down assembly. As we vary $n$ (x-axis), we calculate the probability that a random, feasible system can be built from the ground up via successive invasions (y-axis). We performed 1000 simulations for each $n$, constructed the corresponding assembly graph, and, for each graph, we determined whether a path starting from the empty state reaches the node corresponding to the full community. The probability that such a path exists, and therefore the system can be built from the ground up, rapidly converges to one as $n$ increases.

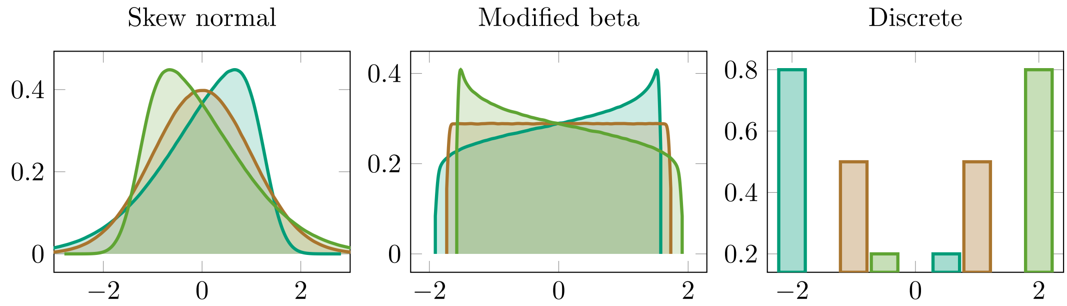


**Supplementary Figure 14**: Distributions from which interaction coefficients are sampled. For our simulations, we sampled the coefficients $B_{ij}$ from distributions with mean zero and unit variance. We choose a continuous distribution with infinite support (Skew normal), a continuous distribution with finite support (Modified beta) and a discrete distribution with finite support (Discrete) to probe the generality of our results. For each distribution, we include a parameter ($\beta$) that controls the skewness (colors).

Finally, we sample interactions from the skew normal distribution with $\mu = \mu^\star$ and $\sigma = \sigma^\star$. The moments of the resulting distribution are given in the table below. Note that for $\beta = 0$ we recover the standard normal distribution.

$$\mu_1 = 0$$

$$\mu_2 = 1$$

$$\mu_3 = -\frac{\sqrt{2}(\pi - 4)\beta^3}{(\pi + (\pi - 2)\beta^2)^{3/2}}$$

$$\mu_4 = \frac{-12\beta^4 + 3\pi^2(1+\beta^2)^2 - 4\pi\beta^2(3+\beta^2)}{(\pi + (\pi-2)\beta^2)^2}$$

$$\mu_5 = \frac{\sqrt{2}\beta^3(-10(\pi-4)\pi + (16 + (20 - 7\pi)\pi)\beta^2)}{(\pi + (\pi-2)\beta^2)^{5/2}}$$

$$\mu_6 = \frac{-40\beta^6 + 15\pi^3(1+\beta^2) - 20\pi\beta^4(9+5\beta^2) - 6\pi^2\beta^2(155 + 10\beta^2 + \beta^4)}{(\pi + (\pi-2)\beta^2)^3}$$

#### C Discrete distribution

The PMF is given by

$$\Phi(x) = \begin{cases} \xi_1 & \text{with probability } p \\ \xi_2 & \text{with probability } 1-p \end{cases}$$

The expectation is

$$\mathbb{E}(x) = \xi_1 p + \xi_2(1-p)$$

and the variance is

$$\mathbb{V}(x) = \xi_1^2 p + \xi_2^2(1-p) - E(x)^2$$

We solve the system of equations imposing that $\mathbb{E}(x) = 0$ and $\mathbb{V}(x) = 1$ for $\xi_2$ and $p$, obtaining

$$p = \frac{1}{1+\xi_1^2} \qquad \beta = -\frac{1}{\xi_1}$$

We call $\beta = \xi_1$; this parameter controls the skewness of the distribution. For $\beta = 1$ we obtain the Bernoulli distribution with outcomes $\pm 1$ and skewness 0. The first six moments for this distribution are

$$\begin{aligned}
\mu_1 &= 0 \\
\mu_2 &= 1 \\
\mu_3 &= \beta - \frac{1}{\beta} \\
\mu_4 &= \beta^2 - 1 + \frac{1}{\beta^2} \\
\mu_5 &= \frac{1}{\beta^3}(\beta^2-1)(\beta^4+1) \\
\mu_6 &= \beta^4 - \beta^2 + 1 - \frac{1}{\beta^2} + \frac{1}{\beta^4}
\end{aligned}$$

#### D Beta distribution

The PDF is given by

$$\Phi(x) = \frac{x^{\xi_1-1}(1-x)^{\xi_2-1}}{\mathtt{B}(\xi_1,\xi_2)}$$

where $\mathtt{B}(\xi_1,\xi_2) = \frac{\Gamma(\xi_1)\Gamma(\xi_2)}{\Gamma(\xi_1+\xi_2)}$, and $\Gamma$ is the Gamma function. The expectation is

$$\mathbb{E}(x) = \frac{\xi_1}{\xi_1+\xi_2}$$

and the variance

$$\mathbb{V}(x) = \frac{\xi_1\xi_2}{(\xi_1+\xi_2)^2(\xi_1+\xi_2+1)}$$

The beta distribution has support between 0 and 1, and therefore it cannot have mean zero and unit variance. For this reason, we perform a change of variables to shift it and re-scale it. That is, we sample $x$ from a beta distribution and then construct another random variable $z$ by subtracting the mean (shift) and dividing by the square root of the variance (re-scale). The result is a random variable following the beta distribution with mean 0 and variance 1. We take $\xi_1 = 1/\beta$, $\xi_2 = \beta$, and define:

$$z = \sqrt{\frac{1+\beta+\beta^2}{\beta^3}}(-1+x+x\beta^2)$$

The first six moments of the distribution defined by the random variable $z$ are

$$\begin{aligned}
\mu_1 &= 0 \\
\mu_2 &= 1 \\
\mu_3 &= \frac{2(\beta-1)\sqrt{1+\frac{1}{\beta}+\beta}}{1+\beta} \\
\mu_4 &= -21+\frac{6}{\beta}+\beta\left(6+\frac{54}{1+\beta(3+\beta)}\right) \\
\mu_5 &= \frac{4(\beta-1)(1+\frac{1}{\beta}+\beta)^{3/2}(6+5\beta+5\beta^3+6\beta^4)}{(1+\beta)(1+\beta(3+\beta))(1+\beta(4+\beta))} \\
\mu_6 &= \frac{5(1+\beta+\beta^2)^2(24+\beta(-22+\beta(16+\beta(-1-22\beta+24\beta^2))))}{\beta^2(1+\beta(3+\beta))(1+\beta(4+\beta))(1+\beta(5+\beta))}
\end{aligned}$$

When $\beta = 1$, we recover the uniform distribution $\mathcal{U}[-\sqrt{3},\sqrt{3}]$, which has mean zero, unit variance, and zero skewness.

### E Calculation of $\sigma_{y^\star}$

Here we present the details of the calculations for the approximation of $\sigma_{y^\star}$.

### F Through iterative solutions of linear systems

We have discussed the approach to approximating the variance $\sigma^2_{y^\star}$ as $\sigma^2_{y^\star} \approx \frac{1}{n}y(1)^T y(1) - \frac{1}{\alpha^2}$, where $y(1)$ is given by the first step of the Richardson iteration:

$$y(1) = y(0) - \omega M y(0) + \omega b$$

We have chosen: a) $y(0)$ as given by the first two terms of the Neumann series, $y(0) = \frac{1}{\alpha}\mathbb{1} - \frac{1}{\alpha^2}C\mathbb{1}$; b) the right-hand side $b = \frac{1}{\alpha}\mathbb{1}$; c) the choice of $\omega$ and $M$ depend on the correlation, as follows:

$$\begin{array}{lll}
\rho = -1 & \rho = 0 & \rho = 1 \\
\omega = \dfrac{\alpha^2}{\alpha^2+2n} & \omega = 1 & \omega = \dfrac{\alpha^2}{\alpha^2-2n} \\
M = \left(I+\dfrac{1}{\alpha}C^T\right)\left(I+\dfrac{1}{\alpha}C\right) & M = \left(I+\dfrac{1}{\alpha}C\right) & M = \left(I-\dfrac{1}{\alpha}C^T\right)\left(I+\dfrac{1}{\alpha}C\right)
\end{array}$$

To compute the approximate variance, we take a generic matrix $B = (B_{ij})$ of size $n$, compute the Richardson iteration, and then the approximate variance:

$$\begin{aligned}\sigma^2_{y^\star} &\approx \frac{1}{n} y(1)^T y(1) - \frac{1}{\alpha^2} \\ &= \frac{1}{n}\left((y(0) - \omega M y(0) + \omega b)^T (y(0) - \omega M y(0) + \omega b)\right) - \frac{1}{\alpha^2} \\ &= \frac{1}{n}\left(y(0)^T y(0) + \omega^2 y(0)^T M^T M y(0) + \omega^2 b^T b - 2\omega y(0)^T M y(0) + 2\omega b^T y(0) - 2\omega^2 b^T M y(0)\right) - \frac{1}{\alpha^2}\end{aligned} \tag{37}$$

Each term of the expansion ($y(0)^T y(0)$, $\omega^2 y(0)^T M^T M y(0)$, etc.) is a polynomial that depends on some of the coefficients $B_{ij}$, as well as $n$, $\alpha$ and $\omega$. To compute the expected variance, we then take the expectation of each of the terms; in particular, we need to determine the expectation of polynomials in $B_{ij}$, such as $B^p_{ij} B^q_{kl}$. When the coefficients are independent (e.g., $\rho = 0$), only the terms in which all the coefficients are raised to a power greater than one will yield a non-zero expectation. In particular, we write:

$$\begin{aligned}\mathbb{E}(B_{ij}) &= 0 \\ \mathbb{E}(B^2_{ij}) &= \mu_2 \\ \mathbb{E}(B^3_{ij}) &= \mu_3 \\ \mathbb{E}(B^4_{ij}) &= \mu_4 \\ &\ldots\end{aligned}$$

Such that for example $\mathbb{E}(B^2_{ij} B^3_{kl}) = \mu_2 \mu_3$. Note that two distributions might have both $\mathbb{E}(B_{ij}) = 0$ and $\mathbb{E}(B^2_{ij}) = 1$, but different higher moments (for example, the standard normal distribution yields $(\mu_2, \mu_3, \mu_4, \mu_5, \mu_6) = (1, 0, 3, 0, 15)$, while when we assign $B_{ij} = 1$ or $B_{ij} = -1$ with equal probability, we obtain $(\mu_2, \mu_3, \mu_4, \mu_5, \mu_6) = (1, 0, 1, 0, 1)$); hence, the approximations will take the first few moments (up to $\mu_6$, for the choice of starting conditions and Richardson iteration) of the distribution of coefficients into account.

When entries are correlated, we have a more difficult task at hand, because for example $\mathbb{E}(B_{ij} B_{ji}) = \rho$, and this relationship needs to be accounted for when taking higher powers (such that $\mathbb{E}(B^2_{ij} B^2_{ji}) \neq 0$). However, the problem is simplified when we consider either $\rho = 1$ or $\rho = -1$, in which cases $\mathbb{E}(B_{ij} B_{ji}) = \mathbb{E}(B^2_{ij}) = \mu_2$ (when $\rho = 1$) or $\mathbb{E}(B_{ij} B_{ji}) = \mathbb{E}(-B^2_{ij}) = -\mu_2$ (when $\rho = -1$), and thus we just need to take the powers of $B_{ij}$ into account. For simplicity, we thus consider only the cases of $\rho = 0$, $\rho = -1$, and $\rho = 1$. The calculations can be extended to the case of a general $\rho$, the cost being to have to keep track of many more combinations of coefficients.

Computing these expectations becomes increasingly complex. We start from the terms that do not depend on the choice of $M$ (and therefore are the same irrespective of the correlation).

The simplest terms in Eq. (37) are $b^T b$:

$$b^T b = \frac{1}{\alpha^2} \mathbb{1}^T \mathbb{1} = \frac{n}{\alpha^2}$$

and

$$b^T y(0) = \frac{n}{\alpha^2}$$

A term that is slightly more complicated to compute is $y(0)^T y(0)$, which will depend on the correlation, but not on the choice of $M$. We report the detailed calculation, because this is the approach we followed to compute all the remaining terms. We can write $y(0)^T y(0)$ as:

$$\begin{aligned}
y(0)^T y(0) &= \left(\left(\frac{1}{\alpha}\mathbb{1}^T - \frac{1}{\alpha^2}\mathbb{1}^T C^T\right)\left(\frac{1}{\alpha}\mathbb{1} - \frac{1}{\alpha^2}C\mathbb{1}\right)\right) \\
&= \mathbb{1}^T\left(\frac{1}{\alpha^2}I + \frac{1}{\alpha^4}C^T C\right)\mathbb{1} \\
&= \frac{n}{\alpha^2} + \frac{1}{\alpha^4}\mathbb{1}^T\left((B^T - m\mathbb{1}^T)(B - \mathbb{1}m^T)\right)\mathbb{1} \\
&= \frac{n}{\alpha^2} + \frac{1}{\alpha^4}\mathbb{1}^T\left(B^T B - n\, mm^T\right)\mathbb{1} \\
&= \frac{n}{\alpha^2} + \frac{1}{\alpha^4}\left(\mathbb{1}^T B^T B\mathbb{1} - n\,\mathbb{1}^T mm^T\mathbb{1}\right) \\
&= \frac{n}{\alpha^2} + \frac{1}{\alpha^4}\left(\mathbb{1}^T B^T B\mathbb{1} - n\,(m^T\mathbb{1})^2\right) \\
&= \frac{n}{\alpha^2} + \frac{1}{\alpha^4}\left(\sum_i\sum_j\sum_k B_{ik}B_{jk} - n\left(\frac{1}{n}\sum_i\sum_j B_{ij}\right)^2\right) \\
&= \frac{n}{\alpha^2} + \frac{1}{\alpha^4}\left(\sum_i\sum_j\sum_k B_{ik}B_{jk} - \frac{1}{n}\left(\sum_i\sum_j B_{ij}\right)^2\right)
\end{aligned}$$

Next, we need to take the expectations for these sums:

$$\mathbb{E}(y(0)^T y(0)) = \frac{n}{\alpha^2} + \frac{1}{\alpha^4}\left(\mathbb{E}\left(\sum_i\sum_j\sum_k B_{ik}B_{jk}\right) - \frac{1}{n}\mathbb{E}\left(\left(\sum_i\sum_j B_{ij}\right)^2\right)\right)$$

These expectations depend on the second moment $\mu_2 = 1$ as well as the correlation $\rho$; in particular, we have:

$$\mathbb{E}\left(\sum_i\sum_j\sum_k B_{ik}B_{jk}\right) = \mathbb{E}\left(\sum_i\sum_k B_{ik}B_{ik}\right) + \mathbb{E}\left(\sum_i\sum_{j\neq i}\sum_k B_{ik}B_{jk}\right) = n(n-1)\mu_2 + 0$$

and, using that $\mu_2 = 1$:

$$\begin{aligned}
E\left(\left(\sum_i\sum_j B_{ij}\right)^2\right) &= E\left(\sum_i\sum_j B_{ij}^2\right) + E\left(\sum_i\sum_j\sum_{k\neq i}\sum_{l\neq j} B_{ij}B_{kl}\right) \\
&= n(n-1)\mu_2 + n(n-1)\rho = n(n-1)(1+\rho)\mu_2
\end{aligned}$$

Thus yielding:

$$\mathbb{E}(y(0)^T y(0)) = \frac{n}{\alpha^2} + \frac{1}{\alpha^4}\mu_2(n-1)(n-1-\rho).$$

***Independent entries,*** $\rho = 0$

For the remaining four terms in Eq. (37), we need to perform calculations that differ for $\rho = 0$, $\rho = -1$ and $\rho = 0$, because in each case we have a different matrix $M$. We start with the simplest case, which is obtained when the coefficients of $B$ are sampled independently, and $\rho = 0$. We have:

$$
\begin{aligned}
M &= I + \frac{1}{\alpha} C \\
\mathbb{E}(b^T b) &= \frac{n}{\alpha^2} \\
\mathbb{E}(b^T y(0)) &= \frac{n}{\alpha^2} \\
\mathbb{E}(y(0)^T y(0)) &= \frac{n}{\alpha^2} + \frac{1}{\alpha^4}(n-1)^2 \mu_2
\end{aligned}
$$

We need to compute the remaining terms. $b^T M y(0)$ is very easy to compute:

$$
b^T M y(0) = b^T y(0) = \frac{n}{\alpha^2}
$$

Computing $y(0)^T M y(0)$ is more involved, but can be accomplished using the same approach used for $y(0)^T y(0)$, and noting that only the symmetric part of $M$ matters. We obtain:

$$
\mathbb{E}(y(0)^T M y(0)) = \frac{n}{\alpha^2} - \frac{(n-1)^2}{n\alpha^5} \mu_3
$$

Finally, the most complicated term, $y(0)^T M^T y(0)$, for which we obtain:

$$
\mathbb{E}(y(0)^T M^T M y(0)) = \frac{n}{\alpha^2} + \frac{(n-1)(n^4 - 3n^3 + 4n^2 - 6n + 3)}{n^2 \alpha^6} \mu_2^2 + \frac{(n-1)^2}{n^2 \alpha^6} \mu_4
$$

Combining the terms according to Eq. (37), we find that the expected variance for $\rho = 0$ is:

$$
\mathbb{E}(\sigma^2_{y^\star}) \approx \frac{\mu_2 (n-1)^2}{\alpha^4 n} + \frac{2\mu_3 (n-1)^2}{\alpha^5 n^2} + \frac{\mu_4 (n-1)^2}{\alpha^6 n^3} + \frac{\mu_2^2 \left(n^4 - 3n^3 + 4n^2 - 6n + 3\right)(n-1)}{\alpha^6 n^3}
$$

A few notable features of this equation are: a) the approximation depends on the first few moments of the distribution, and not only on the mean and variance of $B_{ij}$; b) the moment $\mu_3$, summarizing the skewness of the distribution, is multiplied by a positive coefficient, hence a positive skew will increase the variance (and hence decrease the probability of feasibility, compared to a negative skew); c) however, the terms associated with the higher moments vanish when $n$ is large and $\alpha > \sqrt{n}$, again confirming that for sufficiently large $n$ the behavior will be universal.

***Symmetric entries,*** $\rho = 1$

We repeat the calculation for the case of symmetric matrices $B$, for which we choose a different matrix $M$ and a different parameter $\omega$, in order to speed up the convergence. We have:

$$
\begin{aligned}
M &= \left(I - \frac{1}{\alpha}C^T\right)\left(I + \frac{1}{\alpha}C\right) \\
\mathbb{E}(b^T b) &= \frac{n}{\alpha^2} \\
\mathbb{E}(b^T y(0)) &= \frac{n}{\alpha^2} \\
\mathbb{E}(y(0)^T y(0)) &= \frac{n}{\alpha^2} + \frac{1}{\alpha^4}(n-1)(n-2)\mu_2
\end{aligned}
$$

We need to compute the remaining terms:

$$
b^T M y(0) = \frac{n}{\alpha^2} + \mu_3 \frac{(n-1)(n-2)^2}{n\alpha^5}
$$

$$
\begin{aligned}
\mathbb{E}(y(0)^T M y(0)) &= \frac{n}{\alpha^2} - \frac{(n-1)(n-2)(n^3-5n^2+18n-12)}{n^2\alpha^6}\mu_2^2 \\
&\quad + \frac{2(n-1)(n-2)^2}{n\alpha^5}\mu_3 - \frac{(n-1)(n-2)^3}{n^2\alpha^6}\mu_4 \\
\mathbb{E}(y(0)^T M^T M y(0)) &= \frac{n}{\alpha^2} + \frac{(n-1)(n-2)(n^3-5n^2+18n-12)}{n^2\alpha^6}\mu_2^2 \\
&\quad + \frac{(n-1)(n-2)(n^5-17n^4+105n^3-392n^2+724n-432)}{n^3\alpha^8}\mu_2^3 \\
&\quad - \frac{4(n-1)(n-2)(n^4-12n^3+49n^2-88n+40)}{n^3\alpha^7}\mu_3\mu_2 \\
&\quad + \frac{2(n-1)(n-2)(n^4-14n^3+56n^2-92n+64)}{n^3\alpha^8}\mu_3^2 \\
&\quad + \frac{2(n-1)(n-2)(2n^4-17n^3+76n^2-140n+92)}{n^3\alpha^8}\mu_4\mu_2 \\
&\quad + \frac{2(n-1)(n-2)^2}{n\alpha^5}\mu_3 - \frac{2(n-1)(n-2)^4}{n^3\alpha^7}\mu_5 + \frac{(n-1)(n-2)^4}{n^3\alpha^7}\mu_6
\end{aligned}
$$

Combining all terms, we find that the expected variance for $\rho = 1$ is:

$$
\begin{aligned}
\mathbb{E}(\sigma_{y^\star}^2) \approx{} & \frac{\mu_2(n-1)(n-2)}{\alpha^4 n} - \frac{4\mu_3(n-1)(n-2)^2\omega}{\alpha^5 n^2} \\
& - \frac{4\mu_2\mu_3(n-1)\left(n^4-12n^3+49n^2-88n+40\right)(n-2)\omega^2}{\alpha^7 n^4} \\
& + \frac{\mu_6(n-1)(n-2)^4\omega^2}{\alpha^8 n^4} - \frac{2\mu_5(n-1)(n-2)^4\omega^2}{\alpha^7 n^4} \\
& + \frac{\mu_2^2(n-2)(n-1)\left(n^3-5n^2+18n-12\right)\omega^2}{\alpha^6 n^3} \\
& + \frac{2\mu_2^2(n-2)(n-1)\left(n^3-5n^2+18n-12\right)\omega}{\alpha^6 n^3} \\
& + \frac{2\mu_3^2(n-1)\left(n^4-14n^3+56n^2-92n+64\right)(n-2)\omega^2}{\alpha^8 n^4} \\
& + \mu_4\left(\frac{(n-1)(n-2)^3\omega^2}{\alpha^6 n^3} + \frac{2(n-1)(n-2)^3\omega}{\alpha^6 n^3}\right) \\
& + \frac{2\mu_2\mu_4(n-1)\left(2n^4-17n^3+76n^2-140n+92\right)(n-2)\omega^2}{\alpha^8 n^4} \\
& + \frac{\mu_2^3(n-1)\left(2n^5-17n^4+105n^3-392n^2+724n-432\right)(n-2)\omega^2}{\alpha^8 n^4}
\end{aligned}
$$

***Skew-symmetric entries,*** $\rho = -1$

Finally, we compute the expectation for all the terms when the matrix $B$ is skew-symmetric, again choosing a different matrix $M$ and a different parameter $\omega$, in order to speed up the convergence. We have:

$$\begin{aligned} M &= \left(I + \frac{1}{\alpha}C^T\right)\left(I + \frac{1}{\alpha}C\right) \\ \mathbb{E}(b^T b) &= \frac{n}{\alpha^2} \\ \mathbb{E}(b^T y(0)) &= \frac{n}{\alpha^2} \\ \mathbb{E}(y(0)^T y(0)) &= \frac{n}{\alpha^2} + \frac{1}{\alpha^4}(n-1)n\mu 2 \end{aligned}$$

We find:

$$b^T M y(0) = \frac{n}{\alpha^2}$$

$$\mathbb{E}(y(0)^T M y(0)) = \frac{n}{\alpha^2} + \frac{(n-1)(n-2)(n-3)}{\alpha^6}\mu_2^2 - \frac{(n-1)(n-2)}{\alpha^6}\mu_4$$

$$\begin{aligned} \mathbb{E}(y(0)^T M^T M y(0)) = & \frac{n}{\alpha^2}\frac{(n-1)(n-2)(n-3)}{\alpha^6}\mu_2^2 \\ & + \frac{(n-1)(n-2)(n^3-15n^2+45n-60)}{n\alpha^8}\mu_2^3 \\ & + \frac{2(n-1)(n-2)(n^2-8n+14)}{3n\alpha^8}\mu_3^2 \\ & + \frac{(n-1)(n-2)}{\alpha^6}\mu_4 \\ & + \frac{2(n-1)(n-2)(2n^2-10n+15)}{n\alpha^8}\mu_4\mu_2 \\ & + \frac{(n-1)(n-2)^2}{n\alpha^8}\mu_6 \end{aligned}$$

Putting all these terms together, we find that the expected variance for $\rho = -1$ is:

$$\begin{aligned} \mathbb{E}(\sigma_{y^\star}^2) \approx & \frac{2\mu_3^2(n-2)(n-1)((n-8)n+14)\omega^2}{3\alpha^8 n^2} \\ & + \frac{\mu_6(n-2)^2(n-1)\omega^2}{\alpha^8 n^2} + \frac{\mu_4(n-2)(n-1)\omega\left(30\mu_2\omega + 4\mu_2 n^2\omega + \alpha^2 n(\omega-2)\right)}{\alpha^8 n^2} \\ & + \frac{\mu_2(n-1)\left(\alpha^4 n^2 + \mu_2(n-2)\omega^2\left(\alpha^2(n-3)n + \mu_2(n(n(2n-15)+45)-60)\right)\right)}{\alpha^8 n^2} \\ & - \frac{2\mu_2^2\alpha^2(n-3)(n-2)(n-1)\omega}{\alpha^8 n} - \frac{20\mu_2\mu_4(n-2)(n-1)\omega^2}{\alpha^8 n} \end{aligned}$$

where $\omega = \frac{\alpha^2}{\alpha^2+2n}$.

#### G Using resolvents

***Dyson equation derivation.***

In this section we present the derivation of the Dyson equation (9) for symmetric random matrices. We start from the resolvent of matrix $B$,

$$H(z) = (zI + B)^{-1},$$

i.e., the inverse of matrix $zI + B$. The Neumann series expansion for $H(z)$ is convergent for any $z \in \mathbb{C}$ such that $\mathcal{I}m(z) > 0$. Recall that the expectation of the resolvent is denoted as $G(z) = \mathbb{E}(H(z))$. The equation above can be rewritten as

$$zH(z) - I = -BH(z).$$

We subtract, from both sides, the matrix product $\Sigma H(z)$, with $\Sigma = \mathbb{E}(B\mathbb{E}(H(z))B) = \mathbb{E}(BG(z)B)$. Thus, we get

$$-I + (zI - \Sigma)H(z) = -BH(z) - \Sigma H(z).$$

Taking expectations on both sides of this equation, we obtain

$$\mathbb{E}(-I + (zI - \Sigma)H(z)) = -\mathbb{E}(BH(z) + \Sigma H(z)).$$

Because $\Sigma$ is an expectation, this expression reduces to

$$-I + (zI - \Sigma)G(z) = -\mathbb{E}(BH(z) + \Sigma H(z)).$$

It can be shown in general that the right-hand side of this equation is negligible up to leading order, as $n \to \infty$, for arbitrary distributions [25]. Hence the resolvent satisfies the matrix equation

$$(zI - \mathbb{E}(BG(z)B))G(z) = I.$$

After elementary manipulations, this equation can be rewritten in the usual form, see Eq. (9).

***Proof of Eq.* (11).**

Let $K(z) = \mathbb{E}((zI+B)^{-2})$. Here we show that $K(z) = -G'(z)$, where $G(z)$ is the expectation of the resolvent, $G(z) = \mathbb{E}((zI + B)^{-1})$, and the prime stands for the derivative with respect to $z$.

To compute the derivative, let us introduce a real parameter $q \neq 0$ that we can use to take derivatives with respect to it. We define $H_q(z) = (zI + qB)^{-1}$ and $L_q(z) = (zI + qB)^{-2}$, so $H_1(z) = H(z)$, with $H(z) = (zI + B)^{-1}$, and $L_1(z) = L(z)$, with $L(z) = (zI + B)^{-2}$. Observe that $\mathbb{E}(H(z)) = G(z)$ and $\mathbb{E}(L(z)) = K(z)$. It holds that

$$H_q(z) = \left(q\left(\frac{z}{q}I + B\right)\right)^{-1} = \frac{H(z/q)}{q}.$$

On the other hand, because $\mathscr{I}m(z) > 0$, the Neumann series for $L_q(z)$ converges. We compute it using the Neumann series for $H_q(z)$:

$$H_q(z) = \left(z\left(I + \frac{q}{z}B\right)\right)^{-1} = \frac{1}{z}\sum_{k=0}^{\infty}\left(-\frac{q}{z}\right)^k B^k.$$

Hence

$$L_q(z) = \frac{1}{z^2}\sum_{k=0}^{\infty}\left(-\frac{q}{z}\right)^k B^k \sum_{l=0}^{\infty}\left(-\frac{q}{z}\right)^l B^l.$$

We write the product of the two series using the Cauchy product, i.e.,

$$L_q(z) = \frac{1}{z^2}\sum_{s=0}^{\infty} c_s(z,q),$$

with $c_s(z,q)$ defined as

$$c_s(z,q) = \sum_{j=0}^{s}\left(-\frac{q}{z}\right)^j B^j \left(-\frac{q}{z}\right)^{s-j} B^{s-j} = (s+1)\left(-\frac{q}{z}\right)^s B^s.$$

Therefore

$$L_q(z) = \frac{1}{z^2}\sum_{s=0}^{\infty}(s+1)\left(-\frac{q}{z}\right)^s B^s = \frac{1}{z}H_q(z) + \frac{1}{z^2}\sum_{s=1}^{\infty} s\left(-\frac{q}{z}\right)^s B^s.$$

The second term in the last equation can be written in terms of a derivative of $H_q(z)$ with respect to $q$:

$$\begin{aligned}\frac{\partial}{\partial q}H_q(z) &= \frac{1}{z}\frac{\partial}{\partial q}\sum_{k=0}^{\infty}\left(-\frac{q}{z}\right)^k B^k = -\frac{1}{z^2}\sum_{k=1}^{\infty} k\left(-\frac{q}{z}\right)^{k-1} B^k \\ &= -\frac{1}{z^2}\left(-\frac{z}{q}\right)\sum_{k=1}^{\infty} k\left(-\frac{q}{z}\right)^k B^k = \frac{1}{qz}\sum_{k=1}^{\infty} k\left(-\frac{q}{z}\right)^k B^k\end{aligned}$$

Therefore,

$$\sum_{s=1}^{\infty} s\left(-\frac{q}{z}\right)^s B^s = qz\frac{\partial}{\partial q}H_q(z),$$

and it holds that

$$L_q(z) = \frac{1}{z}H_q(z) + \frac{q}{z}\frac{\partial}{\partial q}H_q(z).$$

Now we use that $H_q(z) = H(z/q)/q$ to compute the partial derivative:

$$\frac{\partial}{\partial q}H_q(z) = -\frac{1}{q^2}H(z/q) - \frac{z}{q^3}H'(z/q).$$

Upon substitution of this result into the equation for $L_q(z)$, and using the expression of $H_q(z)$ in terms of $H(z)$, we find

$$L_q(z) = \frac{1}{qz}H(z/q) + \frac{q}{z}\left(-\frac{1}{q^2}H(z/q) - \frac{z}{q^3}H'(z/q)\right) = -\frac{1}{q^2}H'(z/q).$$

Making the substitution $q = 1$ we get $L(z) = -H'(z)$.

To conclude the proof, we take expectations on both sides of the equation,

$$K(z) = \mathbb{E}(L(z)) = -\mathbb{E}(H'(z)).$$

By differentiating the Neumann series, we obtain

$$H'(z) = -\frac{1}{z^2}\sum_{k=0}^{\infty}(k+1)\left(-\frac{1}{z}\right)^k B^k,$$

and taking expectations

$$\mathbb{E}(H'(z)) = -\frac{1}{z^2}\sum_{k=0}^{\infty}(k+1)\left(-\frac{1}{z}\right)^k \mathbb{E}(B^k).$$

On the other hand, by definition

$$G(z) = \mathbb{E}((zI + B)^{-1}) = \frac{1}{z}\sum_{k=0}^{\infty}\left(-\frac{1}{z}\right)^k \mathbb{E}(B^k).$$

By taking the derivative of $G(z)$ with respect to $z$ we finally obtain

$$G'(z) = -\frac{1}{z^2}\sum_{k=0}^{\infty}(k+1)\left(-\frac{1}{z}\right)^k \mathbb{E}(B^k) = \mathbb{E}(H'(z))$$

and the result $K(z) = -G'(z)$ is proved.
